# Two-way pantothenate/nitrogen exchange reprograms fly metabolism and reinforces microbiota growth

**DOI:** 10.1101/2025.11.15.688587

**Authors:** David R Sannino, Jonathan RH Booth, Jonathan Josephs-Spaulding, Gavin Blackburn, Andrea Andress Huacachino, Alina Malita, Diana Marcu, D. Grace Ross, Konstantinos Gerasimidis, Peter D Newell, Kim Rewitz, Nathaniel W Snyder, Adam J Dobson

## Abstract

Gut microbiota provide nutrients to their animal hosts, but if and how hosts reciprocate with nutrients for microbes is unclear. We show that *Drosophila* and their gut bacteria trade a vitamin for nitrogen sources. *Acetobacter pomorum* supply pantothenate (vitamin B_5_), which nearly doubles downstream fly coenzyme A. This is accompanied by reprogrammed fly carbohydrate metabolism. Bacteria reduce host lipid storage, dependent on both bacterial pantothenate synthesis and fly pantothenate metabolism. In exchange, bacteria promote host excretion of compounds that rescue growth of nitrogen stressed bacteria, dependent on fly pantothenate metabolism and bacterial purine metabolism. Impairing fly pantothenate metabolism reduced the *in vivo* bacterial population ten-fold. These results indicate a *quid pro quo* metabolic trade in a facultative symbiosis.

## Introduction

Gut bacteria have well-established roles in animal physiology^1^, affecting diverse processes including nutrition, pathogen defence^2–5^, detoxification ^6^, immune entrainment ^7^, behaviour ^8,9^, cognition ^10,11^, lifespan ^12–14^, mate-preference ^15^ and reproduction ^16^. These impacts are widespread among diverse hosts, suggesting evolutionary conservation. The most parsimonious explanation for conservation of phenotype is that conserved mechanisms are involved, yet we lack a unified mechanistic framework across the diversity of hosts. Metabolic interactions are strong candidate mechanisms ^17–21^, because many metabolic pathways are conserved between eukaryotes and bacteria, and so bacterial products can feed into host metabolism. Microbial provision of essential nutrients (i.e. that hosts cannot synthesise themselves) is expected to play significant roles ^22–26^, and microbes may remove nutrients from ingesta before host uptake ^27,28^. In addition to nutritional effects, microbiota may more profoundly reprogram host metabolism, reshaping how the host metabolises nutrients^29^. Reprogramming would indicate profound microbial influence on host biology, but evidence so far remains scant.

Comparatively little is known about what hosts provide to their microbiota. Honeybees can provide organic acids ^30^, and hibernating ground squirrels can provide urea^31^. Overall, whether hosts and microbes provide reciprocally one another with nutrients and metabolites is unclear. Addressing this knowledge gap is critical to understanding how symbioses are maintained and managed, by both partners.

Here, we characterise host-microbe metabolite exchange and host metabolic reprogramming. We use *Drosophila melanogaster*, as a simple, reductionist system in which fundamental biology is tractable. Flies can be reared axenic (i.e. germ-free) from the egg, and their simple microbiota, dominated by Acetobacteraceae and Lactobacillaceae ^32^, can be cultured and reintroduced, generating gnotobiotic flies ^33^. Using genome-sequenced strains elucidates biosynthetic capacity, among which causal mechanisms can be established with bacterial mutants. A fully-defined (holidic) diet ^34^ gives absolute control over chemical inputs to the system ^26,35^. We use this model and metabolomics to elucidate impacts of specific bacteria, consequences for host metabolomes, and reciprocal host-microbe feedbacks

## Results

### SCFAs are dispensable for *A. pomorum/L. brevis* impacts on fly biology

We first confirmed previous findings that axenic flies (females) accumulate triacylglyceride (TAG) relative to conventionally-reared (CR) flies, and that this can be rescued by *Acetobacter* species (*A. pomorum, A. tropicalis, A. fabarum)* but not Lactobacillaceae (*L. plantarum*, *L. brevis*) (Figure 1A) ^33,36^. TAG can therefore be used as a marker of microbiota-dependent fly metabolism.

**Figure 1.**
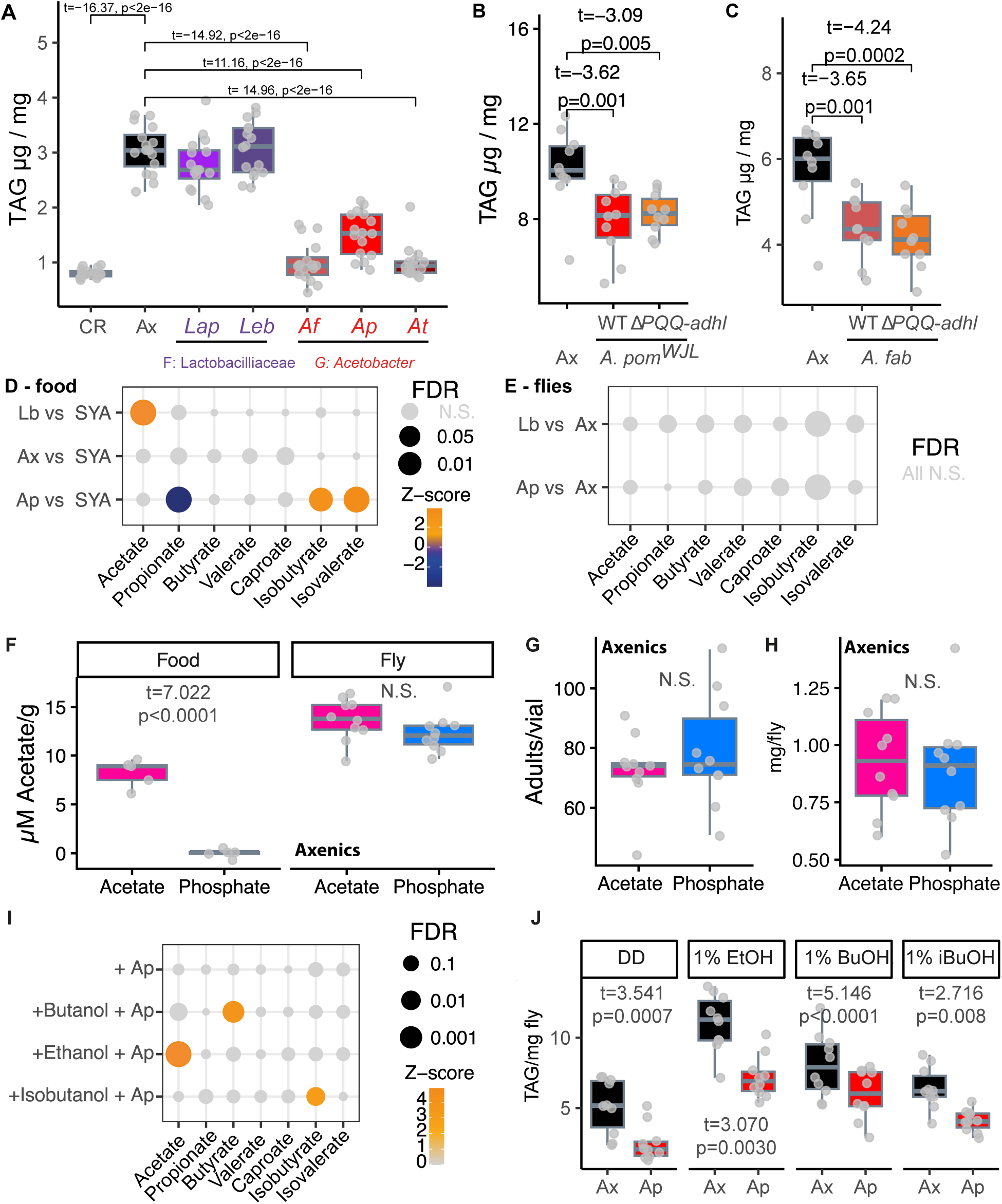
SCFAs are dispensable for *A. pomorum/L. brevis* impacts on fly biology. **A.** Acetobacteraceae rescue host TAG, while Lactobacillaceae do not. Conventionally-reared (CR) and gnotobiotic flies monoassociated with genus *Acetobacter* (*Af - Acetobacter fabarum, Ap – A. pomorum,* and *At – A. tropicalis*) maintain similar TAG levels, while axenics (Ax) and gnotobiotic flies monoassociated with family Lactobacillaceae spp. (*Lap* – *Lactiplantibacillus plantarum* and *Leb* – *Levilactobacillus brevis*) have increased TAG. ANOVA F_6,105_=17.12, p<2.2e-16. **B.** TAG levels of 3-day old females monoassociated with *A. pomorum^WJL^* and an acetate mutant *A. pomorum^WJL^* Δ*PQQ-adhI* are reduced compared to Ax flies. ANOVA F_2,27_=7.64, p=0.002. **C.** TAG levels of 3-day old females monoassociated with wild-type *A. fabarum* and an acetate mutant (Δ*PQQ-adhI*) are reduced compared to Ax flies. ANOVA F_2,27_=10.56, p=0.0004. Statistics on A-C generated by comparing levels of microbiota with axenic set as intercept. **D.** Distinct SCFA production profiles by Ax-flies, Lb-flies and *Ap*-flies. Bubble plot shows change in SCFAs deposited in media (columns) for each condition (rows), relative to unconditioned SYA media. Grey indicates no significant difference. Colours indicate direction and magnitude of change, when significant differences (FDR≤0.05) detected. Bubble size is inverse to FDR (i.e. bigger bubbles indicate more statistically significant difference). Raw data presented in Figure S1G. **E.** SCFA levels in the fly body are insulated from changes in microbial SCFA production. (Lack of) changes in SCFA levels induced by *A. pomorum* (Ap) or *L. brevis* (Lb) are shown, in flies under the same conditions as panel D. Raw data presented in Figure S1G. **F.** Flies reared without microbiota or dietary acetate still accumulate acetate. Boxplots show acetate levels in sterile defined media (food) buffered with either acetate or phosphate, and in corresponding axenic flies (3-day-old adult females) reared on the acetate/phosphate-buffered media from egg to adult. Statistics from post-hoc tests (emmeans) of linear model. **G.** Rearing axenic flies with/without dietary acetate does not impact developmental viability. Wilcoxon rank-sum test p=0.57. **H.** Rearing axenic flies with/without dietary acetate does not impact body mass. T-test p=0.72. **I.** *A. pomorum* SCFA production can be obviated on *Drosophila* defined diet media by omitting alcohol substrates, or specific SCFA production can be induced by adding specific alcohols. Bubble plot shows increase in SCFA levels (Z-scores) relative to unconditioned medium, coloured grey when no significant change detected (FDR≤0.05). SCFA levels were assayed 48h after inoculation. Bubble size increases with statistical significance as per panel D. Raw data presented in Figure S2A. **J.** *A. pomorum* rescues host TAG independent of capacity for SCFA production. Phosphate-buffered defined diets (DD) containing alcohols were generated. *A. pomorum* were inoculated for 48h as per panel I. These diets were fed to female axenic flies reared from egg to adult (on SYA medium, 72h post-eclosion) for seven days, followed by TAG assay. ANOVA: microbiota:diet F_3,72_=1.15, p=0.33; microbiota F_1,72_=52.38, p=4.03e-10; diet F_3,72_=37.10, p=1.32e-14. Pairwise comparisons on figure from post-hoc tests (EMMeans).

Previously, on an imbalanced diet, *A. pomorum*’s TAG-lowering effect was attributed to PQQ-AdhI, which oxidises ethanol to acetate ^17^. Acetate is but one of numerous SCFAs that bacteria can synthesise, e.g. mammalian microbiota also produce propionate and butyrate prolifically ^37^. We were therefore interested in the mechanistic impact of SCFAs on flies. We first tested whether the role of *A. pomorum* PQQ-AdhI on imbalanced diet replicated on a healthy diet, but we found that *PQQ-adhI* mutants rescued TAG equivalently to wild-types, in both *A. pomorum* (Figure 1B) and *A. fabarum* (Figure 1C), indicating an acetate-independent mechanism. Furthermore, both wild-types were net acetate users *in vitro* (Figure S1A-C), suggesting that acetate provision is not constitutive. We therefore assessed whether overall bacterial SCFA production was physiologically relevant, using *A. pomorum* for consistency with prior work. Since the bacteria inhabit both fly gut and medium, microbial impacts on gut nutrient content and availability to the host should be reflected in media composition. By GC-FID, we quantified SCFA output into spent medium from *A. pomorum*-monoassociated flies (“*Ap*-flies”), with CR flies as a positive control and axenics/unconditioned medium as negative controls (Figure S1D). Axenics did not increase media SCFA levels, confirming that the fly alone is not a net SCFA producer. CR flies increased media acetate levels but, unexpectedly, *Ap-*flies did not, suggesting that other bacteria in the CR microbiota were responsible for acetate production. Genomic analysis indicated that *A. pomorum* produce extracellular acetate facultatively from ethanol (Figure S1E), but Lactobacillaceae, such as *L. brevis*, produce acetate following glycolysis or through the oxygen-dependent conversion of lactate to acetate (Figure S1F) ^38^. Accordingly, *L. brevis* increased acetate in spent media >10-fold (Figure 1D, S1G), but this increase did not correlate with altered TAG (Figure 1A), suggesting that acetate provision alone is not sufficient to reduce fly energy storage.

Looking at the greater diversity of SCFAs, we asked whether bacterial modulation correlated with fly SCFA levels. Paired measurements in spent media (Figure 1D) and flies (Figure 1E) showed that flies were buffered against bacterial SCFA supply, suggesting microbiota-independent homeostasis. Together, these data suggested that fly SCFA levels and TAG were independent of microbial SCFA production.

Given prior interest specifically in acetate, we tested whether fly metabolism can generate an acetate pool independent of microbial/dietary supply (as in mammals ^39^), by rearing axenics on acetate-free defined diets. Acetate levels in whole axenics were unaffected by presence/absence of dietary acetate (Figure 1F), and neither viability (Figure 1G) nor body weight (Figure 1H) were compromised, indicating that flies are self-sufficient and do not require bacterial or dietary acetate supply.

As a final test, we fed flies SCFA-free media, on which bacterial SCFA production was selectively induced by adding specific alcohol substrates. Without substrates, *A. pomorum* produced no measurable SCFAs; while adding ethanol, butanol, or isobutanol induced acetate, butyrate, or isobutyrate production, respectively (Figure 1I, Figure S2). Feeding these media to axenic adults for one week (guided by pilot data, Figure S3), showed that *A. pomorum* reduced TAG independent of the presence/absence of SCFA-production substrates (Figure 1J). Altogether, our results indicated microbial effects on the host independent of SCFA production, implicating other, yet-undefined mechanisms.

### Fly metabolomic state corresponds to microbiota-dependent nutrition

To outline novel mechanism(s), we performed LC-MS metabolomics on axenic flies, *Ap-*flies, and *L. brevis-*monoassociated flies (*Lb-*flies). First, we tested (A) assay sensitivity for metabolic responses to microbes, and (B) sexual dimorphism in responses. Metabolic responses to microbes were detectable, and sexual dimorphism in response was negligible, leading us to use adult females in all subsequent experiments (Supplemental note 1).

To assess microbe-dependent nutrition and corollary impacts on the host we performed LC-MS on flies with paired chemical characterisation of their spent (developmental) media, alongside baseline unconditioned (SYA) medium (Figure 2A). Fly and media measurements were paired per container to enable correlative analysis. Principal Components Analysis (PCA) separated fly vs. media on PC1, and bacterial effects on PC2 (Figure S7A), confirming detection of both axes of variation. All three classes of flies increased media chemical complexity (i.e. n. detectable compounds) relative to unconditioned medium, most markedly in *Ap-*flies’ media (Figure S7B-C). Chemical complexity of axenics was less than *Ap-*flies or *Lb-*flies, though changes were more subtle than in media (Figure S7B-C). In media, differential abundance analysis showed that axenics increased more compounds than they downregulated, suggesting they excrete a greater diversity of compounds than they consume (Figure S8A), but relative to this baseline both *Ap-* and *Lb-*flies reduced levels of more compounds than they increased (Figure S8A), suggesting that both symbioses drive a switch toward use of greater chemical diversity. By contrast, in the fly more differential metabolites were increased than decreased, (Figure S8A), suggesting that both bacteria promote nutrition overall.

**Figure 2.**
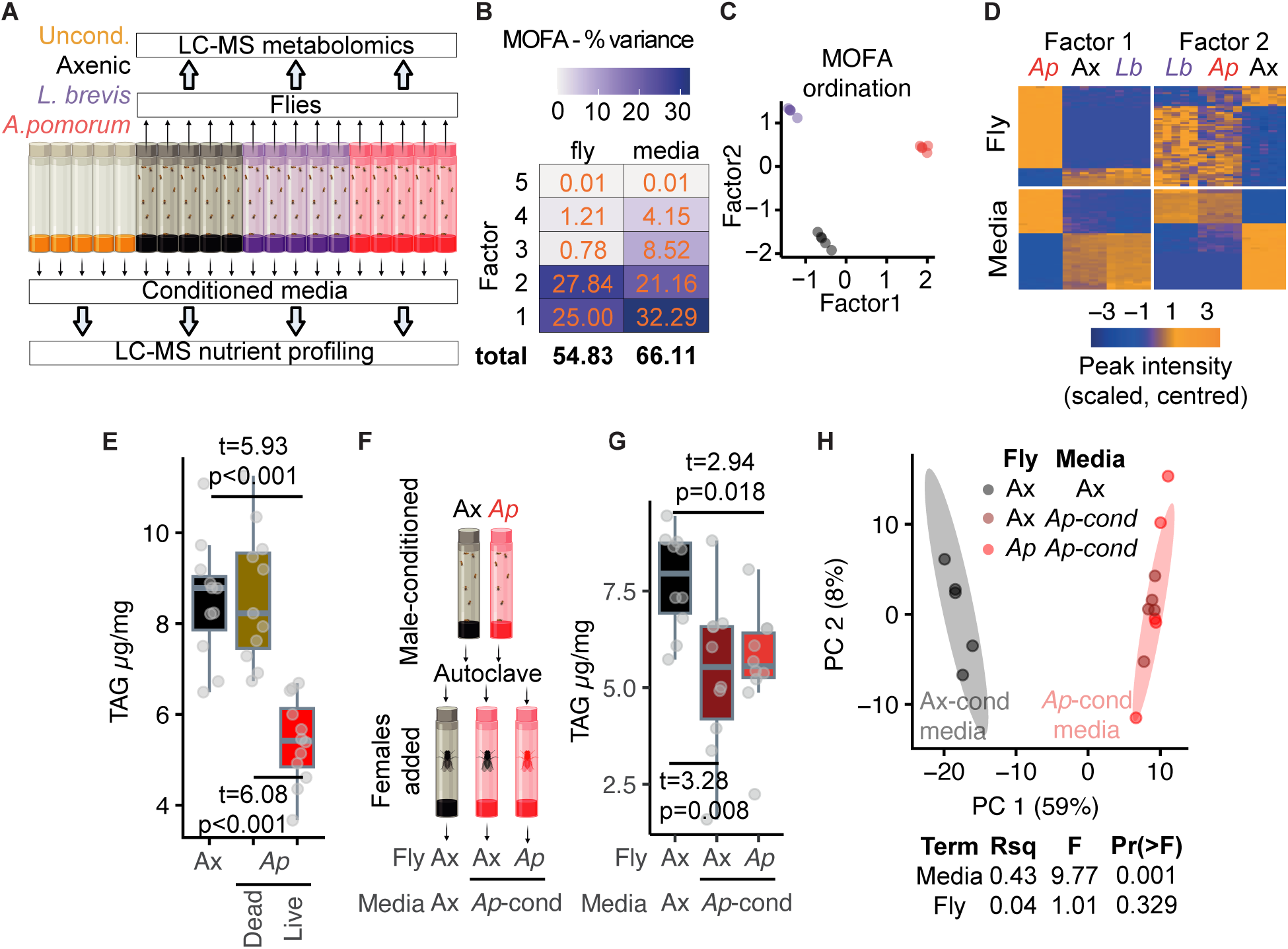
Fly metabolomic state corresponds to microbiota-dependent nutrition. **A.** LC-MS was used to profile paired (i.e. from the same container) metabolomes of flies and media nutrient content, following egg-to-adult culture. Unconditioned sterile medium was included as a baseline control. Media and adult flies were also profiled after rearing under axenic conditions, in the presence of *L. brevis*, or in the presence of *A. pomorum.* Microbiota not only inhabit the gut but also grow in fly food, giving an easily-measured substrate to identify how microbiota modulate host nutrition. **B**. Identifying axes of correlated variation between flies and media using Multi-Omic Factor Analysis (MOFA). MOFA was applied to sample-matched measurements of media and flies, to identify latent variables that explained correlated variation between the two sample types. Differentially-abundant peaks were included for the fly samples, all peaks were included for media. The heatmap shows variance explained in fly and media by the resulting five factors, summed beneath the plot, with ∼55% residual variance. **C.** The scatterplot shows each sample (i.e. the combined view of both media and fly) according to values on the latent variables represented by Factors 1 and 2. Factor 1 appears to differentiate impacts of *A. pomorum* from axenic and *L. brevis*-associated flies/media, while factor 2 appears to differentiate axenics from the bacteria-associated flies/media. **D**. MOFA identifies covariation between fly metabolome and media nutrient availability. The heatmaps show the intensities of the fifty top-ranked peaks (by weights) associated with factors 1 and 2. Each column represents an individual sample, with values from flies given in the upper heatmaps and corresponding values from food beneath. Columns are ordered by factor values, with grouping by microbial condition given above. Each row represents an LC-MS peak, ordered by hierarchical clustering. The correlated behaviours confirms that MOFA has identified clusters of peaks that covary between media and fly. **E.** Inoculating axenic medium and flies with dead *A. pomorum* (adult-onset, replenished on each transfer to fresh media/container) did not recapitulate the impact of live *A. pomorum* inoculation on TAG. **F.** Design of experiment to test causal relationships between microbiota-driven nutritional alteration and fly metabolome. Fly media were conditioned by axenic and *A. pomorum*-associated males, sterilized by autoclaving, then allowed to reset. Adult females were introduced to these media: *Ap*-females were fed *A. pomorum*-conditioned media, while axenic females were fed either Ax-fly-conditioned or *Ap*-fly-conditioned media. **G-H.** *Ap*-fly-conditioned food rescues axenic TAG levels (G) and metabolome (H), equivalent to the presence of live *A. pomorum*. Statistics from post-hoc EMMeans tests of ANOVA (E, G) and PERMANOVA (H).

To relate nutrition to metabolic state, we identified covariation using multi-omic factor analysis (MOFA2 ^40^). To relate nutrition only to microbiota-dependent metabolites, fly samples were restricted to differentially abundant peaks (FDR<0.05, *Ap-*flies or *Lb-*flies versus axenics). The top-two factors together explained ∼53% of fly metabolite variance, while subsequent factors explained negligible variance (Figure 2B). Factor 1 reflected *A. pomorum* effects, while Factor 2 reflected axenic effects (Figure 2C, Figure S9A-B). Heatmap analysis confirmed that Factors 1-2 captured real nutrition-metabolism covariation (Figure 2D). Thus a slight majority (∼53%) of metabolic responses to microbiota can potentially be explained by nutritional covariance, but a substantial remainder (∼47%) cannot.

Having established correlation, we tested whether nutritional effects were causal. We focused on *A. pomorum*, given its larger metabolic impact. Inoculating sterile flies and media with autoclaved *A. pomorum* (refreshed on each transfer to fresh container/medium) did not modulate TAG, suggesting that bacteria-associated molecular patterns (e.g. peptidoglycan or LPS) alone were dispensable (Figure 2E). We then tested effects of bacterial nutrition (Figure 2F-G) by pre-conditioning media with axenic or *A. pomorum*-colonised males (since females would contaminate with eggs/larvae), autoclaving, then re-feeding to test females (Figure 2F). Axenic females on *Ap-*conditioned media fully recapitulated the TAG phenotype of flies bearing live *A. pomorum* (Figure 2G). Encouraged by this result, we characterised the metabolome, finding that metabolic profiles of axenics on *Ap*-conditioned media were indistinguishable from flies with live *A. pomorum* (Figure 2H). Together these results suggested that nutritional change(s) explain metabolic impacts of *A. pomorum*.

### *A. pomorum* conditions flies to support bacterial growth under nitrogen stress

Characterising media composition allowed us to investigate what compounds the host might provide to support bacterial growth. *Acetobacter* are prototrophic for most nutrients, but they require nitrogen. Our media analysis (Figure 2A) showed that flies increased purines and purine breakdown products (henceforth “purines”): allantoin, hypoxanthine, inosine, and urate (Figure 3A). Previous work has also connected fly microbiota to host purine pools ^41^. *A. pomorum* can use purines as an exclusive nitrogen source ^42^. Elevated purines in axenic-conditioned medium must derive from fly excreta (frass); their reduction under gnotobiotic conditions (hypoxanthine and inosine) suggests either bacterial use or lesser output in colonised flies. Either interpretation is consistent with flies excreting a nitrogen source that could support *Acetobacter* growth.

**Figure 3.**
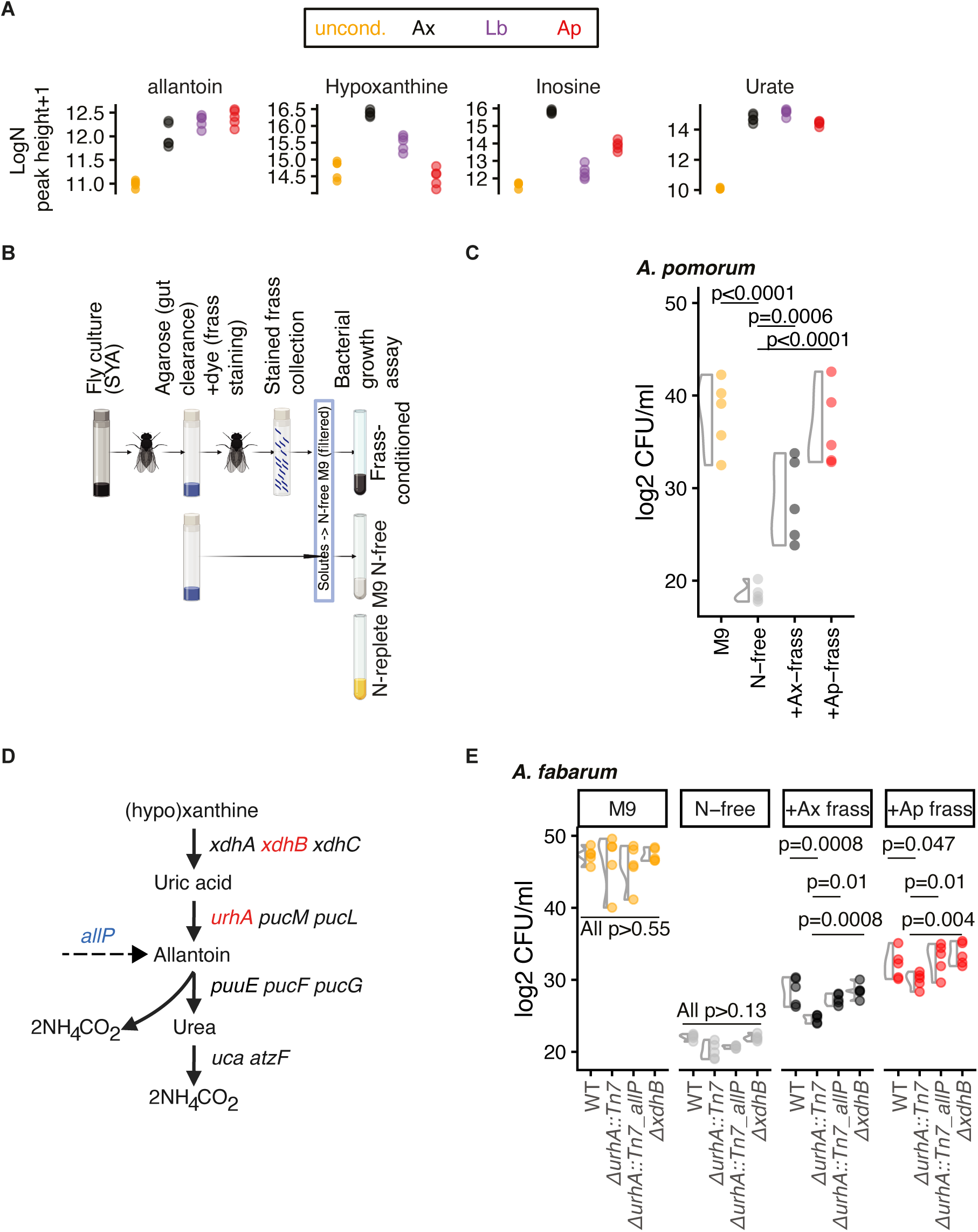
*A. pomorum* conditions flies to support bacterial growth under nitrogen stress. **A.** Flies enrich media with nitrogenous waste. Panels indicate LC-MS peak heights of nitrogen metabolites, identified against definitive standards, in unconditioned SYA medium, or media conditioned by axenic flies, *Lb*-flies, or *Ap-*flies. Significant differences were detected in at least one comparison for all metabolites shown. **B.** Assay design. Flies were cultured until adulthood then transferred to agarose containing blue dye for 10 hours to clear the gut of nutritive medium and replace with nutrient-free medium; before transfer to empty, sterile containers for 24h. Soluble metabolites from frass were collected by washing the containers with nitrogen-free M9L medium. N-free media was established by washing empty vials with M9L medium lacking nitrogenous base. Media were filter-sterilised and normalized before inoculation with *Acetobacter.* **C.** Fly frass solutes rescue *A. pomorum* in nitrogen-free conditions. Bacterial density (72h growth) was reduced in nitrogen-free medium relative to replete medium. Growth was rescued by solutes of frass of axenic flies, and further by frass of *Ap*-flies. ANOVA F_3,16_=28.86, p=1.1e-06, p-values on panel from post-hoc (emmeans) tests. **D.** The purine salvage pathway in *Acetobacter* spp. In subsequent experiment (panel E), genes highlighted in red are mutated in *A. fabarum*, gene highlighted in blue is complemented into *A. fabarum.* **E.** Function of purines in fly assurance of bacterial growth. Frass solutes collected as previously were used to culture *A. fabarum* wild-type, strain unable to salvage purines (*ΔurhA::Tn7*), a strain competent for allantoin utilization (*ΔurhA::Tn7_allP)*, and strain that can use urate but not hypoxanthine (*ΔxdhB)*. Capacity for bacterial growth to respond to media conditioning was strain-dependent, with a requirement for purine salvage in order for axenic frass to fully rescue bacterial growth, and a marginal requirement for full rescue by *Ap*-fly frass. P-values (FDR) from post-hoc (emmeans) tests.

While little is known about what animals provide to gut microbiota, it has been established that hibernating ground squirrels, which do not eat, secrete urea into the gut lumen, which the microbiota metabolise into amino acids that are available to the host ^31^. Our nitrogen provision hypothesis in flies mirrors this process, hinting at an evolutionarily conserved mechanism.

We tested *in vitro* whether frass solutes can rescue *Acetobacter* growth in otherwise nitrogen-free conditions, by clearing media from fly guts with agarose feeding (containing a dye that does not cross the gut epithelium), dissolving frass in nitrogen-free M9L medium, normalising solute density (with the dye), filter-sterilising, and then growing *A. pomorum* in the resulting frass-solute conditioned minimal medium (Figure 3B). In unconditioned M9L medium, *A. pomorum* growth was blocked by omitting nitrogen (ammonium) (Figure 3C). Axenic frass solutes partially rescued this deficit, as expected with solutes from urine, but strikingly *Ap-*fly frass rescued *A. pomorum* growth fully (Figure 3C). This suggested that the impacts of *A. pomorum* on host metabolism modify excretion in a way that potentially safeguards their own growth under nitrogen-limited conditions, indicating a metabolic reciprocation from the host that potentially underpins symbiosis.

To assay purines’ involvement, we used purine salvage mutants in the congeneric *Acetobacter fabarum*, which, unlike *A. pomorum*, lacks allantoin permease, making (hypo)xanthine/uric acid metabolism mutants into purine auxotrophs (Figure 3D). We used Δ*urhA* (cannot use any purines), Δ*urhA::Tn7_allP* (expressing allantoin permease to rescue purine use), and ΔxdhB (urate salvage) (Figure 3D). All strains grew equally in replete M9L, i.e. the mutations did not impair growth in optimal conditions (Figure 3E). Growth was also equally impaired in N-free M9L. Frass solutes rescued growth, but this rescue was significantly diminished in *ΔurhA*, indicating a partial role for purines. Rescuing either allantoin (*allP* expression) or urate (*ΔxdhB*) metabolism rescued the growth defect (Figure 3E). This suggested that the metabolic feedback from the fly partially involves purines, which could promote bacterial growth in ecological contexts where nitrogen is scarce. The parallels between this host-to-microbe nitrogen provision and that in ground squirrels ^31^ suggests phenomenology that may be widespread and should be characterised in other systems.

### Microbiota reprogram the fly metabolome

Next, we asked how *A. pomorum* and *L. brevis* impact host metabolism. We combined three independent replicate experiments (Supplemental note 1, Figure 2), for a robust and comprehensive analysis. We manually curated 130 metabolites identified in all three experiments, batch-corrected, and confirmed with an unsupervised analysis that microbiota-dependent variation was consistent across studies (Figure S10A).

First, for an integrative overview, we performed Multivariate Integrative supervised Partial Least Squares-Discriminant Analysis (MINT-sPLS-DA, henceforth MINT), to identify metabolites that repeatably explained microbiota-dependent variation across experiments. MINT separated *Ap*-flies from axenics/Lb-flies on component 1, with *Lb*-flies partly distinguished on component 2 (Figure 4A). Component 1 loadings implicated pantothenate (vitamin B_5_), lysine (and a lysine ion), xanthine, β-imidazolelactate, and kynurenine as distinguishing *Ap*-flies (Figure 4B); component 2 loadings implicated a purine metabolite (annotated as “allopurinol,” likely hypoxanthine), phenylalanine, ornithine, glycyl-hydroxyproline, adenine, L-β-homolysine, and (iso)nicotinate as distinguishing Lb-flies (Figure 4C). In a complementary differential abundance analysis (i.e. per metabolite) we found 31 increased/17 decreased metabolites in *Ap-*flies (Figure S10B), and seven increased/six decreased in *Lb-*flies (Figure S10C), including metabolites consistent with MINT. To interpret biological impacts, we determined KEGG pathway enrichment using FELLA, which propagates networks of reactions/enzymes/pathways/modules from shared metabolite neighbours, as appropriate for sparse metabolomic data. FELLA produced cohesive networks, ie. differentially-abundant metabolites were in shared pathways (Figure S11).

**Figure 4.**
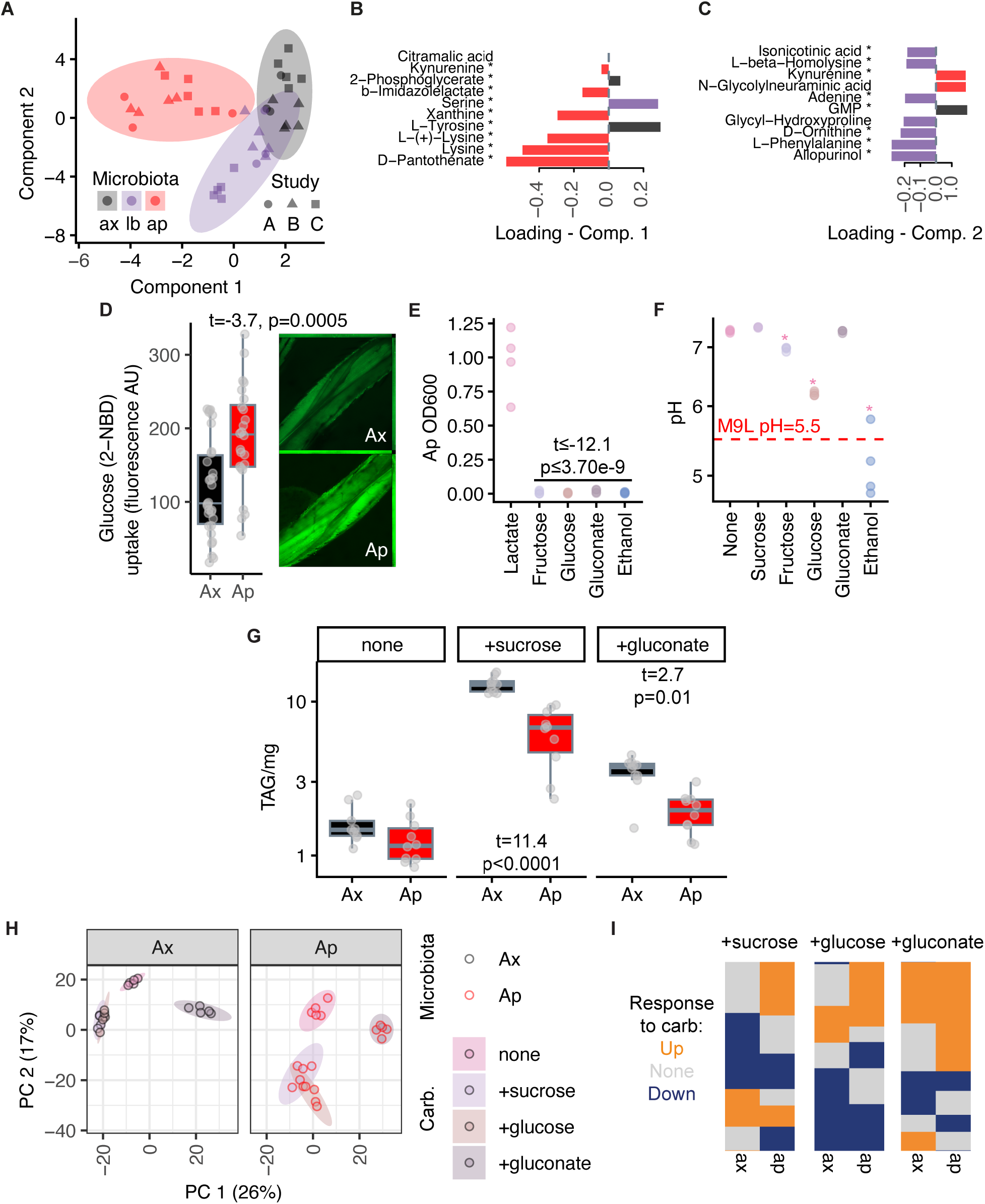
Microbiota reprogram the fly metabolome. Data for 130 metabolites were curated and batch-corrected from preceding LC-MS analyses (Figure 2, Supplemental note 1). **A-C.** MINT-sPLS-DA differentiates axenics, *Ap*-flies and *Lb*-flies (A) on axes loaded with various definitively-identified metabolites (B, component 1; C, component 2). Asterisked metabolites identified from fragmentation data, others by comparison to definitive standards. KEGG pathway and module enrichment among these differentially-abundant metabolites is presented in Figure S12. **D.** *A. pomorum* promotes glucose uptake. Five-day-old axenics and *Ap*-flies were injected with a glucose analogue (100nL 5 mM 2-NBDG, 30min subsequent incubation on 1% agar) that fluoresces upon uptake into cells and is minimally metabolized thereafter, before dissection and imaging. Representative micrographs show transcuticular fluorescence in leg muscle, quantification gives statistics from t-test, suggesting enhanced *Ap*-fly capacity for glucose metabolism. **E.** *A. pomorum* is selective for carbon source in minimal (M9) media. *A. pomorum* were sub-cultured into media containing only indicated compounds as carbon source. Growth (OD_600_ nm) was assayed after 72 hours. For all comparisons between lactate medium (positive control) and indicated conditions, t≤-12.1, p≤3.7e-9 (ANOVA). **F.** *A. pomorum* selectivity for energy source in M9L media. pH change was measured as a proxy for energy source oxidation. With no added energy source (none), *A. pomorum* uses lactate, increasing pH relative to baseline. Positive control (ethanol) reduces pH, consistent with acetate production (Figure 1). Adding glucose or fructose reduces pH, consistent with sugar acid production via oxidation. Adding sucrose or gluconate does not decrease pH, consistent with genomic prediction (Figure S14B). Asterisks indicate significant differences (ANOVA, p<0.05) relative to no added energy source. **G.** *A. pomorum* alteration of host TAG response to gluconate suggest the bacteria can reprogram host energy metabolism. Boxplots show TAG levels of flies (axenic/*Ap*-flies) fed the indicated carbohydrates above on from day 3-10 adulthood (none=yeast+agar, +sucrose=yeast+agar+146mM sucrose as in prior figures, +gluconate=yeast+agar+292mM gluconate). ANOVA microbiota*diet F2,53=32.60, p=5.9e-10. Statistics on graph show effect of microbiota (post-hoc emmeans tests), when significant. Post-hoc tests also show that gluconate increases TAG in axenic flies (t=3.14, p=0.008), but not in *Ap*-flies (p=0.5). **H-I.** Microbiota reprogram host metabolomic response to dietary carbohydrate variation. Flies were fed varied carbohydrate sources (diets as panel G, +glucose=yeast+agar+292mM glucose), days 3-10 of adulthood. **H**. shows PCA: dietary variation is apparent in axenics, but substantially more pronounced in the presence of *A. pomorum* (PERMANOVA, Carb * microbiota p≤0.001). **I**. shows peaks that are significantly up/downregulated by the specified carbohydrates in axenics/*Ap*-flies; showing that for many peaks the response to carbohydrate is contingent on *A. pomorum*. Differential abundance (FDR≤0.05) determined by Integrated Metabolomics Pipeline.

FELLA suggested that *L. brevis* perturbed purine metabolism/PRPP biosynthesis (Figure S12A-B). *A. pomorum* perturbed central energy metabolism (reduced pools of glycolysis/gluconeogenesis/TCA cycle metabolites) and amino acid metabolism (reduced pools), alongside glutathione/PRPP/polyamine/CoA biosynthesis and pyrimidine degradation (increased metabolite pools) (Figure S12A-B). Notably, *A. pomorum* reduced pools of glycolysis/TCA metabolites while increasing pools of pentose-phosphate pathway metabolites (Figure S12A-B). Since these pathways metabolise glucose and share intermediates, their divergence suggested that *A. pomorum* reprograms the balance of pathway activity.

To test whether reduced TCA/glycolysis metabolites in *Ap*-flies was due to deficient glucose uptake, we injected 2-NBDG, a derivative that fluoresces upon cellular uptake, and quantified fluorescence in muscle visible through translucent leg cuticle. We dropped *L. brevis* from these experiments, given its comparatively modest metabolic effects, focussing on *A. pomorum*. However, *Ap*-flies fluoresced more strongly than axenics (Figure 4D), suggesting increased glucose uptake and arguing that reduced levels of TCA/glycolysis metabolites cannot be explained by glucose limitation and may, therefore, indicate reprogrammed carbohydrate use. Accordingly, *A. pomorum* reprogrammed the kinetics of TAG storage upon elevated dietary sugar, and mobilisation upon starvation, with corresponding effects on starvation resistance that required *Akh*, the fly orthologue of glucagon (Supplemental note 2, Figure S13). These altered kinetics further indicated reprogrammed carbohydrate metabolism.

Metabolic reprogramming implies altered nutrient allocation to downstream metabolites/stores. To test this notion, we sought to experiment with a nutrient that flies could metabolise but microbiota could not, so that any microbe-dependent differences host response could not be attributed to pre-ingestive microbial depletion/modification. *Acetobacter* spp. prodigiously oxidise extracellular glucose to gluconate (Figure S14A), which some oxidise further via D-gluconate dehydrogenase ^28^. However, the *A. pomorum* genome lacks both this enzyme and a transporter for D-gluconate, suggesting that extracellular gluconate is unusable waste. Further genomic analysis also suggested that *A. pomorum* can derive energy, but not carbon, from glucose, fructose, and ethanol (Figure S14B). We confirmed these predictions empirically. In defined bacterial medium (M9L), bacteria could not grow (acquire carbon) when lactate was replaced with gluconate/sugars/ethanol (Figure 4E). When lactate was included, fructose/glucose/ethanol were oxidised (media acidification) with variable efficiency, but not gluconate or sucrose (Figure 4F). Inability to use sucrose was consistent with genomic absence of glucosidases/sucrose transporters (Figure S14B), implying that *A. pomorum* in fly cultures requires flies to hydrolyse sucrose to fructose/glucose. More importantly, inability to use gluconate confirmed it as waste that *A. pomorum* cannot use.

We incorporated gluconate into fly medium (with carbohydrate-free and sucrose-containing media as controls) and tested impact on energy storage (Figure 4G). Positive controls with sucrose behaved as previously. Without gluconate or sucrose, *A. pomorum* did not impact TAG, confirming that carbohydrate is required. Adding gluconate increased axenic TAG (t=3.14, p=0.008), confirming flies can use it. However, *A. pomorum* abrogated gluconate’s effect, indicating bacterial reprogramming of host gluconate metabolism. We also found that *A. pomorum* capacity for sugar usage did not predict impact on TAG across diets incorporating various single sugars (Figure S14C), arguing against impacts via pre-ingestive bacterial sugar metabolism. In fact, LC-MS metabolome profiling on carbohydrate-free/sucrose/glucose/gluconate-containing diets (Figure 4H-I) showed lesser carbohydrate-dependent variation in axenics, but variation was released in *Ap-*flies, suggesting that *A. pomorum* amplified host sensitivity to carbohydrate variation, including gluconate (Figure 4H, Figure S14D). Differential abundance analysis also showed that signs of responses to dietary carbohydrates differed significantly between axenics versus *Ap-*flies (Figure 4I), i.e. responses of downstream compounds to dietary carbohydrate were microbe-dependent.

Metabolic reactions are interconnected, generating a network structure. Metabolic reprogramming should change this structure, analogous to microbial restructuring of host transcriptional networks ^29^. Metabolic network structure was apparent within each microbiota condition (clusters of correlated metabolites); but these structures did not transpose across conditions (Figure S15), suggesting microbiota-dependent restructuring. We tested this explicitly with Joint Graphical Lasso (JGL), which shares information across conditions to reveal consensus and condition-specific structure, then quantifies conditionality. JGL confirmed strikingly different network structures between microbiota conditions (Figure S16), consistent with metabolic reprogramming.

We asked whether reprogrammed metabolism could potentially be explained by altered expression of underpinning metabolic genes, via metabolic modelling. Differential gene expression and enrichment analyses found altered expression of genes involved in energy metabolism and respiration in both *Ap-*flies and *Lb-*flies (Figure S17-S19), indicating a bacterium-nonspecific response. Generating sample-specific genome-scale metabolic models (GEMs) suggested broad-scale responses to bacteria (Figure S20A), which propagated to gene-protein-reactions (Figure S20B), with significant impacts on subsystems involved in metabolism of lipids, acyl-coenzyme A, carbohydrates, purines, and folate (Figure S20C). But differential flux-balance analysis, underpinned by various robustness analyses indicated that gene expression differences did not predict differential flux (Supplemental note 3, Figure S20D, Figure S21). We conclude transcriptional changes may play roles in metabolic remodelling, but differences in specific metabolic fluxes are likely stoichiometric rather than transcriptionally-regulated.

Together, these results showed *A. pomorum* reprogramming dynamic host metabolic processes.

### Pantothenate is modulated by microbiota in flies and mice

Our metabolome data strongly highlighted a candidate cause of metabolic reprogramming. Pantothenate (vitamin B_5_), the substrate for *de novo* coenzyme A (CoA) synthesis, was increased in *Ap-*flies (Figure 5A). CoA esters, particularly acylated species (acyl-CoAs) are central signalling metabolites. For example, acetyl-CoA participates in ∼10% of eukaryote enzymatic reactions, including TCA cycle and lipid biosynthesis, plus regulatory processes including protein acetylation ^43,44^. Acyl-CoAs enter mitochondria for energy production via beta-oxidation, while malonyl-CoA donates carbons for fatty acid synthesis and inhibits beta-oxidation. As an essential nutrient that animals cannot synthesise, pantothenate fits the bill of the sort of nutrient that hosts would be selected to acquire from microbiota.

**Figure 5.**
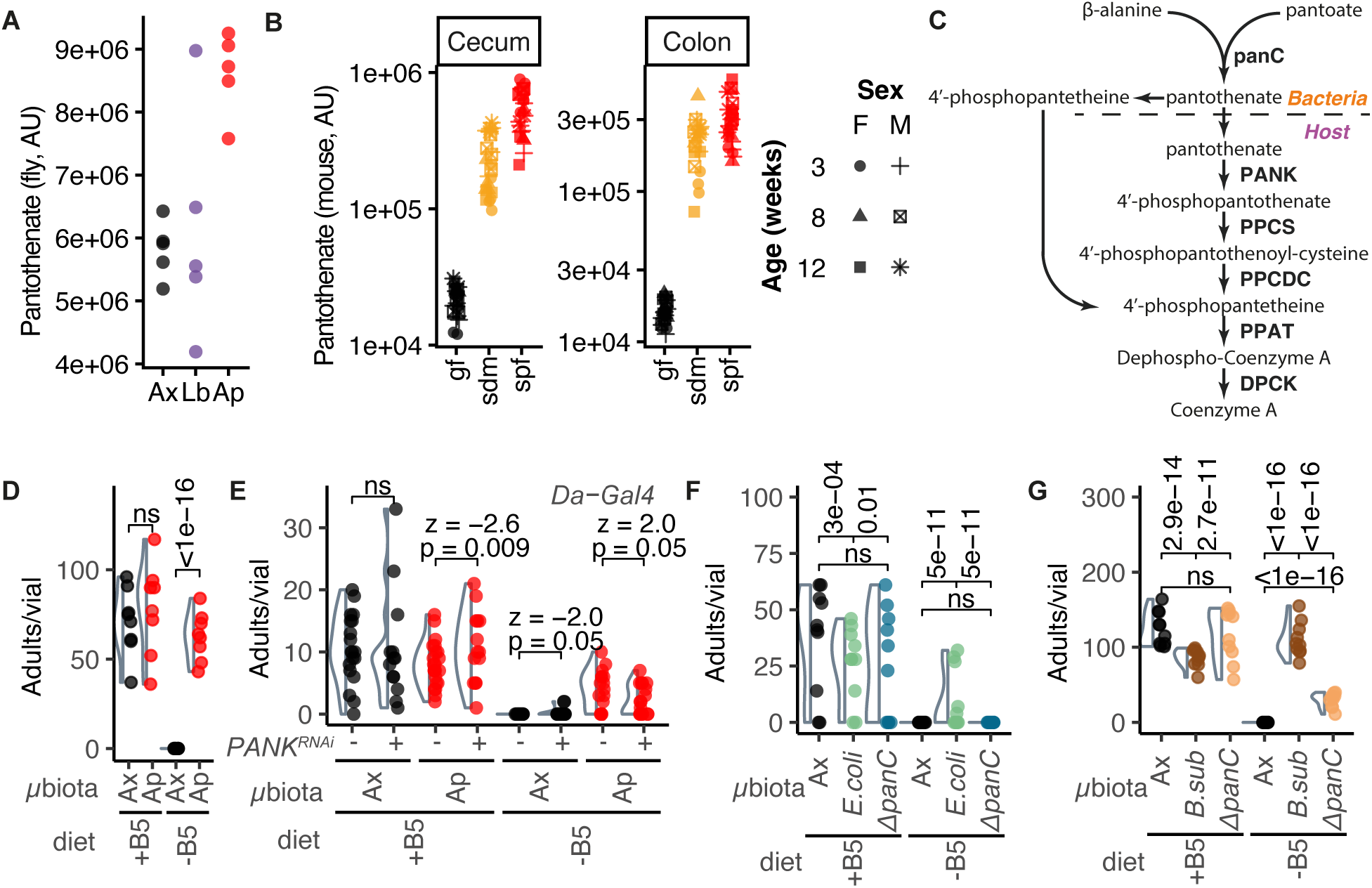
Pantothenate is modulated by microbiota in flies and mice. **A.** *A. pomorum* increases pantothenate in adult female flies. LC-MS peak intensities shown per sample (pools of 5 flies), identified against definitive standards, in 3-day-old adult female axenic, *L. brevis*, and *A. pomorum* gnotobiotic flies. Ap vs axenic FDR<0.0001; Lb vs axenic not significantly different. **B.** Microbiota increase pantothenate in mice. Data from ^45^. LC-MS intensities shown for tissues in which microbiota explain substantial variance. gf=germ-free, sdm=standardized (OMM12) microbiota, spf=specific pathogen-free. Point shape denotes mouse age and sex. **C.** Model of microbe-to-host pantothenate provision, with pantothenate/4’-phosphopantetheine synthesis in *Acetobacter* shown at top, interfacing with host CoA biosynthesis beneath. **D.** *A. pomorum* rescues fly viability in pantothenate-free defined (holidic) diet. Adult counts taken 21 days post egg deposition on pantothenate free medium (B_5_-) and replete medium (B_5_+). Zero-inflated GLM, microbiota*diet F=114.15, p<0.0001. Statistics on figure from post-hoc tests (emmeans), stratified per diet. Violin plots show density per condition. **E.** *A. pomorum* rescue of fly viability is blunted by pantothenate kinase (PANK) RNAi knockdown (PANK-i). The experiment performed in panel D was replicated in presence/absence of PANK-RNAi. All flies bore *Daughterless-Gal4*, ±*UAS-PANK-RNAi* for knockdown. Zero inflated GLM, microbiota*diet*PANK-RNAi F=8.6, p=0.004. Statistics on figure from post-hoc tests (emmeans), stratified per diet. Violin plots show density per condition. **F.** Rescue of fly viability by *Escherichia coli* requires pantothenate biosynthesis, pantothenate auxotroph (Δ*panC*). The experiment in panel D was replicated with *E. coli* wild-type (BW25113) and Δ*panC*. Zero inflated GLM, microbiota*diet F=22.77, p<0.0001. Statistics on figure from post-hoc tests (emmeans), stratified per diet. Violin plots show density per condition. **G.** Rescue of fly viability by *Bacillus subtilis* requires pantothenate biosynthesis, pantothenate auxotroph (Δ*panC*). The experiment in panel D was replicated with *B.subtilis* wild-type (168) and Δ*panC*. Zero inflated GLM, microbiota*diet F=354.97, p<0.0001. Statistics on figure from post-hoc tests (emmeans), stratified per diet. Violin plots show density per condition.

First, to test whether pantothenate provision is evolutionarily conserved, we looked to public mouse metabolome data from germ-free (GF), specific pathogen-free (SPF), and standardised OMM12 microbiota (STM) conditions ^45^. Pantothenate was elevated by microbes in both colon and cecum (Figure 5B). This effect appears physiologically relevant because a second independent study found elevated pantothenate in the brain ^46^. Metagenomic analysis of human microbiota reveals pantothenate biosynthesis to be one of the most highly represented pathways ^47^. Microbial pantothenate provision in animals separated by hundreds of millions of years of evolutionary divergence suggests a potentially ancient function.

We returned to the fly to test whether elevated levels in *Ap*-flies were due to greater supply, or reduced use. *A. pomorum* can synthesise pantothenate ^26^, hinting at provision. We generated pantothenate-free defined fly diet, on which axenic eggs were inviable, but *A. pomorum* provided full rescue (Figure 5D). Expressing RNAi against the first enzyme in *de novo* CoA synthesis (Figure 5C), pantothenate kinase (PANK), blunted rescue (incomplete knockdown was anticipated since RNAi is rarely 100% efficient) (Figure 5E). Direct provision was also suggested by mutating *panC*, the terminal step of bacterial pantothenate synthesis (Figure 5F–G). We could not make *panC* mutants in *Acetobacter*, (likely because they express no pantothenate transporter, making *panC* essential), so we used *Escherichia coli* and *Bacillus subtilis* as substitutes (Figure 5F-G). Wild-types rescued pantothenate-free fly development, but *ΔpanC* abolished (*E. coli*) or diminished this rescue (*B. subtilis*, likely due to suppressor mutations ^48^). *A. pomorum* also rescued dietary riboflavin (B_2_) deficiency (Figure S22A) ^26,49^. Collectively, these experiments demonstrated that *A. pomorum* can provide physiologically sufficient pantothenate.

### Metabolic positive feedback between *Drosophila* and *Acetobacter*

We established the significance for both the fly and microbiota of the host metabolising bacterial pantothenate. In the fly, we quantified unesterified CoA (CoASH) and short-chain acyl-CoA species. Sum total levels (CoASH+(acyl)CoAs) almost doubled in *Ap-*flies (Figure 6A, 1.85-fold mean increase). Integrating acyl-CoA species, PCA differentiated *Ap-*flies from axenics (Figure 6B). Specifically, *A. pomorum* increased succinyl-/propionyl-/glutaryl-CoA (5% FDR) (Figure 6C). By comparison, *L. brevis*, which did not increase fly pantothenate (Figure 5A) increased only succinyl-CoA (Figure S22B–D). These data indicated that *A. pomorum* pantothenate provision impacts host CoA pools.

**Figure 6.**
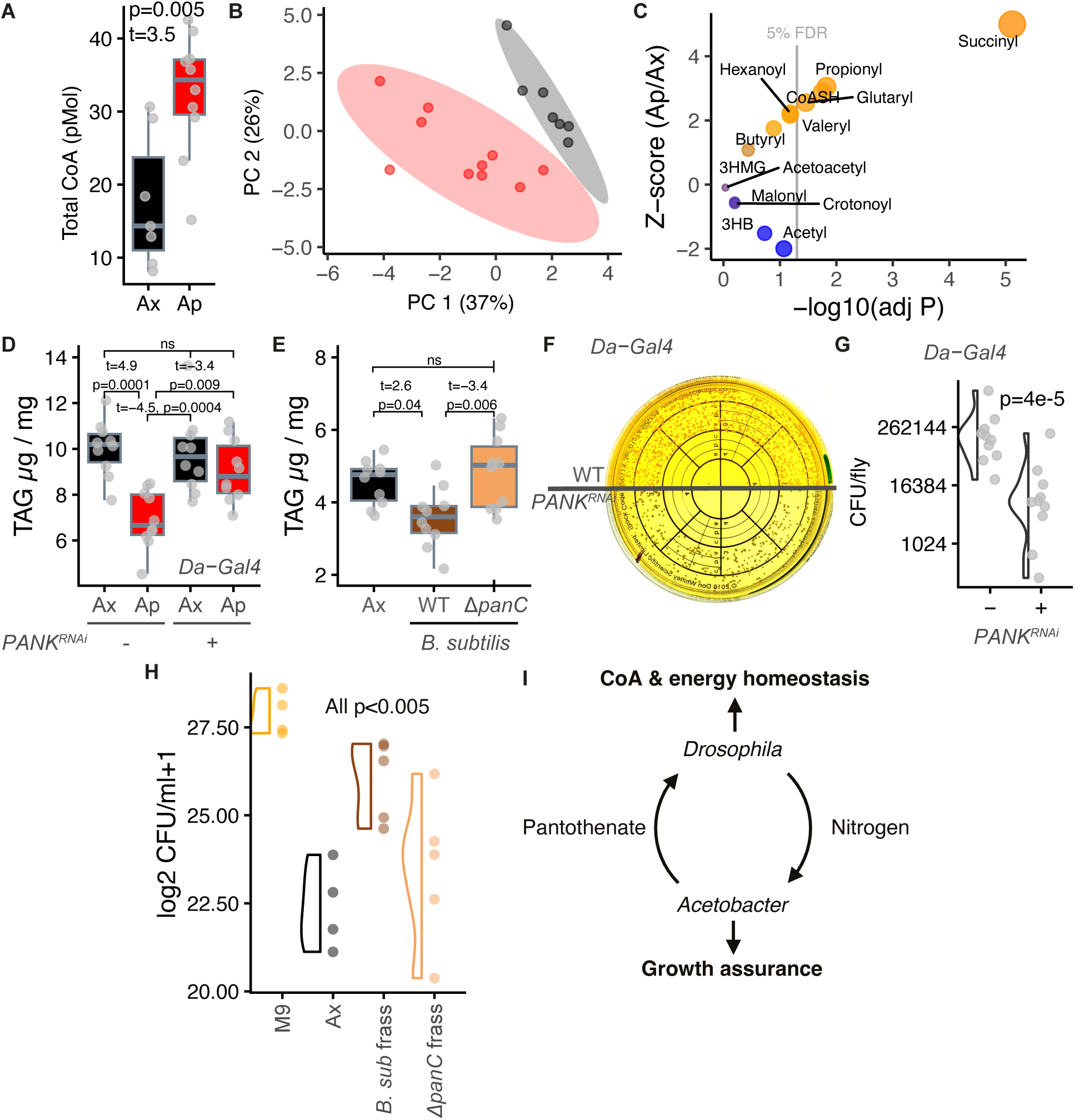
Metabolic positive feedback between *Drosophila* and *Acetobacter*. **A.** Sum total CoA (CoASH and (acyl)CoA species) is increased in *Ap-*flies (three-day-old adult females). Statistics from t-test. **B.** PCA indicates multivariate differences in CoA levels between axenics and *Ap-*flies. **C.** Multiple CoA species are increased by *A. pomorum*, corresponding to increased pantothenate. Z-scores (shown on Y axis and by point colour) show standardised differences between *Ap-*flies and axenics, for each CoA species. More significant differences are shown by higher x-axis values. Vertical lines show significance cutoffs at 5% and 10% FDR (i.e. adjusted p=0.05 or 0.1, respectively). **D.** PANK knockdown (PANK-i) blocks reduction of host TAG by *A. pomorum*. Data from three-day-old females. ANOVA, microbiota*PANK-i F_1,36_=7.24, p=0.01. **E.** Bacterial (*B. subtilis*) *panC* is required for rescue of host TAG. ANOVA F_2,27_=6.37, p=0.005. **F-G.** *A. pomorum* density is reduced in flies with deficient pantothenate metabolism (*PANK^RNAi^*). F: Wild-type shown at top, PANK-RNAi beneath in a composite plate image. For all panels, post-hoc tests were applied using emmeans. G: Quantification. p-value from Wilcoxon rank-sum test. **H.** Flies receiving pantothenate from the microbiota support greater bacterial growth under nitrogen stress. Frass solutes were collected as previously from axenics, and gnotobiotic flies monoassociated with *B. subtilis* wild-type strain *(Bsub)* or pantothenate biosynthesis mutant *(ΔpanC)*. Solutes from frass of *Bsub*-flies better promoted *A. pomorum* growth than solutes from axenic frass, yet the full rescue required pantothenate biosynthesis. GLM (quasipoisson family) p=4.1e-5, all sequential comparisons p<0.005. **I.** Conceptual model. *Drosophila* and *Acetobacter* mutualism is underpinned by reciprocal metabolic provision.

We tested whether pantothenate was required for *A. pomorum* to modulate fly TAG. *PANK^RNAi^* blocked *A. pomorum* rescue of host TAG (Figure 6D), indicating a role for bacterial modulation of host CoA. We also tested effects of bacterial pantothenate provision, using *B. subtilis* (*E. coli* does not reduce fly TAG, Figure S22E, ^28^), with a wild-type that rescued TAG equivalently to *A. pomorum* (Figure S22F). A second experiment showed that pantothenate synthesis (*panC*) was required for this rescue (Figure 6E). We also asked the complementary question of whether pantothenate was sufficient for TAG rescue. Our metabolome data showed that *A. pomorum* modulated a suite of host metabolites, suggesting that pantothenate alone would not be sufficient, and indeed TAG was not altered in axenics provided with a titration of either pantothenate or a cocktail of B-vitamins (Figure S22G). Pantothenate is thus necessary but not sufficient for microbial effects on TAG, indicating a hub among a suite of microbe-dependent metabolic changes.

We then examined how bacterial pantothenate provision feeds back to the bacteria via the host, hypothesising that it would render flies more permissive. Expressing *PANK^RNAi^* in *Ap-*flies reduced bacterial colony-forming units ∼10-fold (Figure 6F–G), suggesting that host pantothenate metabolism is mutually beneficial. In the context of our prior experiments showing that flies can provide microbes with nitrogen (Figure 3), we hypothesised a reciprocal pantothenate-for-nitrogen exchange. We reared flies with *B. subtilis* wild-type, collected frass solutes, and tested whether *in vitro* promotion of bacterial growth in N-depleted media was wild-type or Δ*panC* dependent (Figure 6H). Frass from wild-type-colonised flies rescued growth better than axenics, but bacterial pantothenate synthesis (*panC*) was required for full rescue. These data are consistent with pantothenate brokering reciprocal metabolic feedback, reprogramming host metabolic homeostasis, increasing nitrogen excretion, which can enhance bacterial growth in low nitrogen conditions (Figure 6I).

## Discussion

The conservation of host metabolic responses to microbiota suggests evolutionarily ancient dependence. A parsimonious mechanistic explanation is that microbiota provide essential nutrients that animals require for core cellular functions but cannot synthesise themselves. Association with gut microbiota might solve this problem. Our finding of a central role for pantothenate provision is consistent with this logic, suggesting that hosts benefit from a gain-of-function in metabolic homeostasis mediated by CoA. Further, natural selection should favour mechanisms for host association with beneficial bacteria; our finding of pantothenate-dependent reciprocation from the host to the microbiota appears to illustrate a mechanism for this symbiotic reinforcement. We have every reason to expect similar roles in other systems, because pantothenate provision appears to be a conserved function of gut microbiota, and animals cannot synthesise pantothenate directly. Importantly, CoA also impacts directly on bacterial physiology ^50^.

Our analysis characterised metabolomic impacts of specific gut bacteria on *Drosophila*, indicating that *L. brevis* has modest impacts compared to *Acetobacter* spp. *L. brevis* has limited biosynthetic capability, competent to synthesise only a few essential nutrients, including vitamins B_2_ and B_3_, but not essential amino acids, yet it does appear to impact metabolic gene expression in a broadly similar fashion to *A. pomorum*. This suggests generalised remodelling of metabolic gene expression, independent of bacterial taxa, but a simultaneous change in substrate availability is required for metabolic effects of this changed transcriptional staging. Results suggest that *L. brevis* cannot provide those required substrates, leaving the host poised but unable to respond, which may account for transcriptional and metabolomic signatures of stress. In contrast, *A. pomorum* appears to have multifaceted impacts on host metabolism. We and others ^22,26,34^ show that *A. pomorum* provides numerous essential nutrients (with possible strain-specific effects ^26^), changing the hosts’ nutritional landscape. Our nutrient-transmission experiments now suggest that these nutritional effects are sufficient to account for *A. pomorum* metabolomic impacts *in toto*. However, we also found that *A. pomorum* reprograms how the host uses nutrients: this apparent paradox can be resolved by bacterial provision of a nutrient that changes how other nutrients are used. Indeed, our results show that *A. pomorum* provide substrate (pantothenate) for *de novo* synthesis of central signalling molecules (acyl-CoAs) that act as essential cofactors throughout energy metabolism.

Our results support a central hypothesis in symbiosis: that bacteria direct the fundamental architecture of host biology. While microbial effects on fly metabolism have been linked previously to interactions with dietary carbohydrate ^27,28,51^, we have now made the advance of showing that *A. pomorum* regulates host physiological response to carbohydrate that the bacteria cannot themselves access, which can only be explained by reprogramming host carbohydrate metabolism. Pantothenate and subsequent CoA provision provide an elegant explanation ^50^. CoA is involved in carbohydrate and lipid processing, acetyl-CoA powers the TCA cycle, sits at the nexus of sugar and fatty acid oxidation, and is a precursor for fatty acid biosynthesis ^52,53^. These functions are consistent with (A) diminished metabolic impact of *A. pomorum* in the absence of dietary carbohydrates (Figure 4G), and (B) repeatable signals that microbiota influence host B-vitamin nutrition ^54^.

A second central hypothesis in symbiosis is that hosts should cultivate their microbiota. This hypothesis is relatively untested in facultative symbioses in animals, though it has been demonstrated between plants and symbiotic mycorrhizae ^55,56^, and in obligate animal symbioses e.g. pea aphids and *Buchnera* ^57^. Honeybees can provide organic acids to intestinal *Snodgrasella alvi* ^30^. Most relevant is evidence that ground squirrels secrete urea into the gut lumen during hibernation, which the bacteria recycle into amino acids that are used by the host, mitigating potential uremic toxicity and ensuring a nitrogen/protein source for both partners while the host is not feeding ^31^. In this context, we speculate that host-to-microbe nitrogen provision may occur in multiple systems, including animals and plants. Excreta of *Ap*-flies fully spared bacteria of the requirement for exogenous nitrogen, but axenic excreta only partially rescued growth: bacterial colonization generated a state in the fly that promoted bacterial growth. This indicates a novel feedback loop, since the host provision to the bacteria appears contingent on bacterial pantothenate provision. Altogether, our results show that hosts and microbes can engage in symbiosis-reinforcing positive feedbacks. Mechanistically, the currencies exchanged between hosts and microbiota in this interaction (pantothenate and nitrogen) appear conserved between *Drosophila* and mammals (mice and ground squirrels, respectively), outlining a potentially conserved metabolic basis for mutualism.

Further work is required to identify mechanistically how microbiota-derived CoA impacts host metabolism. Our finding that pantothenate provision is necessary but not sufficient to rescue TAG levels in axenics indicates a multi-step process, e.g. involving *Acetobacter* simultaneously modulating sugar availability, transcriptional control of metabolic programs, other B-vitamins that impact related bioenergetic pathways, mitochondrial biogenesis, etc. A number of genes and tissues may be involved. *Drosophila* encodes a single PANK 1-3 (manipulated above) orthologue, and a fly PANK4 orthologue has recently been identified (dPANK4) that lacks the pseudokinase domain, but retains the phosphatase domain to target 4’-phosphopantetheine, negatively regulating CoA levels ^58^. B_5_ promotes CoA production in the Malpighian tubules, which in turn maintains gut homeostasis via a cascade of pathways dependent on isoprenoid biosynthesis, and this is balanced by the opposing effects of PANK and dPANK4 ^58^. dMyc is a pleiotropic transcription factor that regulates *Drosophila* growth and size as well as metabolism including glycolysis and glutaminolysis ^59^, and it regulates the expression of both PANK and dPANK4, promoting expression of the former and reducing expression of the latter ^58^. In mammals, Myc has been implicated in B_5_ metabolism, increasing uptake via the Smvt transporter ^60^, of which flies have an orthologue. Further, different *Acetobacter* species have been shown to promote ISC proliferation ^61^. dMyc also enhances the uptake of glucose in a *Drosophila* tumour model ^62^, and we demonstrate that *A. pomorum* also enhances glucose uptake. Together these findings generate the hypothesis that *A. pomorum* could potentially be shaping host B_5_ and energy metabolism via dMyc. These hypotheses need to be further tested.

We have mostly characterised metabolic regulation in whole flies, but flies have differentiated tissues which are functionally analogous to vertebrates’ ^63^. The pantothenate-dependent microbial modulation of host lipids implicates roles in the fly gut and adipose (fat body) – the main lipid stores – but CoA is required for metabolic reactions throughout all tissues, suggesting influence on distal tissues. Functions other than TAG storage are likely mediated by impacts of altered CoA pools in other tissues, which remain to be characterised.

In summary our data support two central hypotheses in symbiosis, showing that bacteria reprogram animal metabolism to support homeostasis, and bacterial provision of the metabolic substrate for these processes (pantothenate) is reciprocated with bacterial growth substrate (nitrogen/purines). Conservation of these processes indicates fundamental mechanisms for metabolic and symbiotic homeostasis.

## Materials and Methods

### Fly culture and media

All flies (conventional, axenic, and gnotobiotic) were *Wolbachia*-free *w^Dah^* maintained at 25°C on a 12hr light/dark cycle. For non-defined diet experiments, 5% Sucrose (Fisher), 10% Yeast (MP Biomedicals, lot no. S6853), and 1.5% Agar (Sigma) (SYA) was used for all experiments. For experiments requiring preservatives 0.3% nipagin and propionic acid were incorporated. Mated females were used in all experiments unless otherwise specified.

To generate axenics, adults laid eggs on juice agar (2% agar, 2.6% sucrose, 5.2% glucose, 0.7% yeast (w/v) and 8.88% grape or clementine juice (v/v)), eggs were collected and dechorionated for 3 m in 10% bleach, 1 m sterile dH_2_O, 3 m further in 10% bleach, 1 m in 100% ethanol, 1 m sterile dH_2_O, before resuspension in sterile 1X PBS. 20 µL of egg suspension was pipetted into T75 cell culture flasks with filter caps containing 60 mL SYA. To generate gnotobiotes, OD_600_ of 48h bacterial cultures was measured and normalised to OD_600_ = 1, washed and resuspended in sterile 1X PBS, diluted 1:5 (OD_600_ = 0.2), and 200µL aseptically added to each flask containing sterilized eggs. To generate fly cultures in glass *Drosophila* vials, 8 mL of fly food was used, 5 µL of eggs were deposited, and 50 µL of OD_600_ = 0.2 bacterial cultures were added. All flies were transferred to fresh sterile media upon adult emergence, 10 days after egg deposition. Flies were developed on holidic diet for viability experiments where specified. Where adult-onset dietary manipulations were performed, flies were transferred to experimental media days 3-10 adulthood, after preceding culture on SYA.

Holidic *Drosophila* medium (mol FlyAA formulation) ^34^ was used, 8 mL/vial, with 5% sucrose. For acetate free medium, acetate buffer was replaced with 0.2M Phosphate buffer (16.44 g/L KH_2_PO_4_ and 8.95 g/L Na_2_HPO_4_) pH 6.5 was used from 10X stock. To determine acetate requirement, axenic cultures were established in vials containing 8 mL acetate- or phosphate-buffered holidic medium. Numbers of emerging flies were counted/weighed on day 21. To determine bacterial vitamin provision (*A. pomorum, E. coli, B, subtilis*), cultures were established on 8 mL of holidic diet either containing all B-vitamins or with pantothenate/riboflavin singly omitted, then counted on day 21.

For inducible SCFA production, four holidic diets were made with cholesterol dissolved in water, ethanol, butanol, or isobutanol. Once media solidified, each respective alcohol (1% v/v) was further added to the surface. Diets were then either left alone or conditioned by *A. pomorum* (OD_600_ = 0.2) for 48h to produce SCFAs from the alcohols. Axenic females developed on SYA before transfer to these diets for days 3-10 adulthood (15/vial), with transfer to freshly-prepared and conditioned media every 2-3 days.

When antibiotics were incorporated into medium (Figure S3) 3-day old axenic/conventionally-reared flies were fed (1) SYA, (2) SYA conditioned 48h by 20 conventional males to generate conventionalized flies, or (3) SYA containing 500 μg/ml ampicillin, 50 μg/ml tetracycline, and 200 μg/ml rifamycin ^64^, prepared from a 100× stock in 50% ethanol.

To condition food, 40 male flies (either axenic or *A. pomorum* colonized) were added to sterile SYA (∼3mm deep to maximise bacterial penetrance into total volume) for 48 hrs. The media were then autoclaved for 15-minutes and allowed to cool and re-set without disturbance. Females (n=15) were then added for days 3-10 adulthood, with transfer to freshly-prepared media every 2-3 days.

To assess impact of dead *A. pomorum*, cultures were grown as normal, split, and half were autoclaved. From days 3-10 adulthood, flies reared axenically were fed either sterile SYA+50µL sterile PBS, sterile SYA inoculated with *A. pomorum* (50µL/5mL medium), or sterile SYA with added autoclaved *A. pomorum* (50µL/5mL medium). Bacteria (live or autoclaved) were added 2 days before flies, to condition the food. Flies were transferred to new media every 2-3 days, then assayed for TAG.

To assess impact of vitamin supplementation, axenics were fed diets supplemented with vitamin supplements ^36^ from days 3-10 adulthood before TAG assay. 1X in mg/L: B_1_ = 1.4, B_2_ = 0.7, B_3_ = 8.4, B_5_ = 10.8, B_6_ = 1.7, B_7_ = 0.1, B_9_ = 0.9; 2X and 3X multiplied accordingly. The B_5_ -only condition was fed at the same concentrations with all other vitamins omitted from the supplement.

To assess impact of varied carbohydrates, media were prepared containing sucrose, glucose, fructose, no added carbohydrate, or gluconate, as specified in figure legends. Yeast-agar media were prepared to 80% final volume then sterile 5x carbohydrate solutions were added after autoclaving. Flies developed on SYA before transfer to these media for days 3-10 adulthood, followed by TAG or LC-MS analysis.

To assess bacterial impact on TAG mobilisation, flies developed on SYA until 7 days adulthood. TAG assays were conducted to establish baselines per condition, then flies were allocated either to obesogenic (SYA containing 20% Sucrose w/v) or starvation diet (1.5% Agarose w/v), followed by daily TAG assays (n=3/sample), up to 5 days. Survival on the starvation diet was recorded thrice daily. Flies were flipped onto fresh sterile food every 2/3 days.

For *Akh* knockdown*, Akh-Gal4* (gift from K. Halberg, University of Copenhagen) was backcrossed into *w^Dah^*, and virgins were crossed to either w^Dah^ wild-type or *UAS-Akh^RNAi^*(VDRC stock 113271). Flies were made axenic/gnotobiotic as above. Flies developed on SYA, then 7 day-old adults were flipped onto 1.5% agarose for starvation resistance assay.

For PANK knockdown, *Da-Gal4* (gift from C. Slack, University of Warwick) were backcrossed into *w^Dah^*, and virgins were crossed either to wDah wild-type or *UAS-PANK^RNAi^* (VDRC stock 109437). Flies were made axenic/gnotobiotic as above. CFUs were quantified by homogenisation with a pestle in 500 µL PBS in a 1.5 mL microfuge tube, diluting 1:10, 1:100 and 1:1000, and spiral-plating on YPD agar.

In all experiments, media surfaces were routinely plated on bacterial growth media (1% agar) to ensure sterility, and all experiments where any contamination was detected in any condition were discarded. In all experiments where flies were maintained beyond day 3 of adulthood, mated females were allocated to sterile glass vials (n=15/vial).

To prepare specific tissues flies were either dissected with foreceps on ice after brief CO2 anaesthesia (midgut, fat body by taking abdomens once intestinal/renal/reproductive anatomy was removed); or to collect heads/thoraces/abdomen flies were frozen at −80c for 48 before separation on a series of cold metal sieves (collecting each sequentially smaller body part).

### Bacterial culture

*Levilactobacillus brevis* DmCS003 and *Lactiplantibacillus plantarum* DmCS001 were grown and maintained in YPD (1% Yeast Extract, 2% Peptone, 2% Dextrose (glucose)) medium at 30°C without shaking, while *Acetobacter pomorum* DmCS004, *Acetobacter pomorum* WJL, *Acetobacter fabarum* DsW_054, and *Acetobacter tropicalis* DmCS006 were grown and maintained in M9L medium (5X M9 Salts: 3.9% Na_2_HPO_4_ anhydrous, 1.5% KH_2_PO_4_ anhydrous, 0.25% NaCl, and 0.5% NH_4_Cl all w/v, 0.002M MgSO_4_ and 0.001M CaCl_2_) with 0.5% DL-lactic acid (M9L) at 30°C with shaking at 250 rpm. Δ*PQQ-adhI* mutants *A. pomorum and A. fabarum* strain (transposon insertion, ^65^) were grown in M9L with 50 µg/mL kanamycin at 30°C with shaking at 250 rpm. The *A. fabarum* purine salvage mutants were grown in M9L (Δ*xdhB*) or M9L with 15 µg/mL tetracycline (Δ*urhA::Tn7* and Δ*urhA::Tn7_allP). Escherichia coli* BW25113 was grown in Lysogeny Broth (LB) at 37°C with shaking at 250 rpm, while the mutant strain JW0129 (*panC*750(del)::kan) was grown in LB with 50 µg/mL kanamycin under the same conditions as the wild-type strain. *B. subtilis* 168 and *B. subtilis* Δ*panC* (GP4361 Δ*panC*::*cmr* from ^48^) were grown in YPD and YPD with 5 µg/mL chloramphenicol, respectively, at 30°C with shaking at 250 rpm.

### Tn7 bacterial strain construction

We built a yeast - *E. coli* shuttle vector by introducing a CEN-URA cassette into pUC18T-mini-Tn7T-Tet ^66^ via recombination cloning ^67^. *Nde*I-linearized plasmid was used to transform yeast with a CEN-URA PCR product amplified from pMQ30 ^67^ with primers: 5’-catcagagcagattgtactgagagtgcaccatctctaatttgtgagtttagtatacatgc-3’, and 5’-attttctccttacgcatctgtgcggtatttcacaccgcatgaaagggcctcgtgatacgc-3’. The resulting plasmid, pOSW2, was cut with *Eco*32I to add the *allP* gene within the mini-Tn7 transposon. The linearized plasmid was used to transform yeast with *allP* PCR product amplified from pCM62-allP ^42^with primers: 5’-aattcctcgagaagcttgggcccggtacccagctggcacgacaggtttcc-3’, and 5’-ggttggcctgcaaggccttcgcgaggtaccgccattcgccattcaggctgc-3’. The resulting plasmid, pOSW2_Tn7allP, contains the *allP* gene under the control of a constitutive P*^lac^* promoter within the mini-Tn7 transposon. Triparental mating was used to introduce the completed Tn7 construct into the *A. fabarum* genome, with *E. coli* S17 strains donating pOSW2_Tn7allP and helper plasmid pUX-BF13 ^68^. Transconjugates were selected for on YPD with 20 µg/ml of tetracycline and 0.2% acetic acid. After isolation, insertion of Tn7 downstream of the *glmS* gene in *A. fabarum* was confirmed via PCR and sequencing across the transposon junction with primers: 5’-CTGGCAAAATCGGTAACGGTGG-3’ and 5’-GATACCGTCGACCTCGAACC-3’. In parallel, control strains were generated with pOSW2 to introduce Tn7 without *allP*.

### TAG assay

3-day-old adult females (for Acetobacteraceae vs Lactobacillaceae, *PQQ-adhI* mutant experiments, *E. coli*, and *B. subtilis* experiments) unless stated otherwise, were weighed in groups of 5 (unless otherwise stated) and collected in 2mL screw cap tubes with 125 µL of TEt Buffer (TE buffer with 0.1% triton X-100) and 1.4 mm ceramic beads. For adult-onset manipulations 10-day-old adults were used. Flies were homogenized in a Bead Ruptor Elite bead mill homogenizer at (6.5 metres/second, 30 seconds), incubated at 72 °C for 15 m, spun for 5 m at 4°C at 12000xg. 3µL of supernatant was used when using the Infinity^TM^ Triglycerides reagent (Thermo Scientific), following the manufacturers protocol. After Infinity reagents ceased to be sold in the UK, 2µL homogenate was used with reagents made in-house ^69^. Standard curves were generated 1 mg/mL to 0 mg/mL. TAG levels were normalized to weight of the 5 flies.

### RNA isolation and sequencing

RNA was extracted from 3-day old adult axenic, *L. brevis*, and *A. pomorum* female gnotobiotes as per ^70^. PolyA transcripts were enriched with Perkin Elmer NEXTFLEX beads. 3’ Sequencing was performed by the University of Glasgow MVLS Shared Research Facility using LexoGen QuantSeq library prep and an Illumina NextSeq, and analysed in Searchlight ^71^.

### LC-MS sample preparation

Nutrition was profiled by scraping media from the surface 10 days after bacterial/fly inoculation, using aseptic technique and avoiding larvae. Unconditioned SYA was incubated without bacteria or flies for 10 days. Media was collected into pre-weighed 2 mL screw cap tubes with 1.4 mm ceramic bead. The corresponding flies were subsequently CO_2_-anaesthetized, quickly weighed (in groups of 5), and added to chilled homogenisation tubes. Cold MCW (3 methanol: 1 chloroform: 1 water) was added at 120 µL per mg sample, before 30 s homogenization with a Bead Ruptor Elite bead mill homogenizer at speed 6.5 m/s, centrifugation (12,000 rpm, 5m, 4°C). Supernatants were then frozen at −80 before LC-MS analysis. For sex-specific analysis, males and females were processed as above in groups of 5, on day 3 of adulthood.

### Sample preparation for SCFA quantification (GC-FID)

Flies and media were collected as for LC-MS (except axenics on defined diet ±acetate, which were collected 21 days after egg deposition). Media samples were weighed, 1M NaOH added (100 µL per 100 mg), and frozen at −80°C. Flies were subsequently weighed to >100 mg and NaOH added. Samples were freeze dried ≥36 h then ∼100 mg in a fresh tube was homogenized with a sterile pestle in H_2_O (3:1), and then used directly in the ether extraction. Bacterial cultures were grown from single colonies for 48 h, pelleted and washed in sterile 1X PBS, then resuspended OD_600_ = 1 (1 mL). From these cultures, fresh media were inoculated (OD_600_ = 0.025) and grown for either 48 (YPD) or 72 hours (*Drosophila* defined media). 800 µL supernatant was then mixed 1:1 with 1M NaOH and frozen at −80°C. Sterile media were used as controls.

External standard controls, freeze dried samples, and bacterial culture samples were extracted in similar fashions. Briefly, the samples were brought to a volume of 800 µL with dH_2_O, and given 100 µL of internal standard (2-ethyl butyric acid in 2 M NaOH; 73.8 mM) and 100 µL of orthophosphoric acid. Samples were vortexed to mix. 1.5 mL of diethyl ether was added to the freeze-dried samples, while 3 mL of diethyl ether was added to the bacterial culture samples. Tubes were vortexed for 60 s at 1500 rpm, and the top ether layer was transferred to a new 15 mL falcon tube. This step was repeated 3x. The samples were then transferred to gas-tight vials, and sealed with a crimp, ready for gas chromatography. The SCFAs were analysed on a gas chromatograph equipped with a flame ionization detector (FID) (Agilent 7820A; Agilent Technologies, Stockport, USA). The oven was programmed at an initial temperature of 80 °C for an initial 1 min, increasing 15 °C/min, thereafter increasing to 210 °C; using a DB–WAX UI column (15 m × 0.53 mm id × 1 μm-film thickness, Agilent). The carrier gas (nitrogen) was set at a flow rate of 30 mL/min. The ratio of each SCFA was compared to the internal standard to calculate concentrations. Individual SCFAs were calibrated against external standards dissolved in 2M NaOH (185.825 mM acetate, 144.447 mM propionate, 114.189 mM butyrate, 97.3056 mM isobutyrate, 83.4348 mM valerate, 87.0342 mM isovalerate, 76.522 mM hexanoate, 52.6385 mM isohexanoate, 65.7951 mM heptanoate, and 53.178 mM octanoate).

### Acyl-CoA analysis

Flies were aseptically anesthetized on ice collected individually into sterile microcentrifuge tubes containing 500 µL of ice cold 10% (w/v) trichloroacetic acid, briefly centrifuged at 4°C to submerge flies, and flash frozen in liquid N_2_.

Acyl-CoAs were measured by liquid chromatography-high resolution mass spectrometry as previously described ^72^. Briefly, samples were quenched in 1 mL 10% (w/v) trichloroacetic acid in water for extraction. Samples were homogenized by using a probe tip sonicator in 0.5 second pulses 30 times then centrifuged at 17,000 x g for 10 min at 4°C. Supernatant was purified by solid phase extraction (SPE) cartridges (Oasis HLB 10 mg, Waters) that were conditioned with 1 mL of methanol and 1 mL of water. Acid-extracted supernatants were loaded onto the cartridges and washed with 1 mL of water. Acyl-CoAs were eluted with 1 mL of 25 mM ammonium acetate in methanol and evaporated to dryness under nitrogen. Samples were resuspended in 50 µL of 5% (w/v) 5-Sulfosalicyilic acid and 20 µL injections were analysed on an Ultimate 3000 UHPLC using a Waters HSS T3 2.1×100mm 3.5 µm column coupled to a Q Exactive Plus. The analysts were blinded to sample identity during processing and quantification. Data were analysed using Tracefinder 5.1 (Thermo).

### LC-MS metabolomics

We performed hydrophilic interaction liquid chromatography (HILIC)-mass spectrometry (LC-MS) on an UltiMate 3000 RSLC (Thermo Fisher, San Jose, CA, USA), with 150 × 4.6 mm ZIC-pHILIC column (Merck SeQuant, Umea, Sweden) (at 300 µl/min), and with Orbitrap Exactive detection (Thermo Fisher, San Jose, CA, USA). We used 50,000 resolving power in positive/negative switching mode, electrospray ionisation voltage at 4.5 kV in positive and 3 kV in negative modes. Buffers were A: 20 mM ammonium carbonate in H_2_O and B: Merck SeQuant: acetonitrile. The gradient ran from 20% A: 80% B to 80% A: 20% B in 900 s, followed by a wash at 95% A: 5% B for 180 s, and equilibration at 20% A: 80% B for 300 s.

### Bioinformatics and data analysis

Raw LC-MS files were converted into the mzXML open format using MSConvert from the proteowizard pipeline. During conversion, the *m*/*z* data was centroided. These files were uploaded to the Glasgow Polyomics Integrated Metabolomics Pipeline (IMP) ^73^ and processed using the embedded pipeline. Briefly, chromatographic peaks were extracted from the mzXML files using the centwave detection algorithm, replicates were aligned and combined, then all peaks that were not reproducibly detected within groups when subjected to intensity filtering and noise filtering were removed. The data set was then gap-filled to produce a set of reproducible peaks. Identifications were based on the Metabolomics Standards Initiative proposed minimum reporting standards. The inbuilt pipeline was used to determine differential metabolite abundance between experimental groups, or with LIMMA (in R) where specified.

MINT and multi-sample PCA were performed with the R MixOmics library, FELLA with the FELLA library ^74^. Figures were produced in ggplot2 with custom scripts. All statistical modelling was performed in R, with post-hoc tests from the emmeans library. tSNE was performed with the Rtsne library. Survival was analysed with the survival library, MOFA was performed with the MOFA2 library.

### Genome-scale metabolic modelling and multi-omics integration

Fruitfly-GEM ^75^ was used as the reference *Drosophila* genome-scale metabolic reconstruction. Whole-fly RNA-seq profiles from Ax (n=6), Ap (n=5) and Lb (n=6) biological replicates were mapped to model genes and used to generate 17 replicate-specific transcript-informed metabolic models. In total, 1,528 RNA-seq genes mapped to Fruitfly-GEM, providing expression evidence for 1,528/1,738 model genes and 5,704/6,030 gene–protein–reaction(GPR)-linked reactions. Gene expression was propagated through GPR rules to derive replicate-specific reaction-expression scores and scale reaction capacities while retaining the native bounds of reactions lacking expression evidence.

Because the study focused on adult flies, ATP maintenance/hydrolysis was used as the primary model objective rather than biomass production. All biological conditions were simulated under the same primary SYA-like medium to ensure that between-condition differences arose from transcriptomic constraints rather than different nutrient assumptions. Additional simulations were performed under FLYAA ^34^ and the experimental YA, GYA and GnYA dietary perturbations. Parsimonious flux balance analysis (pFBA) was used to estimate representative optimal flux states, and targeted flux variability analysis (FVA) was used to quantify feasible minimum, maximum, midpoint and interval-width fluxes. Predicted metabolite turnover and exchange fluxes were also derived from the replicate-specific models. These quantities were treated as model predictions and not as measured metabolic fluxes or metabolite concentrations.

Quantitative LC-MS metabolomics from the same study was analysed as an independent experimental evidence layer for fly and conditioned-food samples. Peak-level *Ap*-vs-Ax and *Lb*-vs-Ax effects were assessed with multiple-testing correction, and metabolite annotations were mapped to Fruitfly-GEM using KEGG identifiers or normalized metabolite names. Of 5,880 measured peaks, 1,035 mapped to GEM metabolites, including all 29 prespecified targeted peaks. These mappings were used to link measured metabolite changes to adjacent GEM reactions and their associated RNA/GPR evidence, without treating metabolite abundance changes as direct flux measurements.

For statistical comparison of model-derived states, features were standardized and condition-level separation was tested using PERMANOVA, with PERMDISP used to assess dispersion effects. PCA was used for visualization. Pairwise *Ap*-vs-Ax and *Lb*-vs-Ax tests used exact assignment/permutation inference where available, followed by Benjamini–Hochberg FDR correction. Analyses included mapped metabolic-gene expression, GPR/reaction-expression scores, predicted pFBA flux, metabolite turnover, exchange flux, subsystem-level reaction-expression states and targeted FVA metrics.

### *In vitro* bacterial nitrogen rescue assay

3-day old flies were transferred for 10 h to sterile TC 75 flasks containing 1.5% agarose and 2.5% brilliant blue (Proquimac) (w/v). During this time period, guts and frass became blue, indicating gut clearance and elimination of nitrogenous carryover from food (yeast). Flies were then transferred to sterile T75 flasks, without agarose or food, for 24 hrs defecation (confirming all frass spots were blue), from which frass deposits were resuspended in 15 mL of N-free M9L media (no ammonium), and filter sterilized (0.22 µm pore filter). To normalise solute density, OD_630_ (“blueness”) was quantified and samples were normalised to equivalent OD (lowest common denominator) in N-free media. N-free media were also mixed with the blue agarose, as a negative control to ensure the agarose and blue dye were not supplying nitrogen. Regular M9L was used to wash sterile blue agarose for a positive control to ensure the dye did not hinder growth. These media were then inoculated with *A. pomorum* or *A. fabarum* (starting OD_600_ = 0.025), and incubated for 72 hours at 30°C, shaking 250 rpm. OD_600_ measurements were taken, samples were serially diluted, and plated by sterile spreader beads to determine CFUs. *B. subtilis* CFUs were plated with a Don Whitley spiral plater.

### *Acetobacter pomorum* carbon/energy utilization

To test *A. pomorum* carbon use, 2-day old culture grown in M9L (OD_600_ = 1, washed 3X and resuspended to 1ml in sterile PBS) was subcultured into M9 with 0.5% lactate, glucose, fructose, gluconate, or ethanol, all pH5.5. 3 mL cultures were inoculated to a starting OD_600_ = 0.025. After 72 hrs, OD_600_ were measured to determine growth.

To test *A. pomorum* energy use via partial oxidation, M9L pH 5.5 with 0.025% pH indicator (congo red) were supplemented with either 1% glucose, fructose, gluconate, or ethanol. 5mL of media was inoculated as per carbon utilization testing, and after 72 hrs pH of filtered supernatant was measured on a Mettler Toledo SevenEasy pH meter.

### Acetate assay

*A. pomorum* and *L. brevis* were grown for 48 hrs in 3 mL MRS medium at 30°C with *A. pomorum* cultures shaking at 250 rpm. Samples were spun, supernatant filter sterilized, and processed for the acetate assay, with uninoculated MRS as a control. Acetate was measured using a colorimetric assay kit (MAK605 Sigma Aldrich), following the manufacturer’s protocol and a standard curve.

### 2-NBDG uptake assay

Glucose uptake into muscle was measured using 2-NBDG (2-[N-(7-nitrobenz-2-oxa-1,3-diazol-4-yl)amino]-2-deoxyglucose), a fluorescent glucose analogue that is transported into cells but only minimally metabolized thereafter. Five-day-old females with or without *A. pomorum*, were collected at lights on (before the morning meal) and injected in the thorax with 100 nL of either 5 mM 2-NBDG in PBS or PBS alone. The 2-NBDG injection solution was prepared from a 50 mM stock in 100% ethanol and diluted in PBS to a final concentration of 5 mM, corresponding to an estimated final haemolymph concentration of 0.5 mM 2-NBDG. Following injection, flies were kept on 1% agar for 30 min to allow glucose uptake. Legs were then dissected, mounted in Schneider’s medium, and imaged immediately. 2-NBDG fluorescence was imaged by confocal microscopy (Zeiss LSM 900, 20× objective, 1 µm z-spacing). All imaging settings were kept identical across samples. Z-stacks were maximum projected in ImageJ, and mean fluorescence intensity was quantified after background subtraction from representative regions within each sample.

## Acknowledgments

This work was funded by a UKRI Future Leaders Fellowship (MR/S033939/1, MR/Y019660/1) and University of Glasgow funds (LKAS Fellowship and Reinvigorating Research) to AJD; and BBSRC funds (BB/W510658/1) to AJD and DRS. AM and KR were supported by a Novo Nordisk Foundation grant (NNF24OC0092524) to KR. We thank Nicolas Buchon and Richard Cogdell for comments on drafts of the manuscript, Clement Regnault for running mass spectrometry instrumentation, and Won-Jae Lee and Jörg Stülke for bacterial strains.

## Supplemental note 1

### LC-MS chemical profiling indicates microbe-specific metabolomic impacts of *A. pomorum* and *L. brevis*

To begin a systematic characterization of how defined, minimal microbiota impact host chemical composition, we turned to untargeted global metabolomic approaches. We used an established Liquid-Chromatography-Mass Spectrometry (LC-MS) methodology ^76^, capable of resolving thousands of compounds, among which fragmentation data for some peaks plus comparison to a set of >100 definitive standards enabled identification of a subset of compounds. We first performed a small-scale pilot (n=3 pools of 5 whole females per microbial condition), to confirm reproducible metabolite extraction and sensitivity to microbiota manipulation (Figure S4). In both positive- and negative-ion mode, chromatograms were reproducible among conditions (Figure S4A), indicating repeatable extraction. We identified 4,784 peaks, among which 528 generated fragmentation data, and 109 could be identified by comparison to standards. We quantified intensities for all peaks and performed Principal Components Analysis (PCA) for an aggregate view of among-condition differences (Figure S4B). PCA suggested that metabolic state of *Ap-*flies was robustly differentiated from axenics. By contrast, differences between *Lb-*flies and axenics were negligible. Between axenics and *Ap-*flies, 185 peaks were differentially abundant at FDR≤0.05 (Figure S4C), of which 179 were annotated or could be identified from fragmentation or by comparison to standards. Between axenics and *Lb*-flies, only three peaks were differentially abundant (Figure S4C). Of the peaks that were differentially abundant either in *Ap-*flies or *Lb-*flies, 44 were identified by comparison to standards or from fragmentation data (Figure S4D – noting that some compounds are identified more than once, e.g. when detected in both positive and negative ion mode).

### Negligible sexual dimorphism in metabolomic effect of microbiota

Most studies of the fly microbiota to date have been performed in one sex only, but males and females can differ in their response to metabolic alterations. We characterised how both sexes responded to our axenic/*L. brevis*/*A. pomorum* paradigm, allowing us to partition variation between microbiota, sex, sex-specific impacts of the microbiota, and sexual dimorphism arising only under specific microbiota conditions.

We detected 3,244 peaks in total. PCA separated samples by sex on the first PC, and by microbiota on the second PC, confirming that we could detect effects of each factor, but that their effects might be orthogonal, without substantial sex-specific effects (Figure S5A). Partitioning variance with PERMANOVA showed that sex was the biggest source of variance (r^2^=0.72, p=0.001), and microbiota also had a significant effect (r^2^=0.05, p=0.04), but no sex-by-microbiota interaction was detected (p=0.3) (Figure S5B).

For a peak-by-peak view, we performed two complementary analyses to test differential peak abundance (FDR≤0.05), and identified set overlaps using UpSet. First, we tested whether microbiota altered metabolic sexual dimorphism. Within each microbial condition, we identified sets of sexually dimorphic peaks, the union of which comprised 1727 peaks (Figure S5C). In 881/1727 peaks, dimorphism was microbiota-independent. The remaining 846 peaks were distributed among comparatively small subset: for example, 138 peaks were elevated in females uniquely in *Ap-*flies. However, there was no substantial overall pattern in which sexual dimorphism was released or repressed in one microbiota condition, consistent with the prior PCA and PERMANOVA. Thus, sexual dimorphism does not appear to be substantially released or repressed by microbiota variation.

Next, we tested whether sex altered metabolic response to microbiota. Within each sex, we identified peaks that responded to the microbiota. We analysed impacts of *A. pomorum* and *L. brevis* separately (Figure S5D-E). *A. pomorum* modulated 478 peaks, 66% of which were altered in females, but not males (177+137=314 total). The next-largest set comprised 93 peaks that were upregulated by *A. pomorum* in both sexes, complemented by 16 that were downregulated in both sexes. Only 11% of *A. pomorum*-regulated peaks (31+22=53) responded uniquely in males. Thus, while more effects of *A. pomorum* were detected in females, the effects in males were predominantly a subset of what was observed in females. More sex-specificity was apparent in response to *L. brevis*, where 89 differentially abundant peaks were detected in both sexes. Of these peaks, 21% (18+1=19) showed concordant responses between females and males, but the rest showed sex-specific responses. In summary, these analyses suggested that assaying females was sufficient to capture impacts of *A. pomorum* on both sexes, while the effect of *L. brevis* was perhaps more sex-specific but also comparatively modest.

### The female response to *A. pomorum* encapsulates most effects observed in males and/or in response to *L. brevis*

Given the signal that females appeared to encapsulate the male response to microbiota, we examined the female response to microbiota specifically (Figure S6). We found that 61% of peaks impacted by *L. brevis* were also impacted by *A. pomorum* (38/62), however this overlap accounted for only 9% of *A. pomorum*’s overall effect (38/425), suggesting that *A. pomorum* exerts a greater overall impact and a majority of effects of *L. brevis* are encapsulated by *A. pomorum* (Figure S6A). Effects of *A. pomorum* were also quantitatively greater than those of *L. brevis* (Figure S6B-C). For peaks impacted by both bacteria, direction of impact was correlated, but magnitude was generally greater in response to *A. pomorum* than *L. brevis* (Figure S6C). Of the peaks for which we could derive a definitive metabolite identification from fragmentation or standards, 44 were differentially abundant in *Ap-*flies or *Lb-*flies (Figure S6D). Only one of these metabolites (L-citrulline) was differentially abundant (increased) in *Lb-*flies, but it also increased in *Ap-*flies and to a greater extent. *A. pomorum* thus appeared to have a greater metabolomic impact that *L. brevis*, consistent with TAG data (Figure 1A).

## Supplemental note 2

If *A. pomorum* disinhibit carbohydrate metabolism, we expected altered kinetics of storing energy as TAG upon carbohydrate excess, and mobilising TAG upon energy deficit. On high-sucrose diet, *Ap*-fly TAG rose smoothly over four days, while axenic TAG stayed flat until a sudden rise at 72h (Figure S13A), consistent with impaired metabolic control. After energy deficit was induced by starvation, axenics lost TAG faster over the first 48h, suggesting TAG mobilisation is not intrinsically impaired. However axenic TAG then plateaued above *Ap*-flies (Figure S13B), suggesting a deficiency for mobilising final reserves *in toto*. Interestingly, this corresponded to greater starvation resistance (Figure S13C), suggesting use of alternative energy sources once axenic TAG mobilisation becomes recalcitrant. This might indicate that final TAG depletion portended death. Consistent with this hypothesis, impairing TAG mobilisation constitutively in both microbial conditions, with RNAi against *Akh* (the fly glucagon homologue) rendered both *Ap*-flies and axenics constitutively starvation-resistant (Figure S13D). This suggested that starvation sensitivity in *Ap*-flies is due to reduced TAG storage, revealing a stress-resistance cost of symbiosis.

## Supplemental note 3

To determine whether gut microbiota reshape not only host gene expression but the metabolic network architecture and functional output it supports, we constructed replicate-specific, transcript-constrained genome-scale metabolic models for axenic (Ax, n=6), *Acetobacter pomorum*-associated (Ap, n=5), and *Levilactobacillus brevis*-associated (Lb, n=6) flies using Fruitfly-GEM as the reference reconstruction. Of 12,931 RNA-seq genes analysed, 1,528 (11.8%) mapped to Fruitfly-GEM, providing expression evidence for 87.9% of model genes and informing 5,704/6,030 (94.6%) GPR-linked reactions (Figure S21A). Expression was propagated through gene–protein–reaction rules to generate replicate-specific reaction-expression scores, and parsimonious flux balance analysis (pFBA) and flux variability analysis (FVA) were used to derive predicted optimal and feasible flux states under a fixed ATP-maintenance objective and matched dietary conditions across all three groups, isolating microbiota-associated effects from confounding nutritional variation.

At the transcript level, mapped metabolic-gene states separated significantly by condition (global PERMANOVA R²=0.240, p=0.017; PERMDISP p=0.720, excluding a dispersion-driven artifact), with both *Ap*-vs-Ax and *Lb*-vs-Ax comparisons individually significant (FDR=0.030 and 0.032, respectively; Figure S20A). This condition-associated structure was not only preserved but modestly amplified after propagation through the GPR layer: reaction-expression states showed stronger separation among conditions (R²=0.273, p=0.010; PERMDISP p=0.753), again with significant pairwise effects for both *Ap*-vs-Ax (FDR=0.022) and *Lb*-vs-Ax (FDR=0.030) (Figure S20B). Subsystem-resolved analysis of the *Ap*-vs-Ax reaction-expression signal identified ten metabolic subsystems surviving FDR≤0.05, spanning lipid-handling pathways (formation and hydrolysis of cholesterol esters, FDR=0.033, n=118; acyl-CoA hydrolysis, FDR=0.033, n=24; cholesterol metabolism, FDR=0.033, n=28; glycosphingolipid and sphingolipid metabolism, FDR=0.033 and 0.047, n=8 and 56), carbohydrate metabolism (starch and sucrose metabolism, FDR=0.047, n=27; chondroitin sulfate degradation, FDR=0.033, n=34), and nucleotide/cofactor metabolism (purine metabolism, FDR=0.033, n=51; folate metabolism, FDR=0.033, n=24) (Figure S20C). Effect sizes across these subsystems were substantial, ranging from R²≈0.31 to R²≈0.77, indicating that colonisation-associated transcriptional change is concentrated in specific, biologically coherent metabolic modules rather than distributed diffusely across the network.

This signal collapsed sharply upon propagation to predicted flux (Figure S20D). Global effect size fell from R²=0.273 at the GPR/reaction-expression layer to R²=0.047 at the pFBA layer, with the latter far from significance (p=0.933; PERMDISP p=0.668); only 13 of 11,829 reactions varied above the configured numerical tolerance, and none remained differential after FDR correction (Figure S21B). Predicted metabolite turnover showed a comparably null result (R²=0.050, p=0.918), and predicted exchange flux contained no variable features whatsoever (R²=0.000, p=1.000). Targeted FVA likewise identified no reaction for which the Ax, *Ap*, and *Lb* feasible flux intervals were distinguishable after correction, indicating that the absence of a flux-level phenotype was not an artifact of collapsing the feasible solution space to a single optimum via pFBA.

To test whether this attenuation reflected a fragile choice of medium or maintenance demand, we repeated the robustness analysis of model constraint engagement across five simulated media (SYA, FLYAA, YA, GYA, GnYA) at multiple ATPM fractions. Median active-bound pressure tracked medium and ATPM stringency as expected — rising sharply at 75% ATPM demand under both SYA and FLYAA and dropping to a shared minimum under FLYAA at 25% ATPM — but condition (Ax/*Ap*/*Lb*) trajectories were closely superimposed across all nine scenarios (Figure S21C), with no scenario producing detectable condition-specific separation. Maximum simulated ATP-maintenance capacity varied substantially by diet (132.5 under SYA and GYA, 162.5 under FLYAA, 92.5 under YA and GnYA) but was numerically identical across Ax, *Ap*, and *Lb* within every medium tested (Figure S21D), confirming that the flux-level null result is a property of the modelled metabolic network under matched nutritional input rather than an artifact of one simulation setting.

Together, these results indicate that colonisation drives significant, subsystem-concentrated restructuring of the transcript-informed metabolic-gene and reaction-expression states, but this restructuring does not translate into a distinguishable predicted optimal or feasible flux phenotype under standard dietary conditions and a fixed ATP-maintenance objective. This pattern is consistent with a model in which gut bacteria reconfigure the host’s metabolic gene-expression architecture, potentially altering regulatory capacity or readiness to respond to future perturbation, without a corresponding shift in baseline flux-level metabolic output, at least as captured by the objective and constraints tested here. This suggests that flux alteration is likely mediated by transcription-independent mechanisms

## Supplementary figures

**Figure S1.**
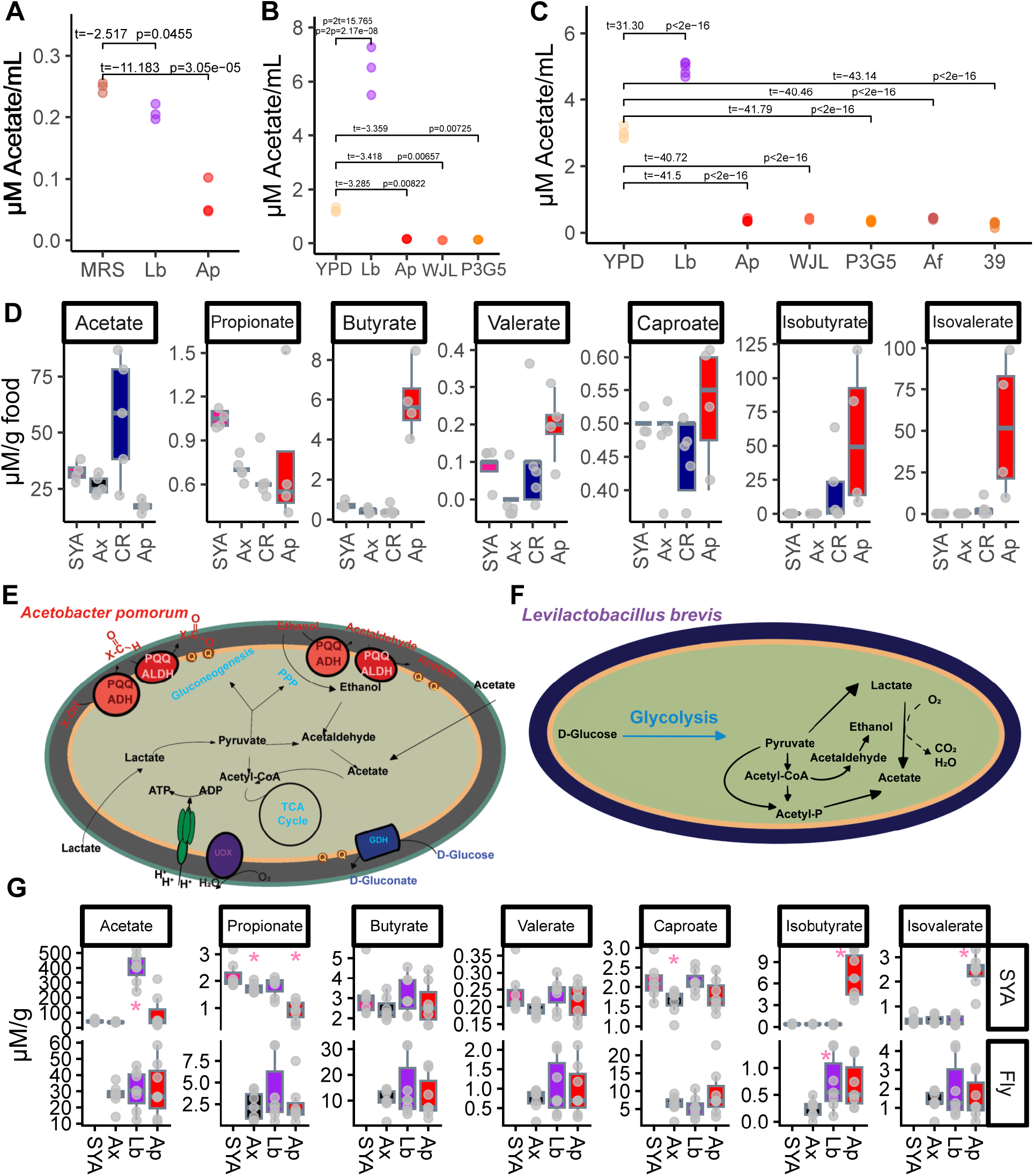
SCFA metabolism of fly associated bacteria in bacterial media, SYA food, and flies. **A.** *A. pomorum* is a net acetate user in MRS medium as measured by acetate assay, with significantly less acetate in *A. pomorum* culture supernatant after 48 hrs. **B.** GC measurements of acetate in YPD media cultured with different *A. pomorum* strains including the *ΔPQQ-adhI* mutant (P3G5) and *L. brevis*. *L. brevis* increased acetate levels, while all *A. pomorum* strains reduced levels after 48 hrs of culture. **C.** GC measurements of acetate in YPD media cultured with different *A. pomorum* and *A. fabarum* strains including Δ*PQQ-adhI* mutants (*A. pomorum* – P3G5, *A. fabarum* – 39) and *L. brevis*. *L. brevis* increased acetate levels, while all *A. pomorum* and *A. fabarum* strains reduced levels after 48 hrs of culture. **D.** GC analysis of food colonized by conventionally reared (CR), axenic (Ax) and *A. pomorum* associated flies (Ap) shows that A. pomorum does not recapitulate CR profiles, as it does not increase acetate levels. **E.** metabolic map of *A. pomorum* depicts how it conserves energy. Alcohols (X-OH) are converted to acylaldehydes via the pyrroloquinoline quinone-dependent alcohol dehydrogenase (PQQ-AdhI) and subsequently to SCFAs via the PQQ-dependent aldehyde dehydrogenase (PQQ-Aldh) in the inner membrane, thus SCFA production is extracellular. Glucose is oxidized to D-gluconate via a membrane bound glucose dehydrogenase. **F.** Metabolic map of *L. brevis*. *L. brevis* can generate acetate via acetyl-CoA or via lactate in the presence of oxygen via lactate oxidase. **G.** GC analysis of uncolonized and colonized SYA food and the flies that developed in the food. SYA food colonized with Lb gnotobiotic flies has increased levels of acetate, while *A. pomorum* fly colonized food has increased levels of isobutyrate and isovalerate. Axenic, *L. brevis*, and *A. pomorum* flies maintain similar SCFA levels despite differences in the food.

**Figure S2.**
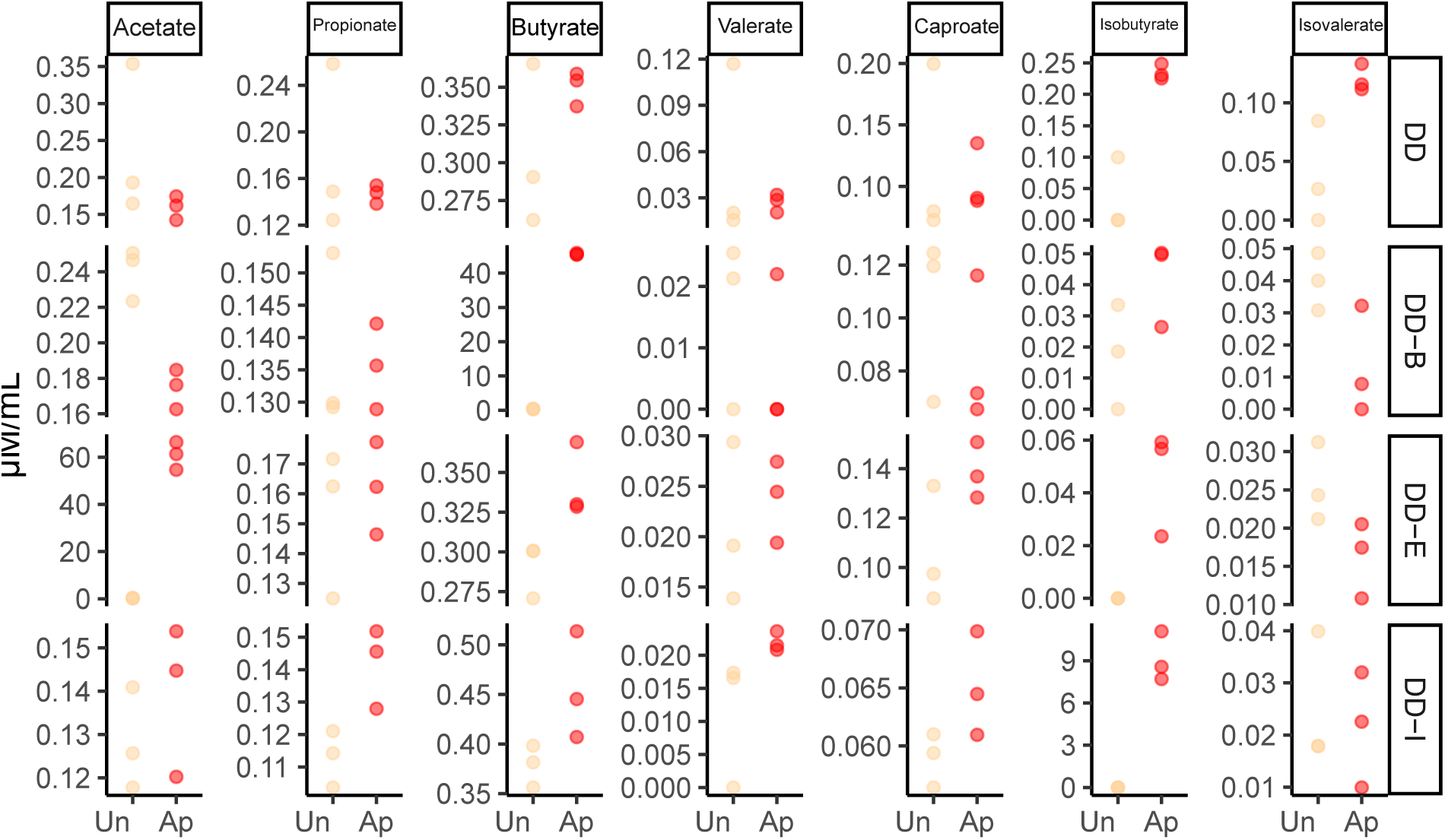
*A. pomorum* SCFA production can be manipulated by media alcohols. GC analysis of SCFAs produced by *A. pomorum* grown in *Drosophila* defined diet bacterial medium (DD), supplemented with nothing (DD) or with 1% butanol (DD-B), 1% ethanol (DD-E), or 1% isobutanol (DD-I). In each case, SCFA production by *A. pomorum* corresponds to what alcohol is present, with acetate increasing only when ethanol is present, butyrate when butanol is present, and isobutyrate when isobutanol is present.

**Figure S3.**
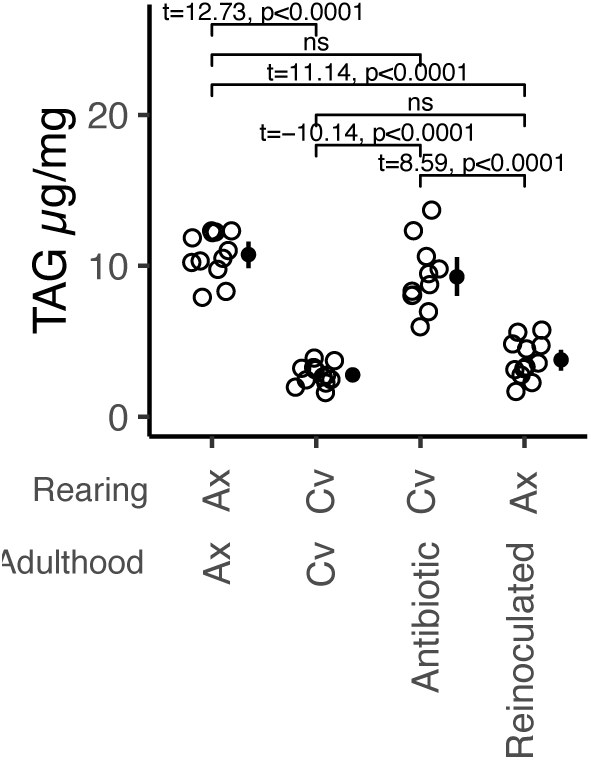
Conventionalized axenics and antibiotic treated conventional flies phenocopy TAG levels after 1 week. Axenic females fed male conditioned food for 1 week have reduced TAG levels equivalent to CR flies. CR flies fed a cocktail of antibiotics for 1 week exhibit TAG levels similar to axenics.

**Figure S4.**
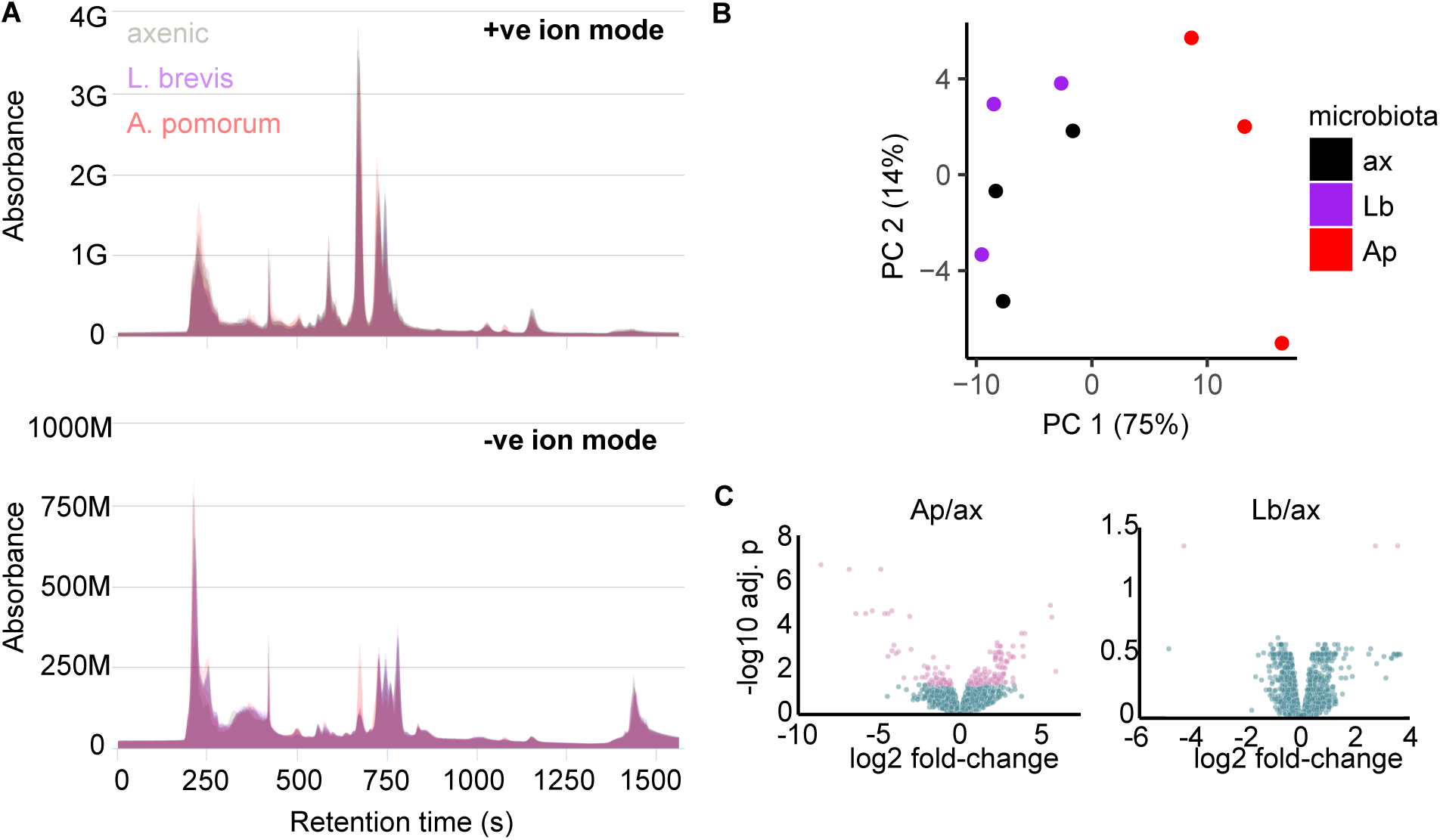
LC-MS small-molecule profiling as a tool to characterise impacts of *A. pomorum* and *L. brevis* on *Drosophila* metabolome. Females were reared to day 3 of adulthood with *A. pomorum, L. brevis*, or under axenic conditions. **A.** Chromatograms in positive and negative ion mode indicate consistent sample prep. **B.** PCA differentiates *Ap*-flies, but not *Lb-*flies, from axenics. **C.** Differential abundance analysis identifies specific peaks modulated in *Ap-*flies and *Lb-*flies. Volcano plots show peaks coloured by p-value threshold (FDR≤0.05), generated by IMP.

**Figure S5.**
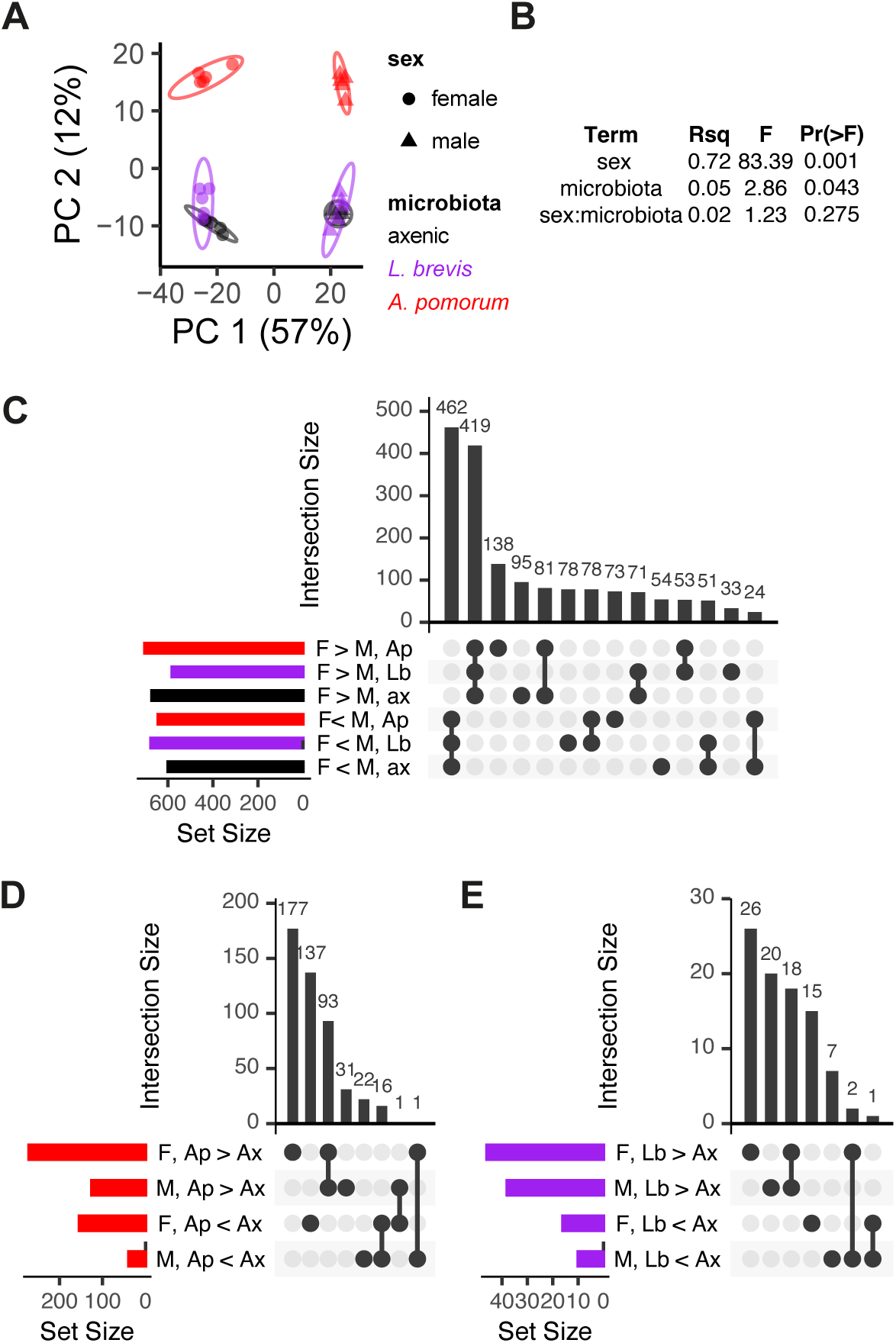
Negligible sexual dimorphism in metabolomic effect of microbiota. Metabolomes were characterised in female and male adult flies (3 days old), reared in gnotobiotic monoassociation with either *A. pomorum* or *L. brevis*. **A.** principal components analysis and **B.** PERMANOVA detected metabolic impacts of both sex and microbiota, but no evidence of interactions, suggesting that sex-specific effects are negligible. **C.** Sexual dimorphism in the metabolome is largely microbe-independent. UpSet plot showing intersections between sets of sexually dimorphic metabolites in the three microbiome conditions. F>M=peaks with significantly higher levels in females than males, *vice versa* F<M. Ap=*A. pomorum*, Lb=*L. brevis*, ax=axenic. Bars to left show total size of the six sets, bars above show sizes of intersections, specified by the connecting matrix. The two largest sets of sexually dimorphic peaks do not differ in response to microbiota, and no overall pattern emerges from the remaining smaller sets, suggesting that metabolomic sexual dimorphism is largely robust to microbiota. **D.** Male metabolomic responses to *A. pomorum* are largely also apparent in females, while responses in females show responses in additional peaks. UpSet analysis as per C, F=female, M=male. Ap>Ax=peaks that are significantly elevated in response to *A. pomorum* relative to axenics, *vice versa* Ap<Ax. **E.** Metabolomic response to *L. brevis* is diminutive, albeit with greater sex-specificity than response to *A. pomorum*. UpSet analysis and labelling as per D, Lb=*L. brevis*.

**Figure S6.**
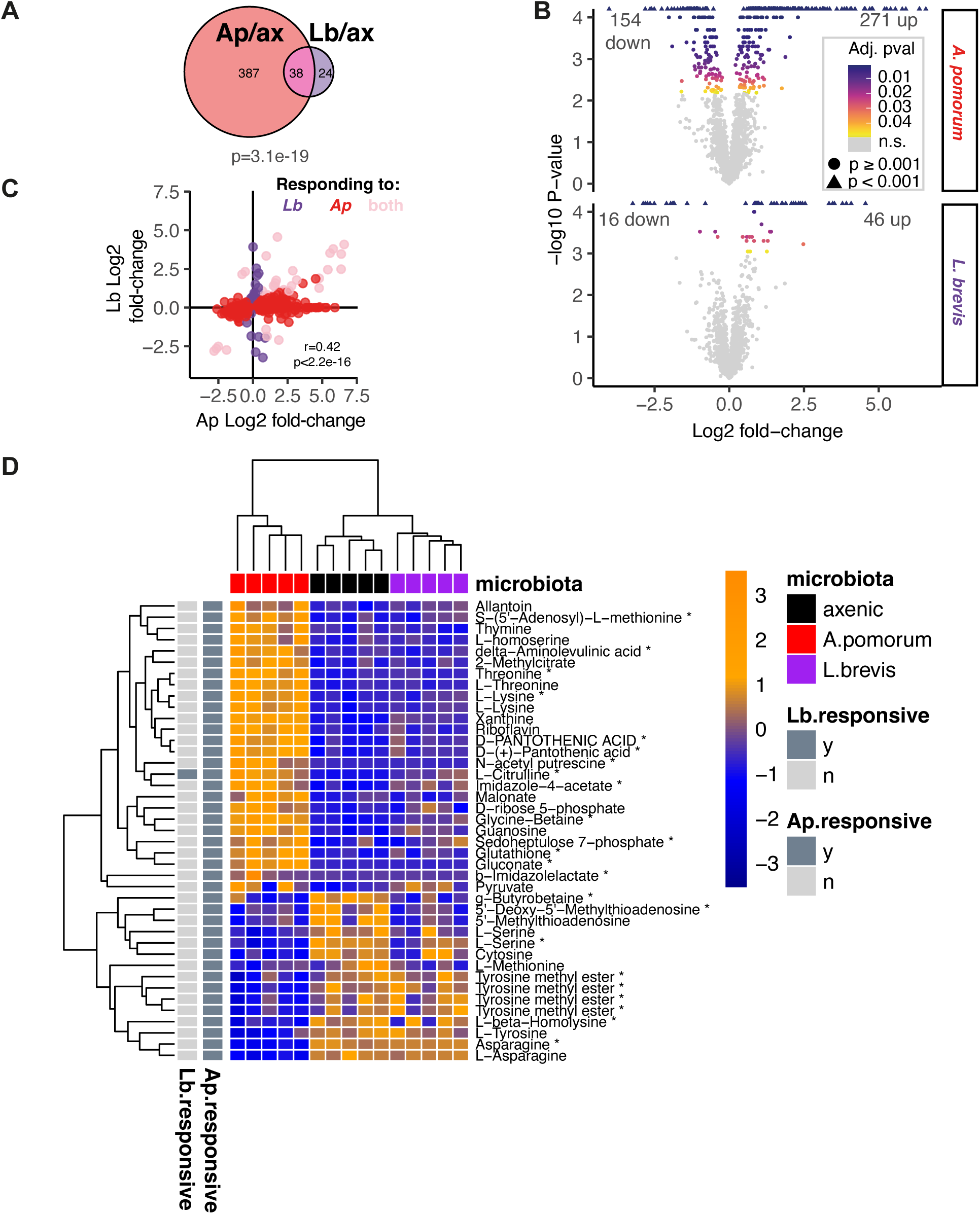
Metabolomic impacts of *A. pomorum* in females encapsulate those of *L. brevis*. **A.** *A. pomorum* has a more substantial metabolomic impact and accounts for the majority of impacts of *L. brevis*. P value from hypergeometric test shows overlap is significantly greater than expected by chance. **B.** Volcano plots of metabolomics data confirm *A. pomorum* influence substantial numbers of metabolites. Each facet shows impact of *A. pomorum* and *L. brevis* relative to axenics, respectively, for 3244 peaks detected by LC-MS. Points are coloured by adjusted P value (FDR), grey for non-significant effects (FDR≥0.05). Y axis gives untransformed P values on -log10 scale, values >4 (i.e. p<0.0001) are censored, represented by triangular points. Numbers of peaks significantly upregulated or downregulated in each condition are given at top corners of each plot. **C.** Peaks regulated by *A. pomorum* and/or *L. brevis* show significantly correlated response to either bacterium. Points are colour-coded by response to either or both bacteria. Spearman’s rho shows significant correlation. **D.** Metabolites identified definitively by comparison to definitive standards and/or fragmentation. Heatmap shows relative abundances (Z-scores) for metabolites that responded to either bacterium, given at right: all peaks shown responded to *A. pomorum*, one of which also responded to *L. brevis*, no other peaks responded to *L. brevis*. Microbiota conditions given at top.

**Figure S7.**
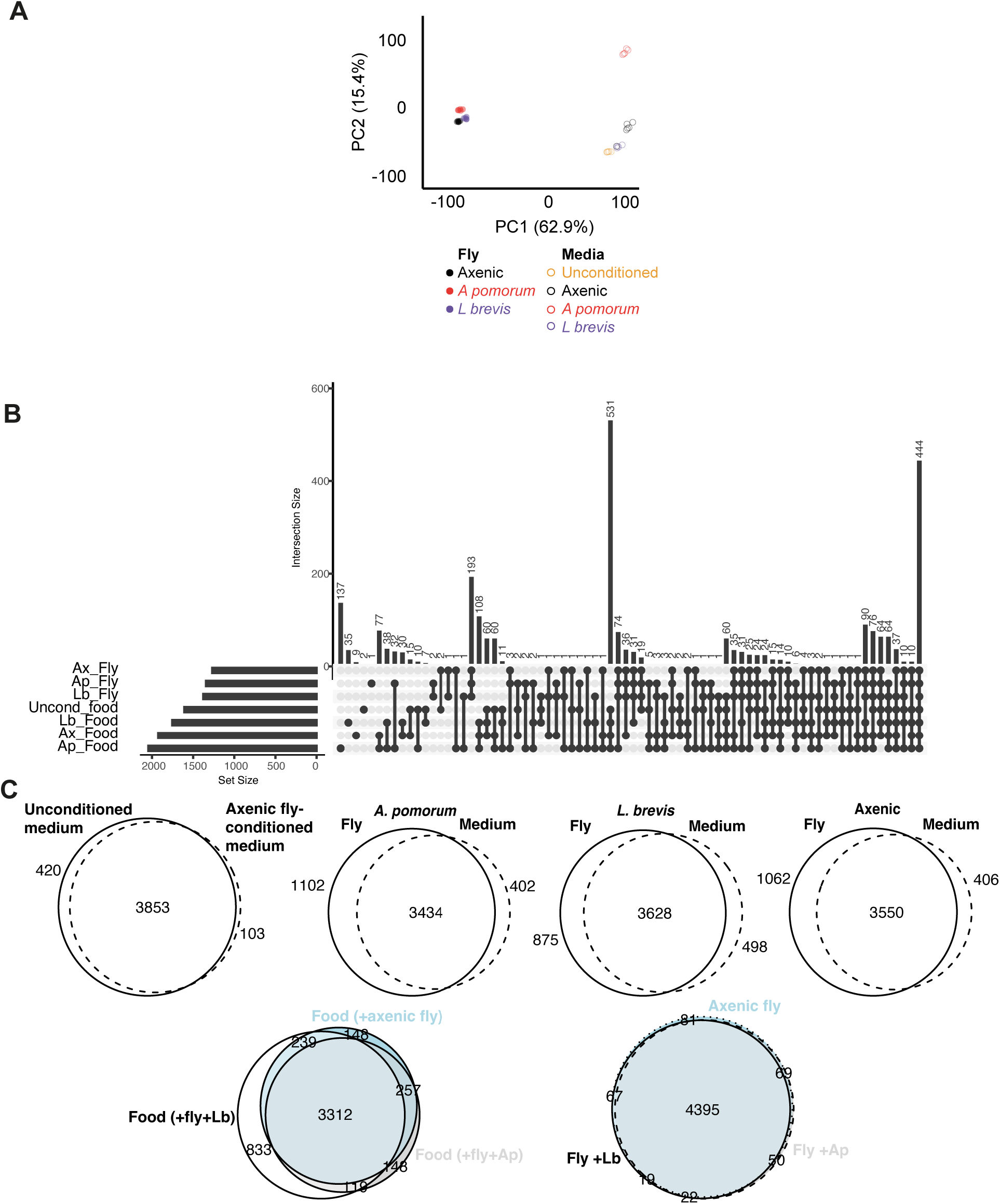
Characterising microbiota-dependent correspondence between chemical composition of flies and media. Experimental design shown in Figure 2A. **A.** The fly metabolome is less responsive to microbial variation than media. Principal components analysis showing the first two principal components shows larger effects of microbial variation on flies (filled points to left) than on media (clear points to right). Differentiation between unconditioned media and axenic media suggest that flies alter media composition independent of microbiota. **B.** Upset plot showing intersections among LC-MS peaks detected in media and fly samples. A peak is defined as detected in a given condition if detected in >20% of samples. Each row of the matrix represents the four different media conditions and three different fly conditions. Set sizes are given by barplot at left. Intersections between conditions are indicated by connected dots on the matrix, with intersection size given by barplot above. **C.** Euler diagrams showing specific intersections of particular interest. Peak presence/absence is defined as in A.

**Figure S8.**
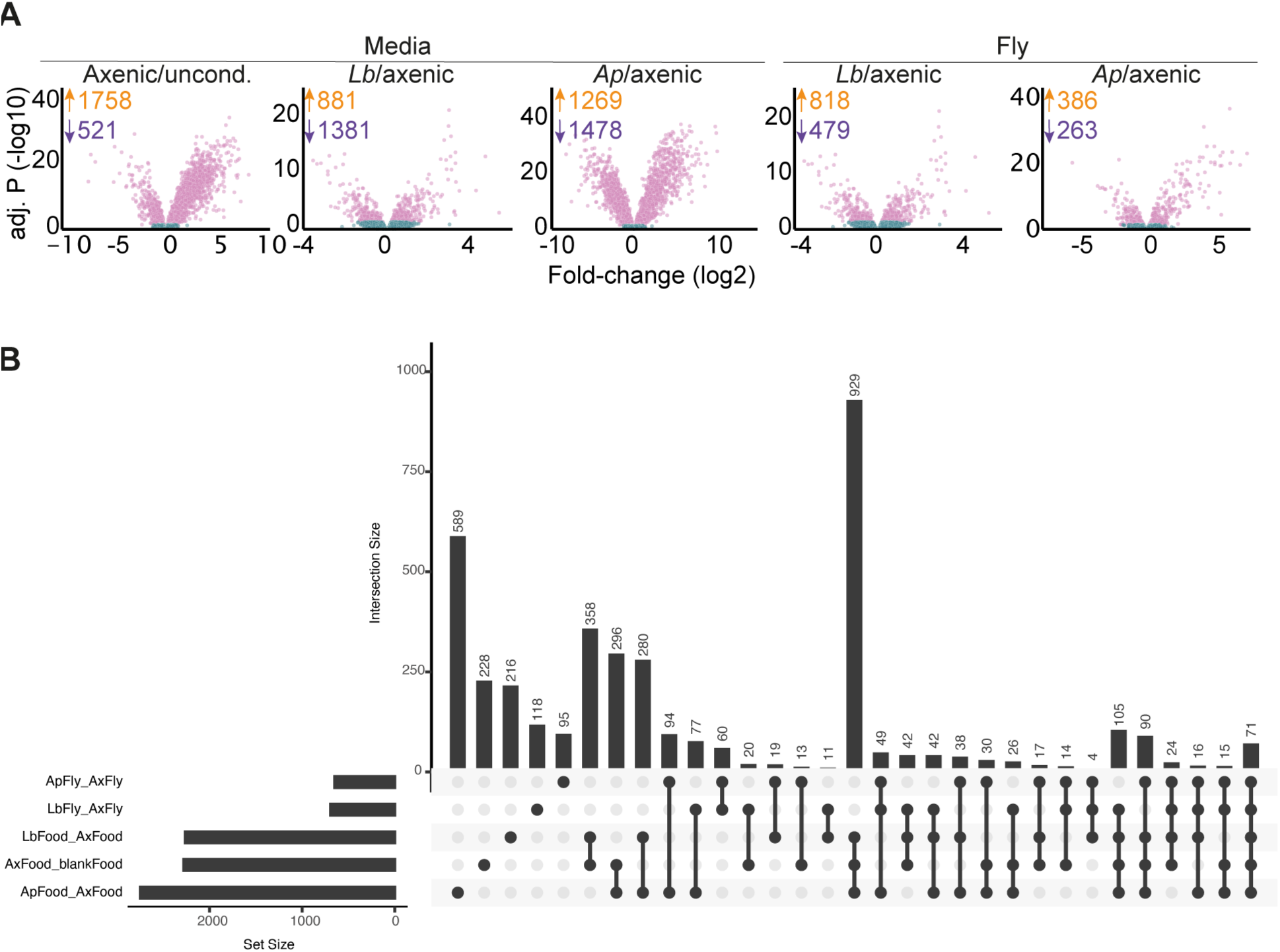
Differentially abundant LC-MS peaks in response to microbiota variation in flies and media. **A.** Volcano plots showing differentially abundant peaks (5880 total) in flies and media, in response to microbiota manipulation. Sample type (media or fly) and comparison of microbial conditions are given above each facet. Significantly different peaks (FDR ≤ 0.05) are coloured pink. Numbers of upregulated peaks are given in orange, numbers of downregulated peaks are given in purple. **B.** Upset plot showing intersections between sets of differentially-abundant peaks. Each row of the matrix represents the set of peaks found to be differentially abundant in each specified comparison. Set sizes are given by barplot at left. Intersections between conditions are indicated by connected dots on the matrix, with intersection size given by barplot above.

**Figure S9.**
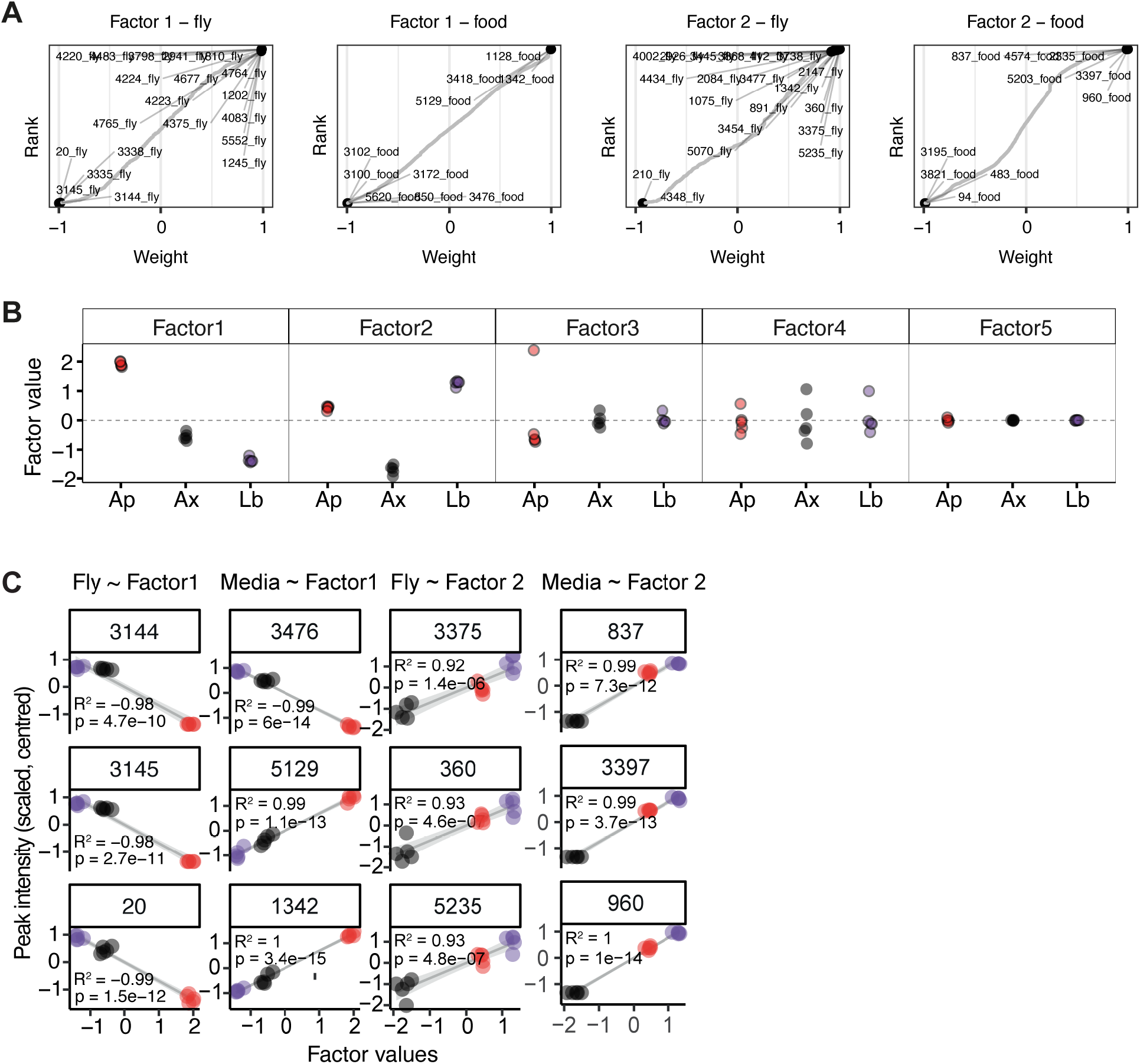
Multi-omic factor analysis (MOFA) QC plots. **A.** Factors 1 and 2, which explain most variance in fly and media (see Figure 2), are weighted by specific peaks. Facets show scaled weights for all peaks in the analysis, plotted against rank weight. Peak numbers for the top-weighted peaks are given with labels. The figures demonstrate that the factors represent abundances of peaks non-randomly. **B.** Factors 1 and 2 discriminate samples by microbiota status. Factor 1 appears to reflect impact of *A. pomorum* (Ap) on peak abundances, while Factor 2 appears to reflect difference between axenic versus bacteria-associated conditions. Factors 3-5 do not discriminate among experimental conditions, consistent with explaining little overall variance (see Figure 2). **C.** MOFA Factors 1 and 2 predict peak intensities well both for fly metabolomes and media nutrient composition. For each factor (1 or 2) and each sample view (media or fly) the peaks with top-three ranked weights (i.e. contribution to factor values) are shown. Latent variable values are plotted on X axes for factors 1 and 2, as specified to the top of each column. Sample view (media or fly) also specified to the top of each column. Peak ID given to the top of each individual plot. Grey lines represent linear regression, with variance in peak intensity explained by latent variable (R^2^) and statistically significant (P) for each peak-factor combination. The significant relationships between latent variables and peak intensities confirm that the latent variables represented by factors 1 and 2 are informative axes of variation in both the metabolome and nutrition.

**Figure S10.**
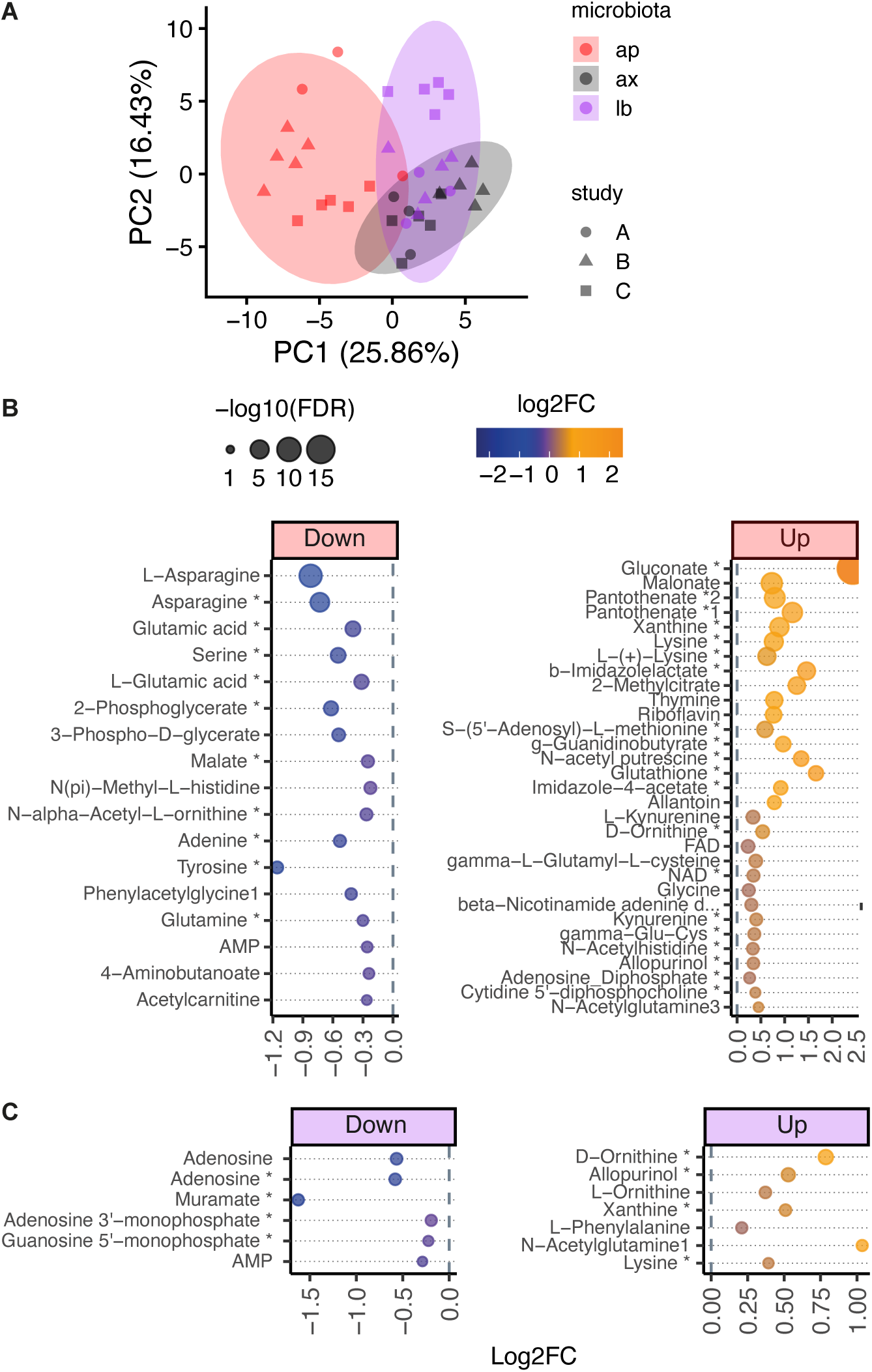
Supplementary materials for multi-study metabolomics. **A.** Unsupervised analysis finds repeatable microbiota-dependent variation in 130 metabolites across three biologically and technically independent LC-MS analyses. Ordination plot from multigroup PCA, coloured by microbiota condition, with point shapes showing replicate studies. Study A shown in Figure S4, study B Figures S5–S6, study C Figure 2. Ellipses represent 95% CIs around centroids. **B-C.** Differential abundance analysis (LIMMA) identifies metabolites up/down regulated relative to axenics in *Ap*-flies (B) and *Lb*-flies (C). Metabolites (y-axis) are ranked and point sizes are scaled by -log10 p-values; point colours reflect log2 fold-changes, also given on x-axes.

**Figure S11.**
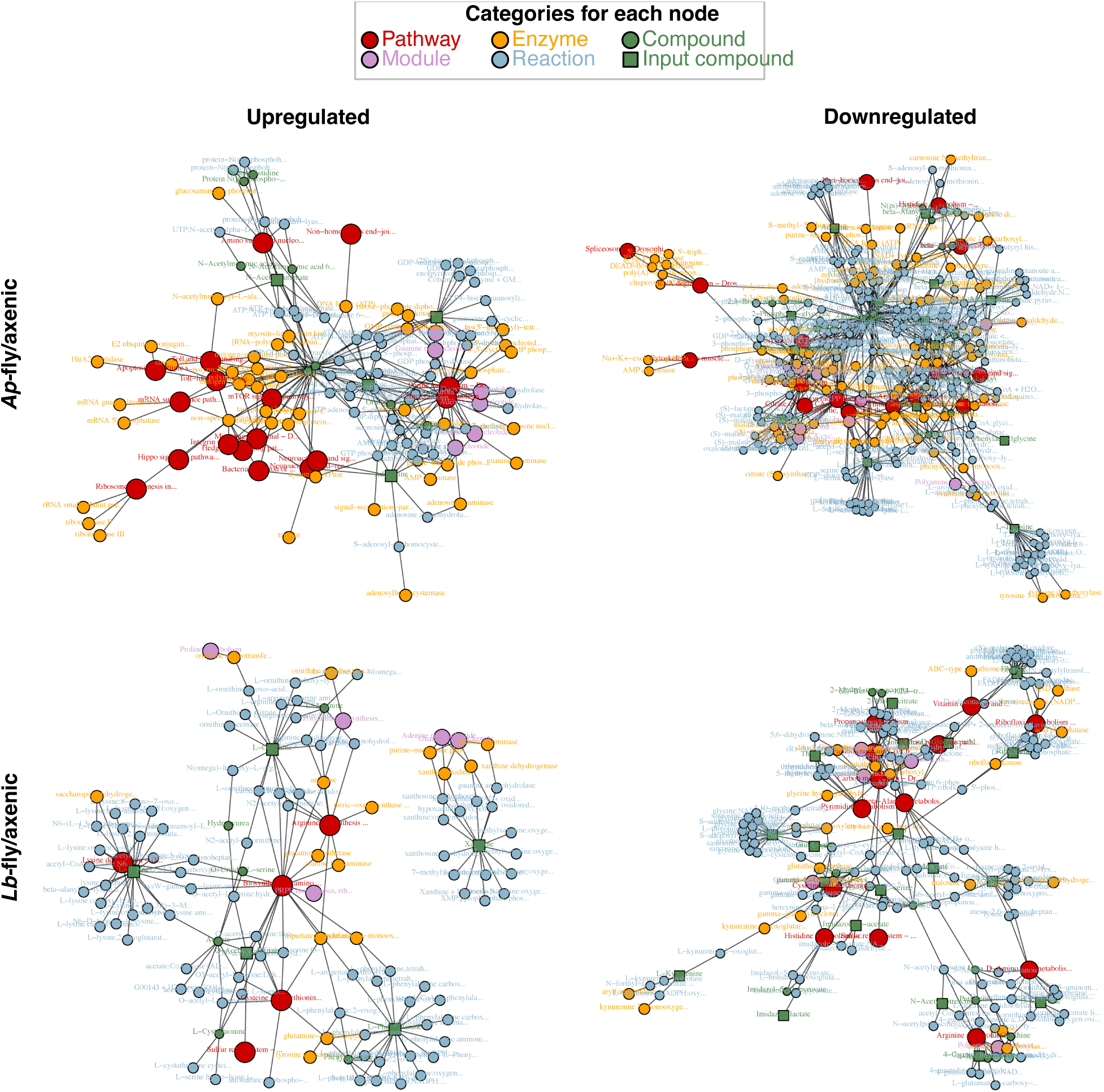
FELLA identifies coherent impacts of microbiota on KEGG enzymes, reactions, modules and pathways. Differentially-abundant metabolite sets (Figure S10B–C) were used as input. FELLA propagates a network from inputs, using the known graph structure of KEGG to incorporate enzymes, reactions, modules and pathways using guilt-by-association, and calculating enrichment. The method is appropriate given the sparsity of metabolomic data, i.e. the full set of metabolites in a given pathway will rarely all be detected. Enrichment scores are calculated to determine probability of findings at random. **A.** Graphs show networks propagated from input metabolites, upregulated/downregulated in response to *A. pomorum* and *L. brevis*. The coherence of the resulting networks indicates that input metabolites are found in closely-related metabolic networks.

**Figure S12.**
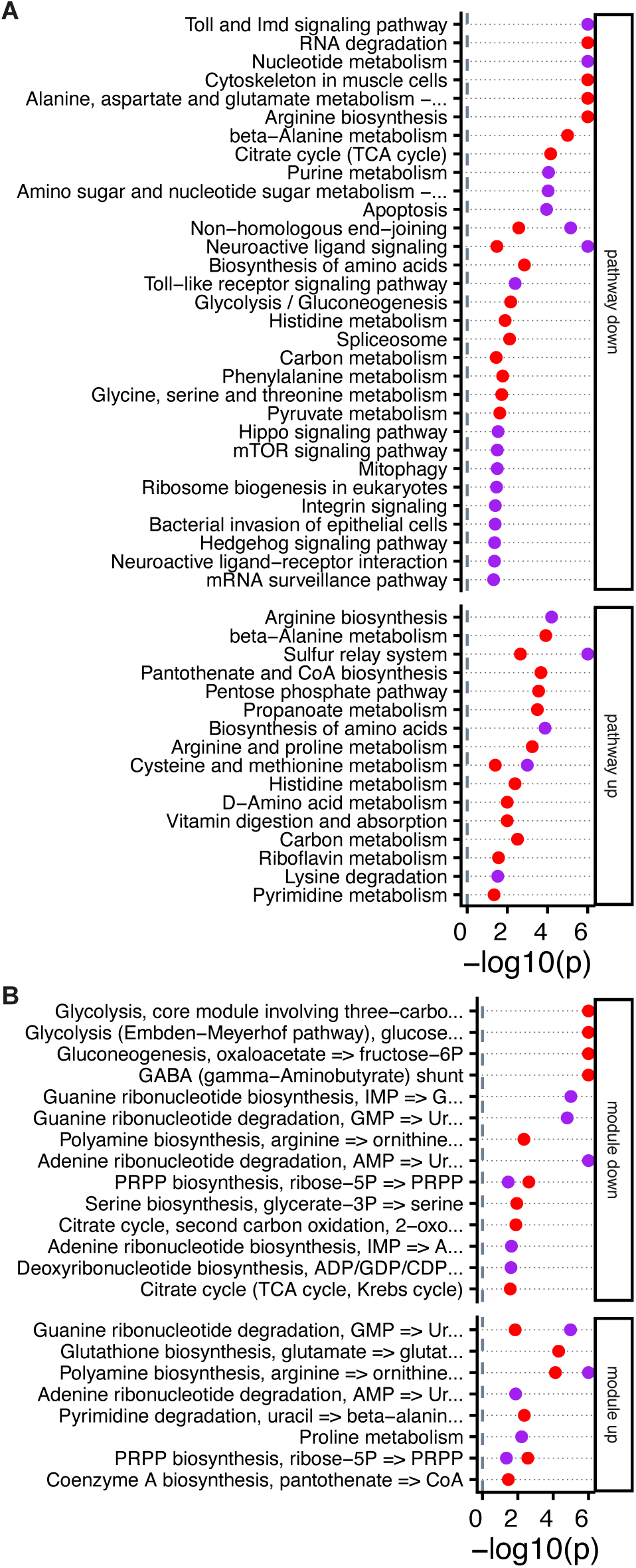
KEGG pathway and module enrichment among microbiota-responsive metabolites. **A-B.** KEGG pathway (A) and module (B) enrichment among differentially-abundant metabolites. Differential abundance testing performed by LIMMA (Figure S10B–C), enrichment analysis by FELLA (Figure S11). Panels distinguish enrichment among up/downregulated metabolites. Impacts of *A. pomorum*/*L. brevis* were analysed separately, but are presented on the same axes by red/purple dots (respectively).

**Figure S13.**
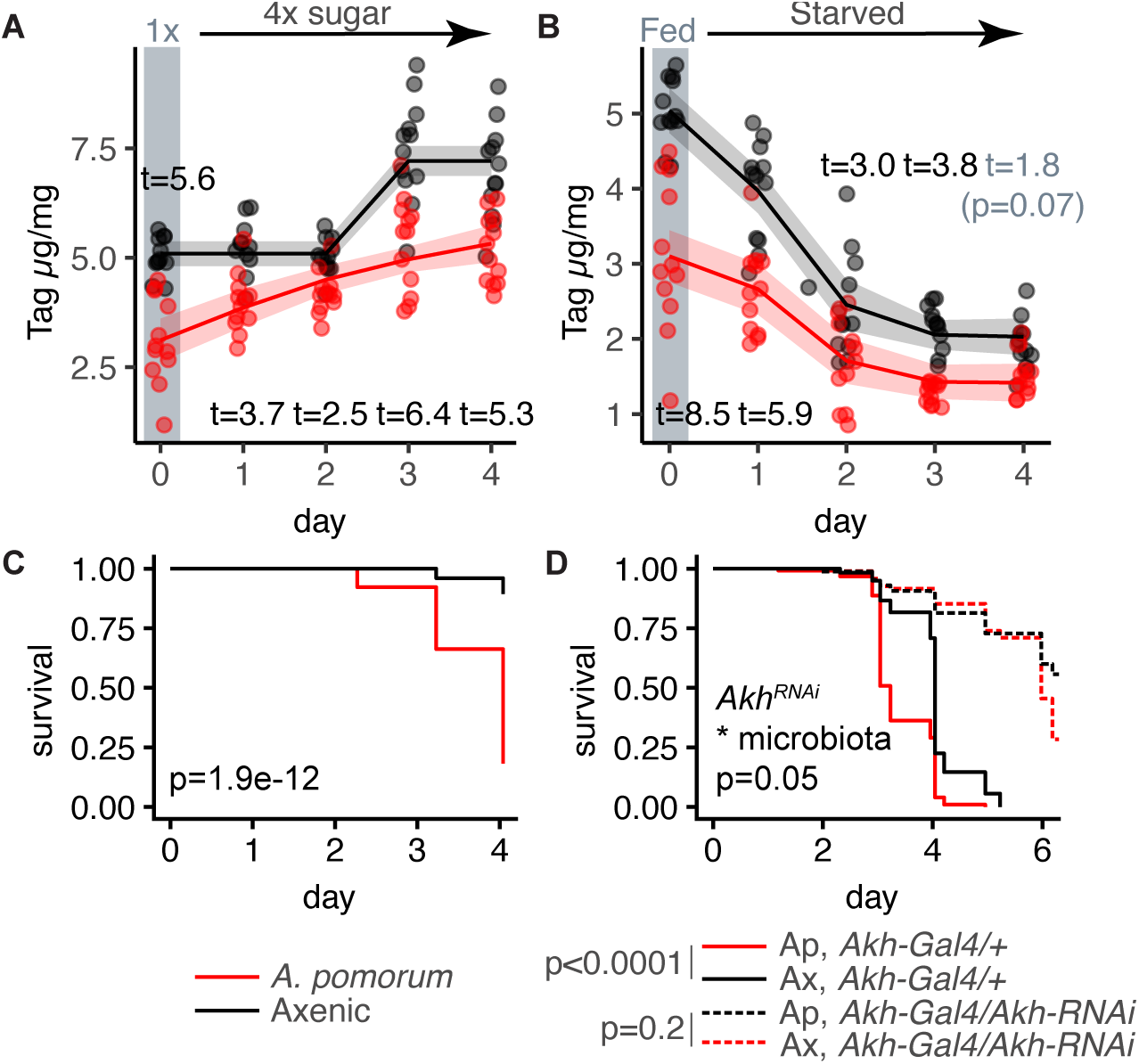
*A. pomorum* alters the dynamics of host lipid storage and mobilisation. **A.** *A. pomorum* alters kinetics of storing sugar as TAG. TAG was assayed in seven-day-old axenics/*Ap*-flies (1x/5% sucrose) before switching to 4x (20%) sucrose, followed by four days further TAG assayed. *Ap*-flies appear to accrue TAG monotonically while axenics initially fail to respond, followed by sharp increase after three days. Lines and ribbons indicate Weibull model fit with 95% confidence interval. t-statistics (on panel) from post-hoc tests (emmeans) of a 2-way ANOVA (time fitted as a factor), p<0.05 unless indicated (marginal effect in grey). **B-C.** Microbiota alter kinetics of lipid mobilisation upon starvation, with corresponding impacts on starvation resistance. Seven-day-old axenics/*Ap*-flies were switched to starvation medium (1.5% agarose). Statistics on B presented as per panel A (t-statistic at marginal significance highlighted in grey), showing that axenics mobilise lipid stores more rapidly at onset of starvation (<48h) before recalcitrance (>48h), after which axenic TAG plateaus above that of *Ap*-flies. Less complete *Ap*-fly lipid mobilisation (B) corresponds to reduced *Ap*-fly starvation resistance (C) – data presented as Kaplan-Meier plot, p-value from Cox proportional hazards model (axenics N=75, Ap-flies N=77). **D.** Starvation sensitivity in *Ap*-flies requires plasticity in lipid mobilisation. Capacity to upregulate lipid mobilisation was impaired genetically by knocking down adipokinetic hormone (AKH, fly orthologue of glucagon), constitutively rendering axenics and *Ap*-flies equivalently starvation-resistant. Statistics from Cox proportional hazards model, pairwise comparisons (key) from post-hoc emmeans tests.

**Figure S14.**
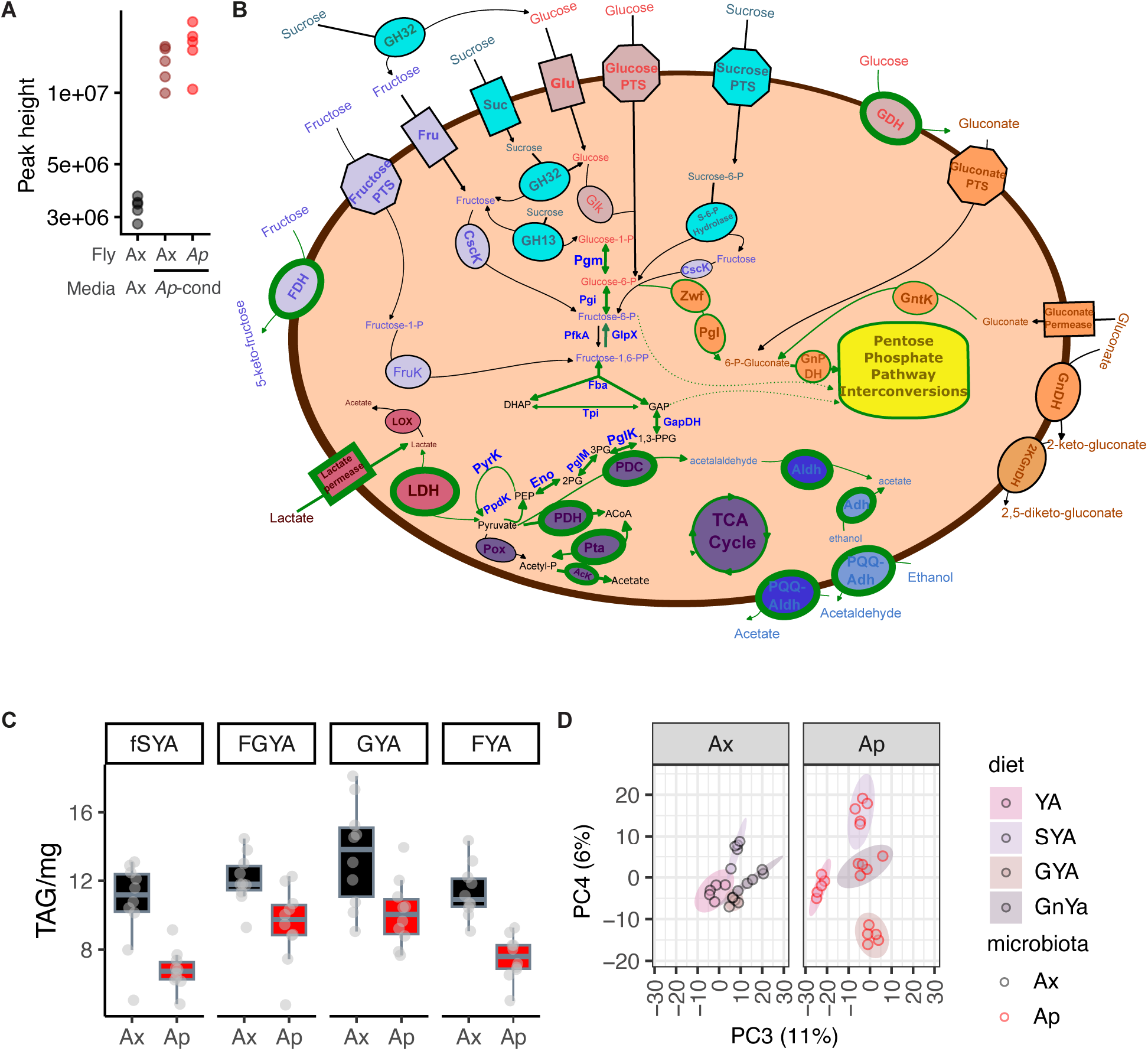
Roles of gluconate and sugars in fly/bacterial metabolism. **A.** *A. pomorum* produce gluconate under physiological condition in the fly. Data from metabolome analysis presented in Figure 2H. **B.** Bacterial metabolism of sucrose, fructose, glucose, lactate, and ethanol. The schematic depicts an amalgamation of strategies bacteria use to metabolize different carbon sources. Permeases and non-PEP dependent transporters are depicted as rectangles, PTS systems are depicted as octagons, and non-transporter enzymes are depicted as ovals. Proteins with a green outline are present in the *Acetobacter pomorum* DmCS004 genome, and reactions that A. pomorum can do are in green arrows. Proteins outlined in black, and black arrows are not predicted by the genome. Dotted lines mean the metabolites feed into the non-oxidative branch of the Pentose Phosphate Pathway (PPP). *A. pomorum* can gain energy from oxidizing ethanol to acetaldehyde, and then acetate, as well as from the oxidation of fructose, and the oxidation of glucose to gluconate. It does not further oxidize gluconate, and lacks transporters for all compounds except lactate (ethanol can enter the cell without a transporter). There are no glycoside hydrolases for sucrose degradation (GH32 family enzymes like invertase convert sucrose to glucose and fructose, and GH13 family contains sucrose phosphorylase). There are no kinases for the incorporation of glucose or fructose compounds into glycolysis, meaning that the EMP pathway functions in the direction of gluconeogenesis/the PPP, as lactate can be used for both carbon and energy via lactate dehydrogenase. The genome encodes gluconokinase (GntK) to incorporate free gluconate into the PPP, however, no gluconate transporter was identified in the genome. Protein abbreviations: Fru – general fructose transporter, Glu – general glucose transporter, Suc – general sucrose transporter. PTS phosphotransferase system. GDH – glucose dehydrogenase, GnDH – gluconate dehydrogenase, 2KGnDH – 2-keto-gluconate dehydrogenase, FDH – fructose dehydrogenase, LDH – lactate dehydrogenase, Adh – alcohol dehydrogenase, Aldh – aldehyde dehydrogenase, PDH – pyruvate dehydrogenase, GnPDH – 6-phosphogluconate dehydrogenase, PQQ-Adh – pyrroloquinoline quinone dependent alcohol dehydrogenase, PQQ-Aldh – pyrroloquinoline quinone dependent aldehyde dehydrogenase, GapDH - Glyceraldehyde 3-phosphate dehydrogenase. POX – pyruvate oxidase, LOX – lactate oxidase. AcK – acetate kinase, FruK – 1-phosphofructikinase, CscK – fructokinase, PfkA – phosphofructikinase, PglK – phosphoglycerate kinase, PyrK – pyruvate kinase, PpdK – pyruvate, phosphate dikinase, GntK - gluconokinase. Pta - phosphate acetyltransferase, Pgm – phosphoglucomutase, Pgi – phosphoglucose isomerase, GlpX – fructose 1,6 bisphosphatase, Fba – fructose bisphosphate aldolase, Tpi - triosephosphate isomerase, PglM – phosphoglycerate mutase, Eno – enolase. **C.** *A. pomorum* reduces fly TAG across a range of sugars which it is (variably) competent to use directly. Axenic and *Ap-*flies were fed diets containing different sugars. (fSYA=yeast + agar + 146mM sucrose i.e. the standard yeast-based medium used in previous experiments, filter-sterilised sucrose to prevent hydrolysis by autoclaving; FGYA yeast + agar + 146mM glucose and fructose (each), GYA=yeast + agar + 292mM glucose, FYA=yeast + agar + 292mM fructose), from days 3-10 of adulthood. No interaction of sugar source and response to *A. pomorum* detected (ANOVA p=0.74, post-hoc comparisons of *A. pomorum* effect p<0.005 for each diet). **D.** Microbiota reprogram host metabolomic response to dietary variation. Flies were fed diets with varied carbohydrate sources (YA=yeast + agar, SYA/GYA as panel C, GnYA=yeast + agar +292mM gluconate), from days 3-10 of adulthood. Panels show PCA of metabolomic analysis, components 3-4 (complementing PCs 1-2, Figure 4H): dietary variation in apparent in axenics, but substantially more pronounced in the presence of *A. pomorum*.

**Figure S15.**
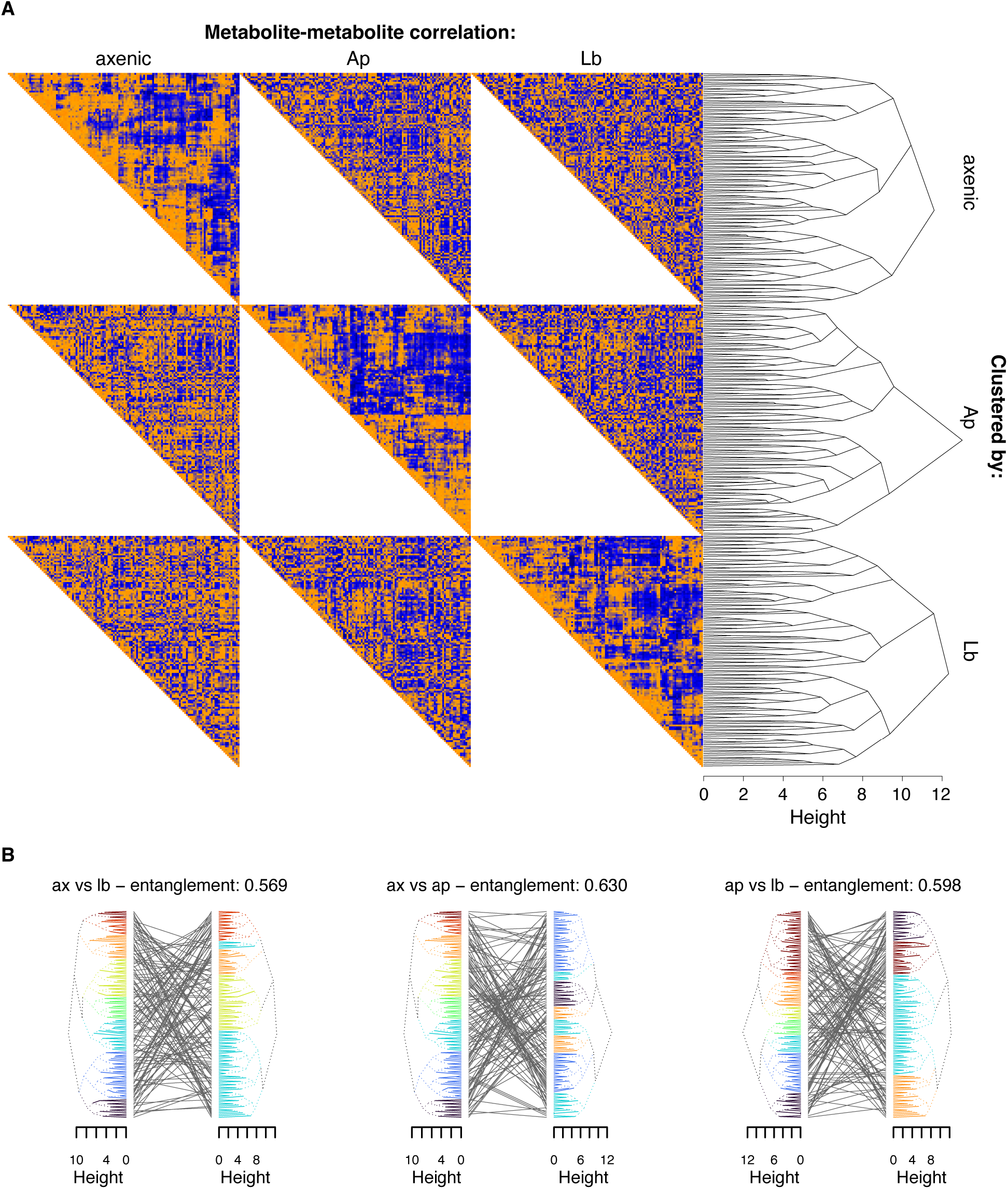
Microbiota alter metabolite co-regulation structure in the host metabolome. Among 130 metabolites (Figure 4), **A.** correlation matrices in each microbial condition (orange=high, blue=low). Matrices are organised by hierarchical clustering (to right). Each condition is organised by each clustering tree, showing that gross metabolic network structure is not static among microbiome conditions. **B.** Tanglegrams give pairwise comparisons of clustering in each microbial condition. For each facet, clustering trees (vis A) are compared by crossed lines in middle. Leaves are coloured by cluster membership, set by trees to the left – majority membership of clusters to the right are shown by equivalent colours – e.g. when comparing Ax vs *Lb*, the largest cluster in *Lb* (at bottom) contains more metabolites from the axenic teal cluster than any other. Entanglement scores indicate degree of differentiation between each pair of trees, (1=perfectly dissimilar, 0=identical).

**Figure S16.**
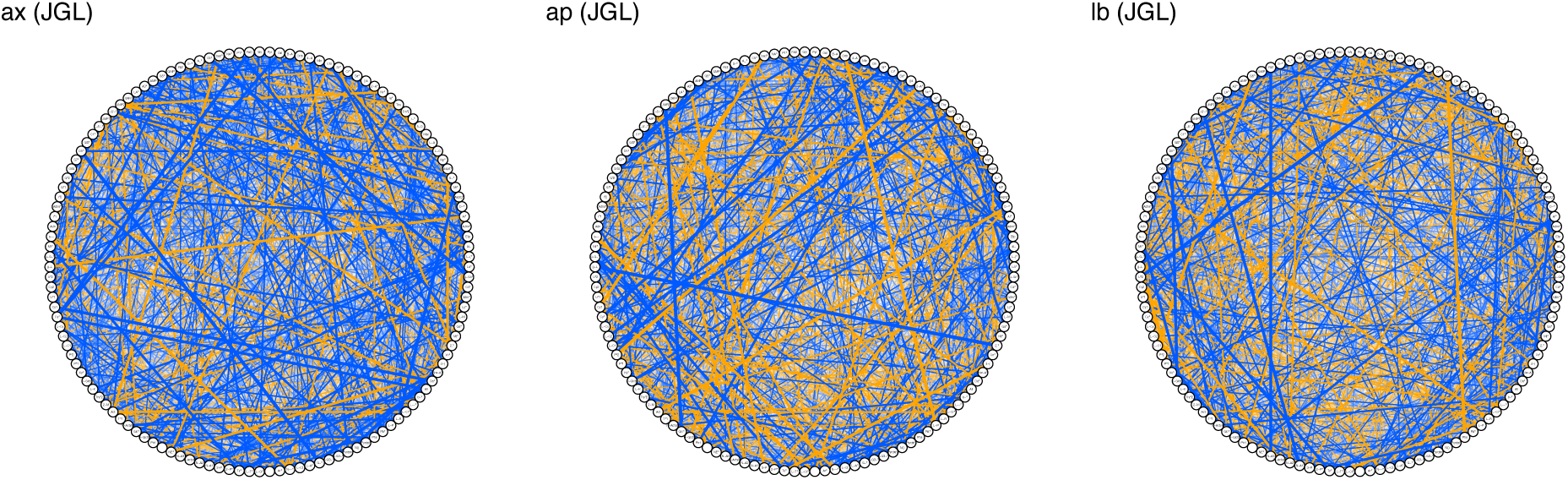
Microbiota restructure the host metabolic network. Metabolic networks were modelled using Joint Graphical Lasso to identify a consensus structure among the three conditions – given here by the ordering of 130 nodes (metabolites) around circles. Edge thickness indicates strength of correlation, colour indicates sign (orange=positive, blue=negative). All edges are shown.

**Figure S17.**
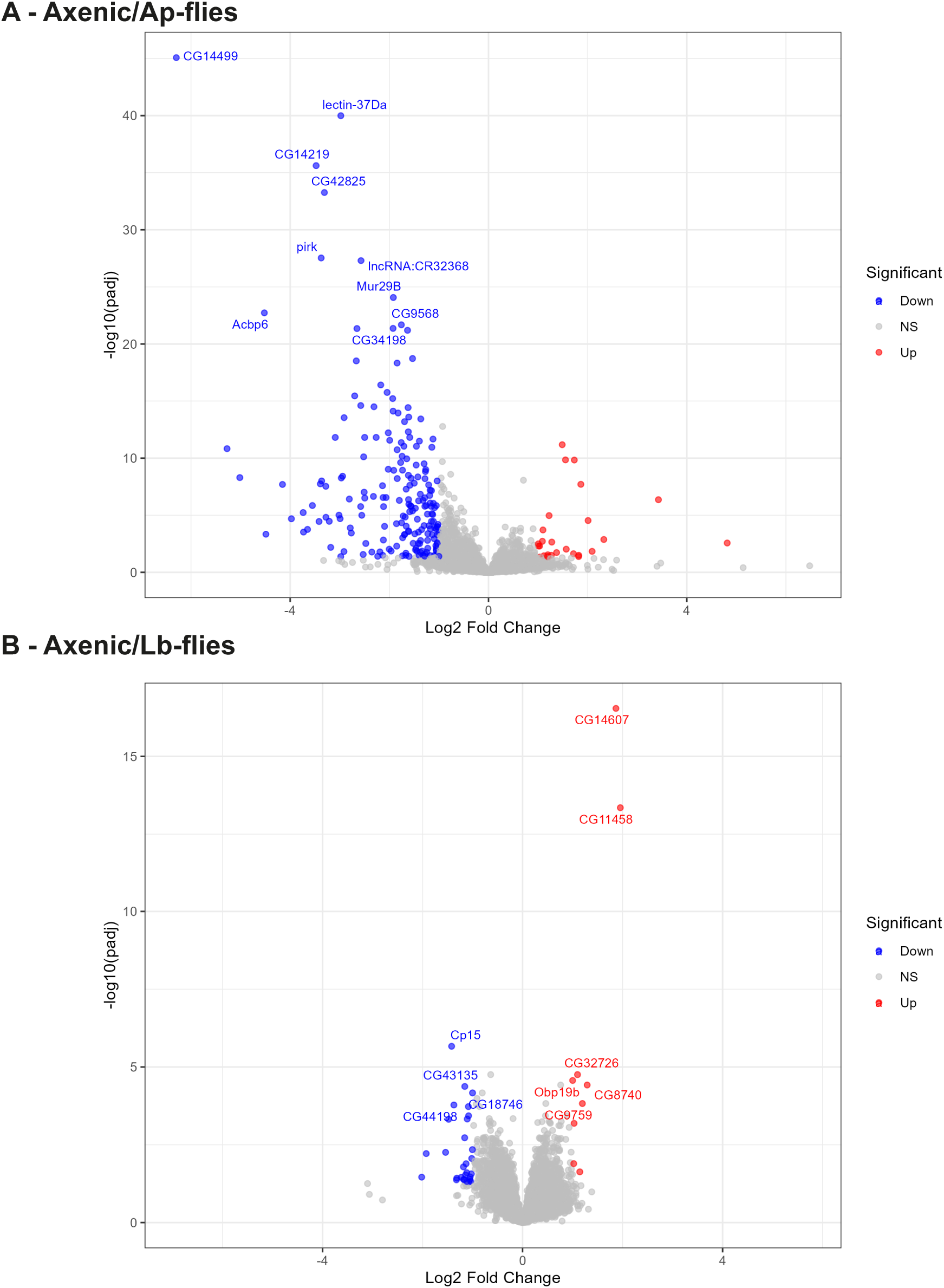
Differential gene expression in *Ap-*flies and *Lb-*flies, relative to axenics. Volcano plots show transcript-level differential abundance (each bacterial condition relative to axenics).

**Figure S18.**
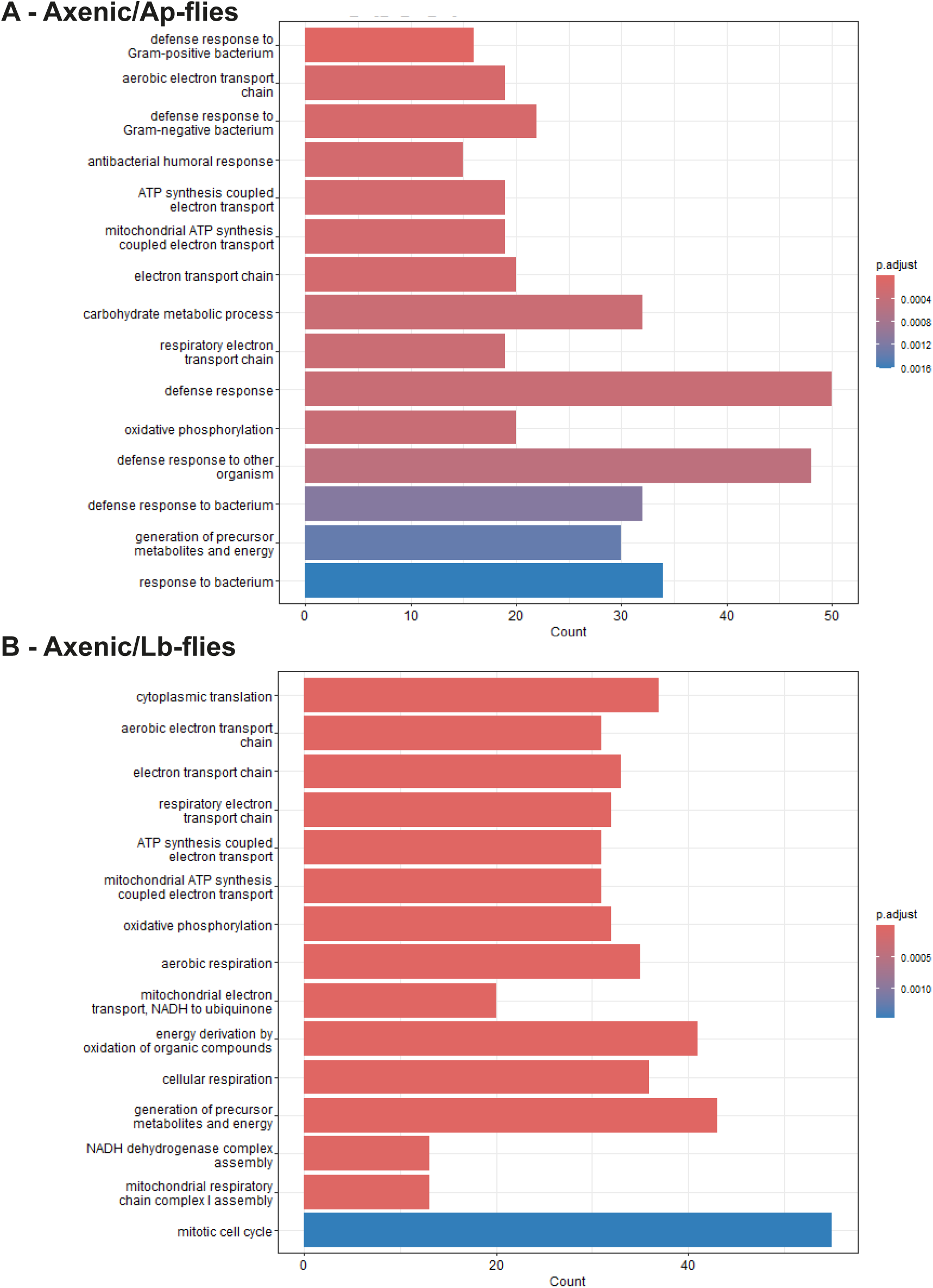
GO enrichment (biological process) in differential gene expression in *Ap-*flies and *Lb-*flies, relative to axenics.

**Figure S19.**
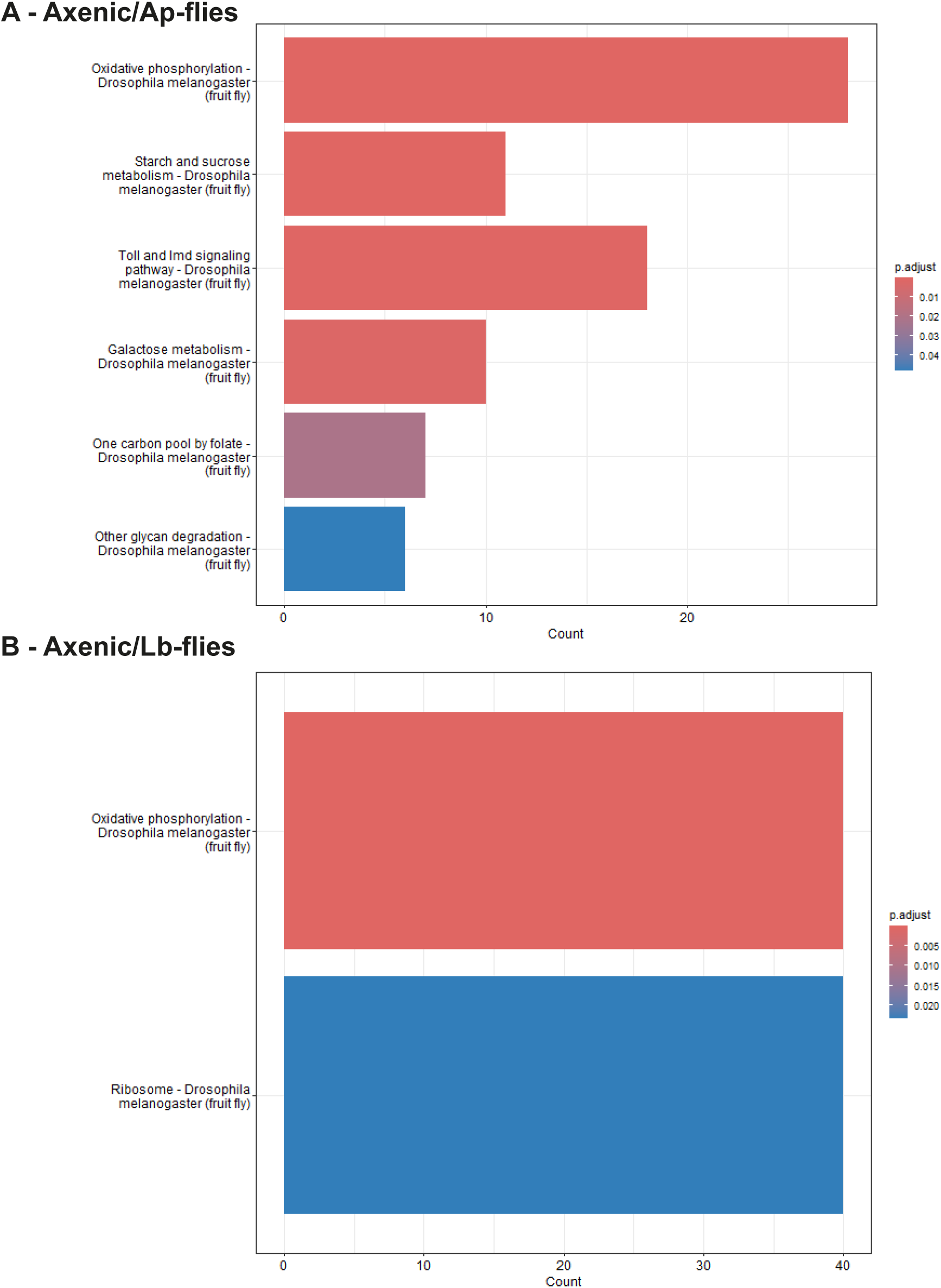
KEGG enrichment in differential gene expression in *Ap-*flies and *Lb-*flies, relative to axenics.

**Figure S20.**
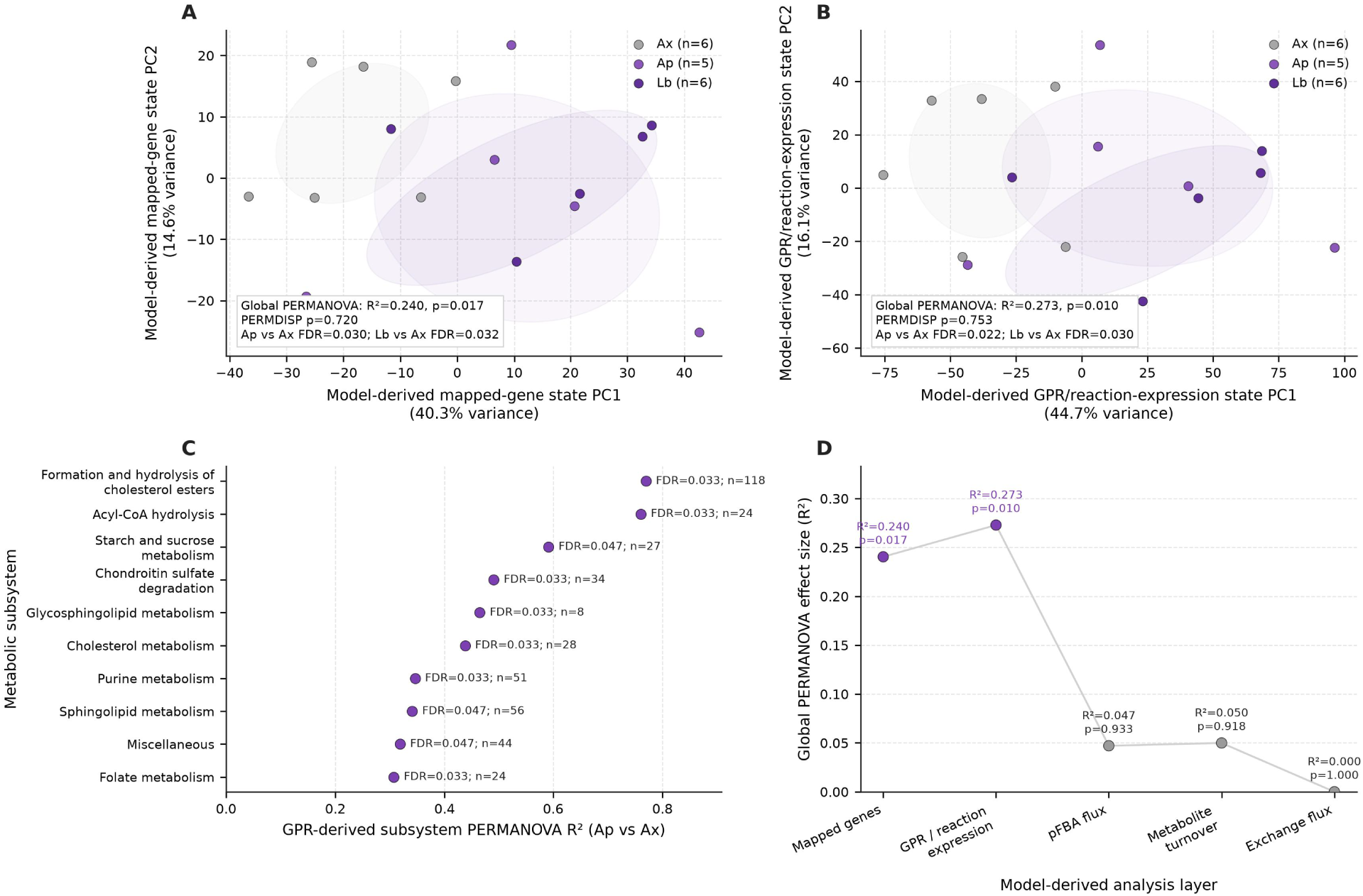
Differential flux-balance analysis shows wide-scale signature of GEM remodelling, without propagation to (predicted) transcriptional flux modulation. **A.** PCA of the model-derived mapped metabolic-gene state. **B.** PCA of the model-derived GPR/reaction-expression state. **C.** Ap-vs-Ax reaction-expression subsystems passing FDR <= 0.05; point position gives PERMANOVA R² and labels report FDR and variable-reaction count. **D.** global PERMANOVA effect size across successive model-derived analysis layers.

**Figure S21.**
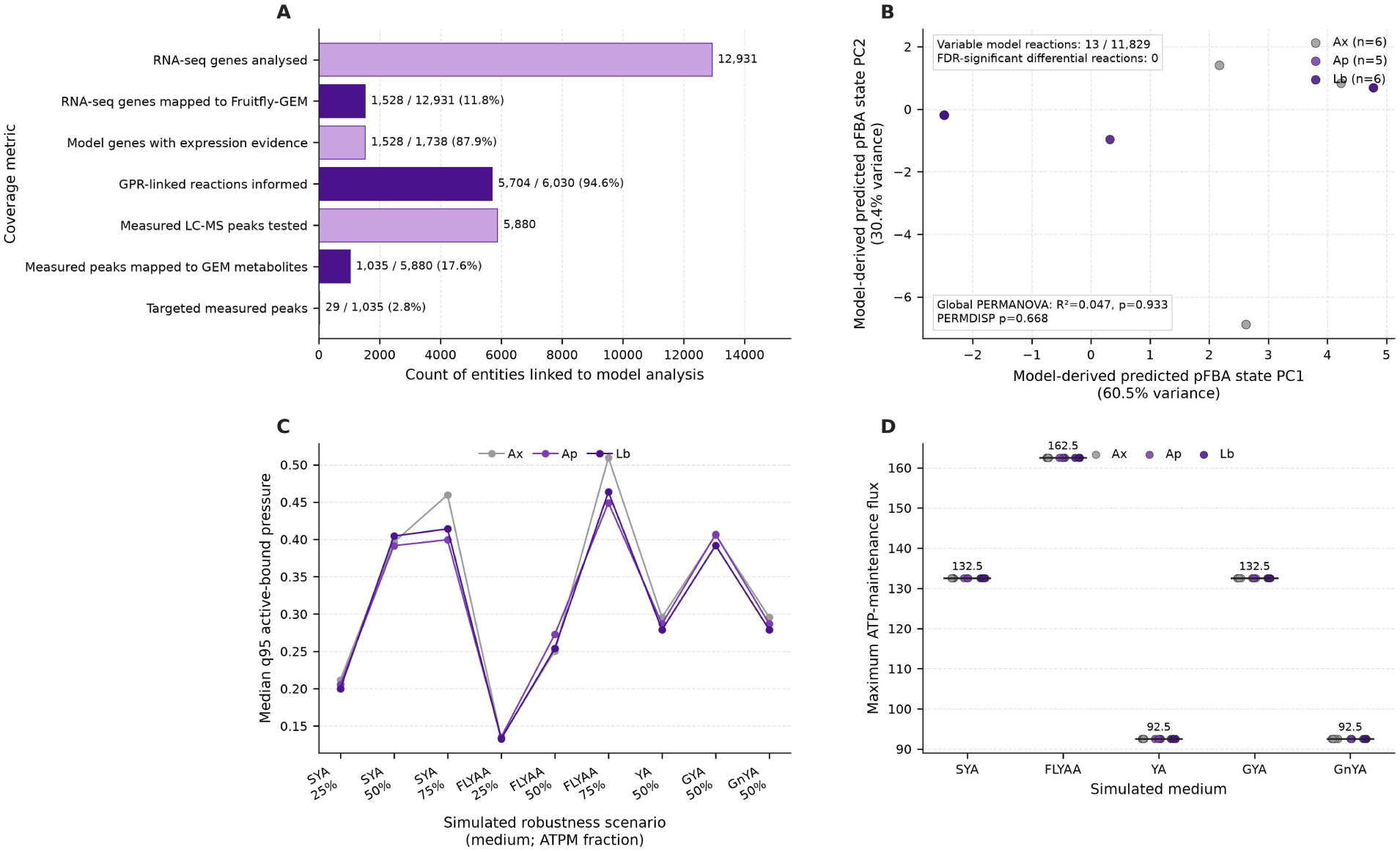
Differential flux-balance analysis: QC plots. **A.** coverage of RNA-seq, model-gene, GPR/reaction and metabolomics entities used in the modelling workflow. **B.** PCA of the predicted pFBA state; 13 of 11,829 reactions varied above the numerical tolerance and the global condition test was non-significant. **C.** robustness analysis of model-derived constraint engagement across simulated media and ATPM scenarios. **D.** simulated maximum ATP-maintenance capacity across media; values are identical across Ax, Ap and Lb within each medium.

**Figure S22.**
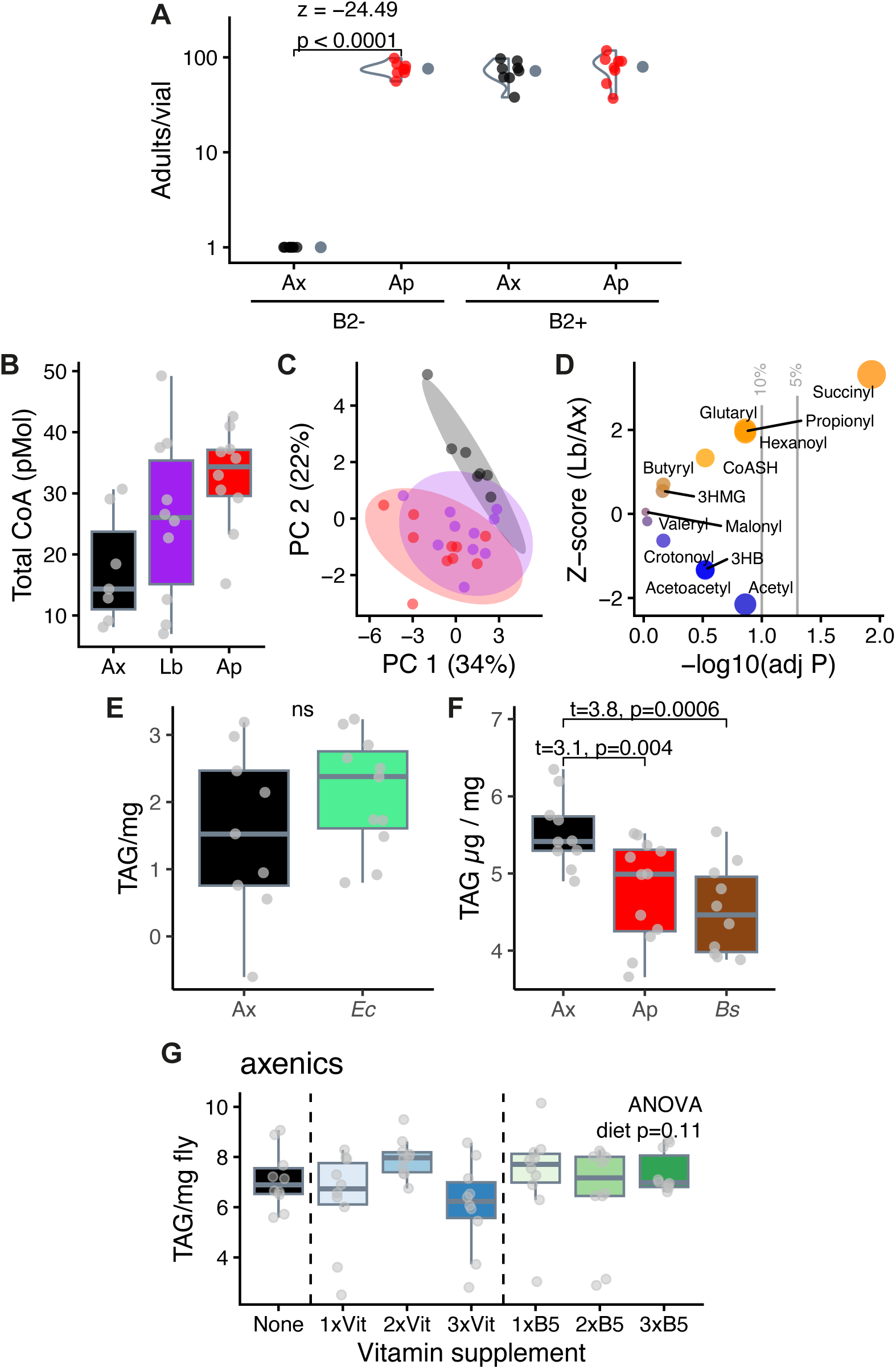
Effects of microbiota on fly vitamin biology. **A.** *A. pomorum* rescues fly development in riboflavin-free defined (holidic) diet. Adult counts taken 21 days post egg deposition on riboflavin-free defined diet (B_2_-) and complete diet (B_2_+). Zero-inflated GLM, microbiota*diet F=161.84, p<0.0001. Violin plots show density per condition, estimated marginal means given to right. **B-D.** *Levilactobacillus brevis* does not affect *Drosophila* coenzyme A metabolism substantially. Data from axenic (black) *Ap-*flies (red) are also presented in Figure 6. and **B.** Sum total CoA (CoASH + (acyl)CoA) is intermediate but not significantly increased by *L. brevis* (Ax vs. *Lb*, p=0.3, *Lb* vs. *Ap* p=0.35, *Ax* vs. *Ap* p=0.03). **C.** PCA indicates that overall multivariate differences in individual CoA species are also intermediate but not substantially different from axenics. **D.** The only significant impact of *L. brevis* is an increase in succinyl-CoA levels. **E-F** Impacts of *Escherichia coli* and *Bacillus subtilis* on host TAG. 3-day-old adult females reared on SYA medium, all flies with bacteria were monoassociated gnotobiotes. Data expressed as µg TAG per mg fly. **E.** *E. coli (Ec)* does not reduce TAG, ANOVA p=0.2. **F.** *B. subtilis (Bs)* recapitulates impact of *A. pomorum (Ap)*. ANOVA F_2,29_=8.24, p=0.001. Statistics on figure from post-hoc (emmeans) tests. **G.** Pantothenate and/or B-vitamin supplementation alone is not sufficient to rescue fly TAG. 3-day-old axenics were fed a B-vitamin supplement titration (after ^36^), or pantothenate alone for 1 week. Data expressed as µg TAG per mg fly.

## Declaration of generative AI and AI-assisted technologies in the manuscript preparation process

During the preparation of this work, the author(s) used Claude for manuscript proofing. The author(s) reviewed and edited the output as needed and take full responsibility for the content of the published article.

## Bibliography

1. McFall-Ngai, M., Hadfield, M.G., Bosch, T.C., Carey, H.V., Domazet-Lošo, T., Douglas, A.E., Dubilier, N., Eberl, G., Fukami, T., and Gilbert, S.F. (2013). Animals in a bacterial world, a new imperative for the life sciences. Proc Natl Acad Sci U S A 110, 3229–3236.

2. Shi, Z., Zou, J., Zhang, Z., Zhao, X., Noriega, J., Zhang, B., Zhao, C., Ingle, H., Bittinger, K., and Mattei, L.M. (2019). Segmented filamentous bacteria prevent and cure rotavirus infection. Cell 179, 644–658. e613.

3. Steele, M.I., Motta, E.V., Gattu, T., Martinez, D., and Moran, N.A. (2021). The gut microbiota protects bees from invasion by a bacterial pathogen. Microbiology spectrum 9, e00394–00321.

4. Yang, J.L., Zhu, H., Sadh, P., Aumiller, K., Guvener, Z.T., and Ludington, W.B. (2025). Commensal acidification of specific gut regions produces a protective priority effect against enteropathogenic bacterial infection. Applied and Environmental Microbiology 91, e00707–00725.

5. Blum, J.E., Fischer, C.N., Miles, J., and Handelsman, J. (2013). Frequent replenishment sustains the beneficial microbiome of Drosophila melanogaster. MBio 4, e00860–00813.

6. Collins, S.L., and Patterson, A.D. (2020). The gut microbiome: an orchestrator of xenobiotic metabolism. Acta Pharmaceutica Sinica B 10, 19–32.

7. Rescigno, M. (2014). Intestinal microbiota and its effects on the immune system. Cellular Microbiology 16, 1004–1013.

8. Vuong, H.E., Yano, J.M., Fung, T.C., and Hsiao, E.Y. (2017). The microbiome and host behavior. Annual review of neuroscience 40, 21–49.

9. Schretter, C.E., Vielmetter, J., Bartos, I., Marka, Z., Marka, S., Argade, S., and Mazmanian, S.K. (2018). A gut microbial factor modulates locomotor behaviour in Drosophila. Nature 563, 402–406.

10. Cabirol, A., Quinn, A., Schafer, J., Neuschwander, N., Kesner, L., Liberti, J., and Engel, P. (2026). A defined community of core gut microbiota members promotes cognitive performance in honey bees. Proceedings of the National Academy of Sciences 123, e2608600123.

11. Kolobaric, A., Andreescu, C., Jašarević, E., Hong, C., Roh, H., Cheong, J., Kim, Y., Shin, T., Kang, C., and Kwon, C. (2024). Gut microbiome predicts cognitive function and depressive symptoms in late life. Molecular Psychiatry 29, 3064–3075.

12. Cabreiro, F., and Gems, D. (2013). Worms need microbes too: microbiota, health and aging in Caenorhabditis elegans. EMBO molecular medicine 5, 1300–1310.

13. Guo, L., Karpac, J., Tran, S.L., and Jasper, H. (2014). PGRP-SC2 promotes gut immune homeostasis to limit commensal dysbiosis and extend lifespan. Cell 156, 109–122.

14. Thevaranjan, N., Puchta, A., Schulz, C., Naidoo, A., Szamosi, J.C., Verschoor, C.P., Loukov, D., Schenck, L.P., Jury, J., and Foley, K.P. (2017). Age-associated microbial dysbiosis promotes intestinal permeability, systemic inflammation, and macrophage dysfunction. Cell host & microbe 21, 455–466. e454.

15. Sharon, G., Segal, D., Ringo, J.M., Hefetz, A., Zilber-Rosenberg, I., and Rosenberg, E. (2010). Commensal bacteria play a role in mating preference of *Drosophila melanogaster*. Proc Natl Acad Sci U S A 107, 20051–20056.

16. Ashonibare, V.J., Akorede, B.A., Ashonibare, P.J., Akhigbe, T.M., and Akhigbe, R.E. (2024). Gut microbiota-gonadal axis: the impact of gut microbiota on reproductive functions. Frontiers in immunology 15, 1346035.

17. Shin, S.C., Kim, S.-H., You, H., Kim, B., Kim, A.C., Lee, K.-A., Yoon, J.-H., Ryu, J.-H., and Lee, W.-J. (2011). *Drosophila* microbiome modulates host developmental and metabolic homeostasis via insulin signaling. Science 334, 670–674.

18. Dodd, D., Spitzer, M.H., Van Treuren, W., Merrill, B.D., Hryckowian, A.J., Higginbottom, S.K., Le, A., Cowan, T.M., Nolan, G.P., and Fischbach, M.A. (2017). A gut bacterial pathway metabolizes aromatic amino acids into nine circulating metabolites. Nature 551, 648–652.

19. Cox, L.M., Yamanishi, S., Sohn, J., Alekseyenko, A.V., Leung, J.M., Cho, I., Kim, S.G., Li, H., Gao, Z., and Mahana, D. (2014). Altering the intestinal microbiota during a critical developmental window has lasting metabolic consequences. Cell 158, 705–721.

20. Turnbaugh, P.J., Ley, R.E., Mahowald, M.A., Magrini, V., Mardis, E.R., and Gordon, J.I. (2006). An obesity-associated gut microbiome with increased capacity for energy harvest. nature 444, 1027–1031.

21. Kawano, Y., Edwards, M., Huang, Y., Bilate, A.M., Araujo, L.P., Tanoue, T., Atarashi, K., Ladinsky, M.S., Reiner, S.L., and Wang, H.H. (2022). Microbiota imbalance induced by dietary sugar disrupts immune-mediated protection from metabolic syndrome. Cell 185, 3501–3519. e3520.

22. Sannino, D.R., Dobson, A.J., Edwards, K., Angert, E.R., and Buchon, N. (2018). The Drosophila melanogaster Gut Microbiota Provisions Thiamine to Its Host. mBio 9, e00155–00118.

23. Harrison, R.E., Yang, X., Eum, J.H., Martinson, V.G., Dou, X., Valzania, L., Wang, Y., Boyd, B.M., Brown, M.R., and Strand, M.R. (2023). The mosquito Aedes aegypti requires a gut microbiota for normal fecundity, longevity and vector competence. Communications Biology 6, 1154.

24. Torrallardona, D., Harris, C.I., and Fuller, M.F. (2003). Lysine synthesized by the gastrointestinal microflora of pigs is absorbed, mostly in the small intestine. American Journal of Physiology-Endocrinology and Metabolism 284, E1177–E1180.

25. Guetterman, H.M., Huey, S.L., Knight, R., Fox, A.M., Mehta, S., and Finkelstein, J.L. (2022). Vitamin B-12 and the gastrointestinal microbiome: a systematic review. Advances in Nutrition 13, 530–558.

26. Consuegra, J., Grenier, T., Baa-Puyoulet, P., Rahioui, I., Akherraz, H., Gervais, H., Parisot, N., Da Silva, P., Charles, H., and Calevro, F. (2020). Drosophila-associated bacteria differentially shape the nutritional requirements of their host during juvenile growth. PLoS biology 18, e3000681.

27. Huang, J.-H., and Douglas, A.E. (2015). Consumption of dietary sugar by gut bacteria determines Drosophila lipid content. Biology letters 11, 20150469.

28. Chaston, J.M., Newell, P.D., and Douglas, A.E. (2014). Metagenome-wide association of microbial determinants of host phenotype in *Drosophila melanogaster*. MBio 5, e01631–01614.

29. Dobson, A.J., Chaston, J.M., and Douglas, A.E. (2016). The Drosophila transcriptional network is structured by microbiota. BMC genomics 17, 975.

30. Quinn, A., El Chazli, Y., Escrig, S., Daraspe, J., Neuschwander, N., McNally, A., Genoud, C., Meibom, A., and Engel, P. (2024). Host-derived organic acids enable gut colonization of the honey bee symbiont Snodgrassella alvi. Nature microbiology 9, 477–489.

31. Regan, M.D., Chiang, E., Liu, Y., Tonelli, M., Verdoorn, K.M., Gugel, S.R., Suen, G., Carey, H.V., and Assadi-Porter, F.M. (2022). Nitrogen recycling via gut symbionts increases in ground squirrels over the hibernation season. Science 375, 460–463.

32. Chandler, J.A., Lang, J.M., Bhatnagar, S., Eisen, J.A., and Kopp, A. (2011). Bacterial communities of diverse Drosophila species: ecological context of a host–microbe model system. PLoS genetics 7, e1002272.

33. Newell, P.D., and Douglas, A.E. (2014). Interspecies interactions determine the impact of the gut microbiota on nutrient allocation in *Drosophila melanogaster*. Appl Environ Microbiol 80, 788–796.

34. Piper, M.D., Blanc, E., Leitão-Gonçalves, R., Yang, M., He, X., Linford, N.J., Hoddinott, M.P., Hopfen, C., Soultoukis, G.A., and Niemeyer, C. (2014). A holidic medium for *Drosophila melanogaster*. Nat Methods 11, 100–105.

35. Leitão-Gonçalves R, C.-S.Z., Francisco AP, Fioreze GT, Anjos M, Baltazar C, et al. (2017). Commensal bacteria and essential amino acids control food choice behavior and reproduction. PLoS Biol 15, e2000862.

36. Wong, A.C.-N., Dobson, A.J., and Douglas, A.E. (2014). Gut microbiota dictates the metabolic response of *Drosophila* to diet. J Exp Biol 217, 1894–1901.

37. Hays, K.E., Pfaffinger, J.M., and Ryznar, R. (2024). The interplay between gut microbiota, short-chain fatty acids, and implications for host health and disease. Gut microbes 16, 2393270.

38. Guo, T., Zhang, L., Xin, Y., Xu, Z., He, H., and Kong, J. (2017). Oxygen-inducible conversion of lactate to acetate in heterofermentative Lactobacillus brevis ATCC 367. Applied and environmental microbiology 83, e01659–01617.

39. Liu, X., Cooper, D.E., Cluntun, A.A., Warmoes, M.O., Zhao, S., Reid, M.A., Liu, J., Lund, P.J., Lopes, M., and Garcia, B.A. (2018). Acetate production from glucose and coupling to mitochondrial metabolism in mammals. Cell 175, 502–513. e513.

40. Argelaguet, R., Velten, B., Arnol, D., Dietrich, S., Zenz, T., Marioni, J.C., Buettner, F., Huber, W., and Stegle, O. (2018). Multi-Omics Factor Analysis—a framework for unsupervised integration of multi-omics data sets. Molecular systems biology 14, MSB178124.

41. Yamauchi, T., Oi, A., Kosakamoto, H., Akuzawa-Tokita, Y., Murakami, T., Mori, H., Miura, M., and Obata, F. (2020). Gut bacterial species distinctively impact host purine metabolites during aging in Drosophila. Iscience 23.

42. Newell, P.D., Preciado, L.M., and Murphy, C.G. (2022). A Functional Analysis of the Purine Salvage Pathway in Acetobacter fabarum. Journal of Bacteriology 204, e00041–00022. doi:10.1128/jb.00041-22.

43. Shi, L., and Tu, B.P. (2015). Acetyl-CoA and the regulation of metabolism: mechanisms and consequences. Current opinion in cell biology 33, 125–131.

44. Pietrocola, F., Galluzzi, L., Bravo-San Pedro, J.M., Madeo, F., and Kroemer, G. (2015). Acetyl coenzyme A: a central metabolite and second messenger. Cell metabolism 21, 805–821.

45. Brown, K., Thomson, C.A., Wacker, S., Drikic, M., Groves, R., Fan, V., Lewis, I.A., and McCoy, K.D. (2023). Microbiota alters the metabolome in an age-and sex-dependent manner in mice. Nature communications 14, 1348.

46. Hulme, H., Meikle, L.M., Strittmatter, N., Swales, J., Hamm, G., Brown, S.L., Milling, S., MacDonald, A.S., Goodwin, R.J., and Burchmore, R. (2022). Mapping the influence of the gut microbiota on small molecules across the microbiome gut brain axis. Journal of the American Society for Mass Spectrometry 33, 649–659.

47. Tarracchini, C., Lugli, G.A., Mancabelli, L., van Sinderen, D., Turroni, F., Ventura, M., and Milani, C. (2024). Exploring the vitamin biosynthesis landscape of the human gut microbiota. Msystems 9, e00929–00924.

48. Warneke, R., Herzberg, C., Klein, M., Elfmann, C., Dittmann, J., Feussner, K., Feussner, I., and Stülke, J. (2024). Coenzyme A biosynthesis in Bacillus subtilis: discovery of a novel precursor metabolite for salvage and its uptake system. Mbio 15, e01772–01724.

49. Gnainsky, Y., Zfanya, N., Elgart, M., Omri, E., Brandis, A., Mehlman, T., Itkin, M., Malitsky, S., Adamski, J., and Soen, Y. (2021). Systemic regulation of host energy and oogenesis by microbiome-derived mitochondrial coenzymes. Cell Reports 34.

50. Böttcher, J., Sibon, O.C., and El Aidy, S. (2025). Coenzyme A metabolism: a key driver of gut microbiota dynamics and metabolic profiles. FEMS Microbiology Reviews 49, fuaf051.

51. Hung, Y.-T., Tuan, S.-J., and Wong, A.C.-N. (2025). Systematic analysis of nutrient-microbiome interactions and their effects on host phenotypes in Drosophila. MBio 16, e02480–02425.

52. Barritt, S.A., DuBois-Coyne, S.E., and Dibble, C.C. (2024). Coenzyme A biosynthesis: mechanisms of regulation, function and disease. Nature Metabolism 6, 1008–1023.

53. Shichkin, V.P. (2025). Vitamin B5 and vitamin U review: justification of combined use for the treatment of mucosa-associated gastrointestinal pathologies. Frontiers in pharmacology 16, 1587627.

54. Wan, Z., Zheng, J., Zhu, Z., Sang, L., Zhu, J., Luo, S., Zhao, Y., Wang, R., Zhang, Y., and Hao, K. (2022). Intermediate role of gut microbiota in vitamin B nutrition and its influences on human health. Frontiers in nutrition 9, 1031502.

55. Cameron, D.D., Leake, J.R., and Read, D.J. (2006). Mutualistic mycorrhiza in orchids: evidence from plant–fungus carbon and nitrogen transfers in the green-leaved terrestrial orchid Goodyera repens. New Phytologist 171, 405–416.

56. Wang, W., Shi, J., Xie, Q., Jiang, Y., Yu, N., and Wang, E. (2017). Nutrient exchange and regulation in arbuscular mycorrhizal symbiosis. Molecular plant 10, 1147–1158.

57. Russell, C.W., Poliakov, A., Haribal, M., Jander, G., van Wijk, K.J., and Douglas, A.E. (2014). Matching the supply of bacterial nutrients to the nutritional demand of the animal host. Proceedings of the Royal Society of London B: Biological Sciences 281, 20141163.

58. Miao, T., Liu, Y., Qadiri, M., Dasseux, A., Asara, J.M., Hu, Y., Sun, X., Pliego-Alcántara, L.d.C., Dibble, C.C., and Perrimon, N. (2026). Renal coenzyme A (CoA) production from VB5 fuels stem cell proliferation and tumor growth. Nature Communications.

59. Grifoni, D., and Bellosta, P. (2015). Drosophila Myc: a master regulator of cellular performance. Biochimica et Biophysica Acta (BBA)-Gene Regulatory Mechanisms 1849, 570–581.

60. Kreuzaler, P., Inglese, P., Ghanate, A., Gjelaj, E., Wu, V., Panina, Y., Mendez-Lucas, A., MacLachlan, C., Patani, N., and Hubert, C.B. (2023). Vitamin B5 supports MYC oncogenic metabolism and tumor progression in breast cancer. Nature metabolism 5, 1870–1886.

61. Onuma, T., Yamauchi, T., Kosakamoto, H., Kadoguchi, H., Kuraishi, T., Murakami, T., Mori, H., Miura, M., and Obata, F. (2023). Recognition of commensal bacterial peptidoglycans defines Drosophila gut homeostasis and lifespan. PLoS Genetics 19, e1010709.

62. Wong, K.K.L., Liao, J.Z., and Verheyen, E.M. (2019). A positive feedback loop between Myc and aerobic glycolysis sustains tumor growth in a Drosophila tumor model. Elife 8, e46315.

63. Chintapalli, V.R., Wang, J., and Dow, J.A. (2007). Using FlyAtlas to identify better Drosophila melanogaster models of human disease. Nature genetics 39, 715–720.

64. Brummel, T., Ching, A., Seroude, L., Simon, A.F., and Benzer, S. (2004). Drosophila lifespan enhancement by exogenous bacteria. Proceedings of the National Academy of Sciences 101, 12974–12979.

65. White, K.M., Matthews, M.K., Hughes, R.C., Sommer, A.J., Griffitts, J.S., Newell, P.D., and Chaston, J.M. (2018). A metagenome-wide association study and arrayed mutant library confirm Acetobacter lipopolysaccharide genes are necessary for association with Drosophila melanogaster. G3: Genes, Genomes, Genetics 8, 1119–1127.

66. Choi, K.-H., and Schweizer, H.P. (2006). mini-Tn7 insertion in bacteria with single attTn7 sites: example Pseudomonas aeruginosa. Nature protocols 1, 153–161.

67. Shanks, R.M., Caiazza, N.C., Hinsa, S.M., Toutain, C.M., and O’Toole, G.A. (2006). *Saccharomyces cerevisiae*-based molecular tool kit for manipulation of genes from gram-negative bacteria. Appl Environ Microbiol 72, 5027–5036.

68. Zuber, S., Carruthers, F., Keel, C., Mattart, A., Blumer, C., Pessi, G., Gigot-Bonnefoy, C., Schnider-Keel, U., Heeb, S., and Reimmann, C. (2003). GacS sensor domains pertinent to the regulation of exoproduct formation and to the biocontrol potential of Pseudomonas fluorescens CHA0. Molecular plant-microbe interactions 16, 634–644.

69. Fossati, P., and Prencipe, L. (1982). Serum triglycerides determined colorimetrically with an enzyme that produces hydrogen peroxide. Clinical chemistry 28, 2077–2080.

70. Marcu, D., Sannino, D.R., Dornan, A.J., Ibrahim, R., Kapoor, A., Wood, M., and Dobson, A.J. (2025). Microbiota impact Drosophila ageing via Acetobacter, Tachykinin, and TkR99D. eLife 14.

71. Cole, J.J., Faydaci, B.A., McGuinness, D., Shaw, R., Maciewicz, R.A., Robertson, N.A., and Goodyear, C.S. (2021). Searchlight: automated bulk RNA-seq exploration and visualisation using dynamically generated R scripts. BMC bioinformatics 22, 411.

72. Frey, A.J., Feldman, D.R., Trefely, S., Worth, A.J., Basu, S.S., and Snyder, N.W. (2016). LC-quadrupole/Orbitrap high-resolution mass spectrometry enables stable isotope-resolved simultaneous quantification and 13C-isotopic labeling of acyl-coenzyme A thioesters: AJ Frey et al. Analytical and bioanalytical chemistry 408, 3651–3658.

73. Gloaguen, Y., Morton, F., Daly, R., Gurden, R., Rogers, S., Wandy, J., Wilson, D., Barrett, M., and Burgess, K. (2017). PiMP my metabolome: an integrated, web-based tool for LC-MS metabolomics data. Bioinformatics 33, 4007–4009.

74. Picart-Armada, S., Fernández-Albert, F., Vinaixa, M., Yanes, O., and Perera-Lluna, A. (2018). FELLA: an R package to enrich metabolomics data. BMC bioinformatics 19, 538.

75. Wang, H., Robinson, J.L., Kocabas, P., Gustafsson, J., Anton, M., Cholley, P.-E., Huang, S., Gobom, J., Svensson, T., and Uhlen, M. (2021). Genome-scale metabolic network reconstruction of model animals as a platform for translational research. Proceedings of the National Academy of Sciences 118, e2102344118.

76. Burgess, K., Creek, D., Dewsbury, P., Cook, K., and Barrett, M.P. (2011). Semi-targeted analysis of metabolites using capillary-flow ion chromatography coupled to high-resolution mass spectrometry. Rapid Communications in Mass Spectrometry 25, 3447–3452.

